# A new method for multi-ancestry polygenic prediction improves performance across diverse populations

**DOI:** 10.1101/2022.03.24.485519

**Authors:** Haoyu Zhang, Jianan Zhan, Jin Jin, Jingning Zhang, Wenxuan Lu, Ruzhang Zhao, Thomas U. Ahearn, Zhi Yu, Jared O’Connell, Yunxuan Jiang, Tony Chen, Dayne Okuhara, 23andMe Research Team, Montserrat Garcia-Closas, Xihong Lin, Bertram L. Koelsch, Nilanjan Chatterjee

## Abstract

Polygenic risk scores (PRS) increasingly predict complex traits, however, suboptimal performance in non-European populations raise concerns about clinical applications and health inequities. We developed CT-SLEB, a powerful and scalable method to calculate PRS using ancestry-specific GWAS summary statistics from multi-ancestry training samples, integrating clumping and thresholding, empirical Bayes and super learning. We evaluate CT-SLEB and nine-alternatives methods with large-scale simulated GWAS (∼19 million common variants) and datasets from 23andMe Inc., the Global Lipids Genetics Consortium, All of Us and UK Biobank involving 5.1 million individuals of diverse ancestry, with 1.18 million individuals from four non-European populations across thirteen complex traits. Results demonstrate that CT-SLEB significantly improves PRS performance in non-European populations compared to simple alternatives, with comparable or superior performance to a recent, computationally intensive method. Moreover, our simulation studies offer insights into sample size requirements and SNP density effects on multi-ancestry risk prediction.

## Introduction

Genome-wide association studies (GWAS) have identified tens of thousands of single nucleotide polymorphisms (SNPs) associated with complex traits and diseases^1^. Polygenetic risk scores (PRSs), summarize the combined effect of individual SNPs, offering potential to improve risk stratification for various diseases and conditions^2–7^. However, GWAS to date have primarily focused on populations predominately comprised of European (EUR) origin individuals^8^. Consequently, the PRSs generated from these studies tend to underperform in non-EUR populations, particularly in African (AFR) ancestry populations^9–12^. The limited representation of non-EUR populations in PRS research raises concerns that using current PRSs for clinical applications may exacerbate health inequities^13–16^.

In addition to the critical importance of addressing inequalities in representation of non-EUR population in genetic research, there is also an important need to develop statistical methods that leverage genetic data across populations to develop better performing PRS. Most existing PRS methods have been developed to analyze data from a single ancestry group^17–26^, and subsequently, their performance was primarily evaluated in EUR populations^3–6^. While the same methods can also be used to build PRS in non-EUR populations, the resulting PRS tend to have limited performance due to smaller training data sample sizes compared to EUR populations^9,14^. Some studies have conducted meta-analyses of GWAS across diverse populations to develop a multi-ancestry PRS^27–29^. While this approach may lead to a single PRS that performs more “equally” across diverse groups, it does not account for heterogeneity in linkage disequilibrium (LD) and effect sizes across populations, and is not designed to derive the best PRS possible for each population^30,31^.

Recent methods aim to develop more optimal PRS in non-EUR populations by combining available GWAS from the target population with “borrowed” information from larger GWAS in the EUR populations. One such study developed PRS in separate populations and then combined them by optimal weighting to maximize target population performance^32^. Other studies proposed Bayesian methods using multivariate priors for effect-size distribution to borrow information across populations^30,33,34^. Despite these developments, methods leveraging multi-ancestry datasets for PRS remain limited. Both theoretical and empirical studies have indicated that the optimal PRS building depends on multiple factors^17,35,36^, including sample size, heritability, effect-size distribution and LD, and thus exploration of alternative methods with complementary advantages are needed to build optimal PRS in any given setting. Moreover, and perhaps more importantly, evaluation of multi-ancestry methods for building improved PRS remains quite limited to date due to the lack of large GWAS for various non-EUR populations, especially of African origin, where risk prediction remains the most challenging.

In this paper, we propose CT-SLEB, a computationally simple and powerful method for generating PRSs using GWAS across diverse ancestry population. CT-SLEB is a model-free approach that combines multiple techniques, including a two-dimensional extension of the popular clumping and thresholding (CT) method^17,18^, a super-learning (SL) model for combining multiple PRS and an empirical-Bayes (EB) approach for effect-size estimation. We compare CT-SLEB’s performance with nine alternative methods using large-scale simulated GWAS across five ancestry groups. Additionally, we develop and validate population-specific PRS for thirteen complex traits using GWAS data from 23andMe, Inc., the Global Lipids Genetics Consortium (GLGC)^37^, All of Us (AoU) and UK Biobank (UKBB) across EUR (*N* ≈ 3.91 million), AFR (primarily African American (AA), *N* ≈ 265K), Latino (*N* ≈ 574K), East Asian (EAS, *N* ≈ 270K), and South Asian (SAS, *N* ≈ 77K). Both simulation studies and empirical data analyses indicate CT-SLEB as a scalable and powerful method for generating PRS for non-EUR populations. Further, our simulation studies and evaluation of various methods in large datasets provide insights into the future yield of multi-ancestry PRSs as GWAS in diverse populations continues to grow.

## Results

### Method overview

CT-SLEB is designed to generate multi-ancestry PRSs, incorporating large GWAS from EUR population and smaller GWAS from non-EUR populations. The method has three key steps (**Fig. 1, Extended Data Fig. 1**): 1. <u>Clumping and Thresholding (CT)</u> for selecting SNPs to be included in a PRS for the target population; 2<u>. Empirical-Bayes</u> <u>(EB)</u> method for SNPs coefficient estimation; 3. <u>Super-learning (SL)</u> model to combine a series of PRSs generated under different SNP selection thresholds. CT-SLEB requires three independent datasets: (1) GWAS summary statistics from training datasets across EUR and non-EUR populations; (2) a tuning dataset for the target population to find optimal model parameters; and (3) a validation dataset for the target population to report the final prediction performance.

**Figure 1:**
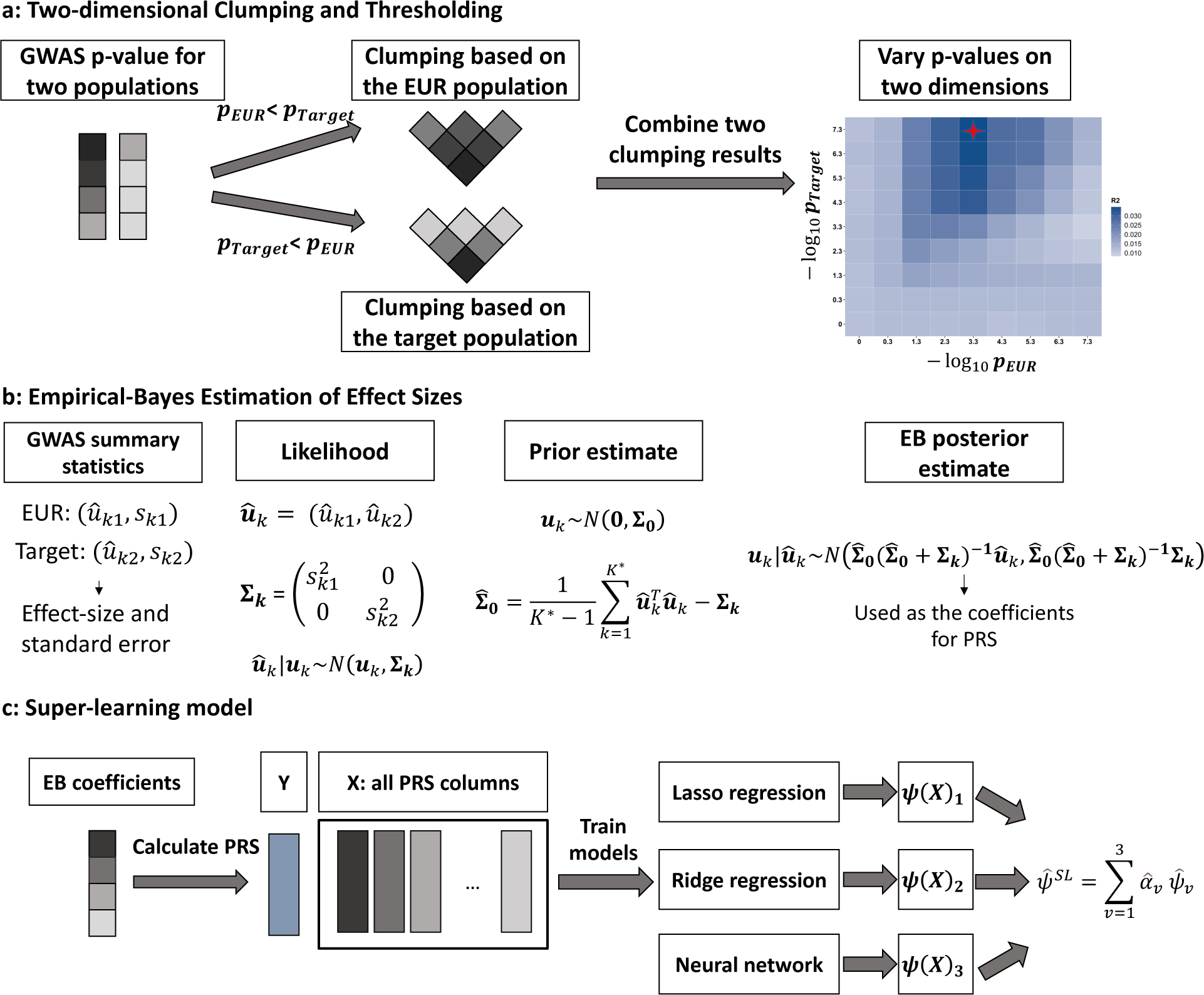
CT-SLEB Workflow. The method contains three key steps: 1. Two-dimensional clumping and thresholding method for selecting SNPs (Figure 1a); 2. Empirical-Bayes procedure for incorporating correlation in effect sizes of genetic variants across populations (Figure 1b); 3. Super-learning model for combing the PRSs derived from the first two steps under different tuning parameters (Figure 1c). The GWAS summary-statistics data are obtained from the training data. The tuning dataset is used to train the super-learning model. The final prediction performance is evaluated using an independent validation dataset.

#### Two-dimensional Clumping and Thresholding

In step one, CT-SLEB uses two-dimensional clumping and thresholding on GWAS summary-statistics data to incorporate SNPs with either shared effects across the EUR and the target populations or population-specific effects in the target population (**Fig. 1a**). Each SNP is assigned to one of two groups based on p-value from EUR and target populations: 1. SNPs with a p-value smaller in the EUR population; 2. SNPs with a p-value smaller in the target population or those which exist only in the target population. SNPs in the first group are ranked by EUR p-value (smallest to largest) and then clumped using LD estimates from EUR reference sample. SNPs in the second group are ranked by the target population p-value and clumped using LD estimates from target population reference sample. Clumped SNPs from both groups form a candidate set for the next step. In the thresholding step, p-value thresholds vary over a two-dimensional grid. Each dimension corresponds to the threshold for p-value from one population. At any threshold combination, a SNP may be included in the target population PRS if its p-value from either the EUR or the target population achieves the corresponding threshold.

#### Empirical-Bayes Estimation of Effect Sizes

Since SNP effect sizes are expected to be correlated across populations^38,39^, we propose an EB method to efficiently estimate effect sizes for SNPs to be included in the PRSs (**Fig. 1b**). Based on the selected SNP set from the CT step, we first estimate a “prior” covariance matrix of effect sizes between the EUR and target population. Then, we estimate each SNP’s effect size in the target population using the corresponding posterior mean, which weighs the effect-size estimate from each population based on the bias-variance trade-off (**Methods**).

#### Super Learning

Previous research has shown that combining PRSs under different p-value thresholds can efficiently increase prediction performance^20^. Therefore, we propose a super-learning model to predict the outcome using PRSs generated under different tuning parameters as training variables (**Fig. 1c**). The super-learning model is a linear combination of predictors based on multiple supervised learning algorithms^40–42^. The set of prediction algorithms can be self-designed or chosen from classical prediction algorithms. We use the R package SuperLearner version 2.0-26^43^ and choose Lasso^44^, ridge regression^45^, and neural networks^46^ as three different candidate models in the implementation. We train the super-learning model on the tuning dataset and evaluate the final PRS performance using the independent validation dataset.

#### Design of Simulation Studies

We conduct simulation studies comparing ten methods across five broad categories: 1. *single ancestry methods* using only target population data; 2. *EUR PR*S, generated using single ancestry methods on EUR only GWAS data; 3. *weighted PRS,* applying single ancestry PRS separately to the EUR and target population, and deriving an optimal linear combination of the two; 4. *Bayesian method,* assuming a multi-variate Bayesian framework for PRS construction; and 5. our proposed approach CT-SLEB. The single ancestry methods include CT^17,18^ and LDpred2^19,26^. EUR PRSs are generated using CT and LDpred2. Weighted PRS approaches include: CT-based, LDpred2-based, and PolyPred-S+^47^ . The last method linearly combines PRSs using EUR and target population PRS from SBayesR^21^, and EUR PRS from PolyFun-pred^48^, which integrates functional annotation information to identify causal variant across the genome and thus uses additional information that is not incorporated in the other methods compared. Bayesian methods include: 1. <u>XPASS method</u>^34^, assuming a multivariate normal distribution for effect-size and using the posterior mean of the target population to construct PRS. 2. <u>PRS-CSx</u>^30^, using a continuous shrinkage Bayesian framework to calculate the posterior mean of effect sizes for EUR and non-EUR populations, and subsequently deriving an optimal linear combination of all populations using a tuning dataset.

All methods use the target and EUR population training data to construct PRS for the target population. Additionally, CT-SLEB and PRS-CSx are evaluated using data from all five ancestries. For computational efficiency, most analyses are restricted to ∼2.0 million SNPs included in Hapmap3 (HM3)^49^, or the Multi-Ethnic Genotyping Arrays (MEGA)^50^ chips array, or both. However, the PolyPred-S+ and PRS-CSx methods are currently limited to ∼1.3 million HM3 SNPs in the provided software.

## Simulation Study Results

Results from simulation studies **(Fig. 2** and **Supplementary Fig. 1-5)** show that multi-ancestry methods generally lead to the most accurate PRSs in different settings. When the training data sample size for the target population is small (**Fig. 2a, Supplementary Fig. 1a, 2-5 a**), PRSs from single ancestry methods perform poorly compared to EUR-based PRS. Conversely, when the target population training sample size is large (**Fig. 1b, Supplementary Fig. 1b, 2-5 b-d**), PRSs from single ancestry methods can outperform EUR PRS. PRS generated from multi-ancestry methods can achieve substantial improvement in either setting.

**Figure 2:**
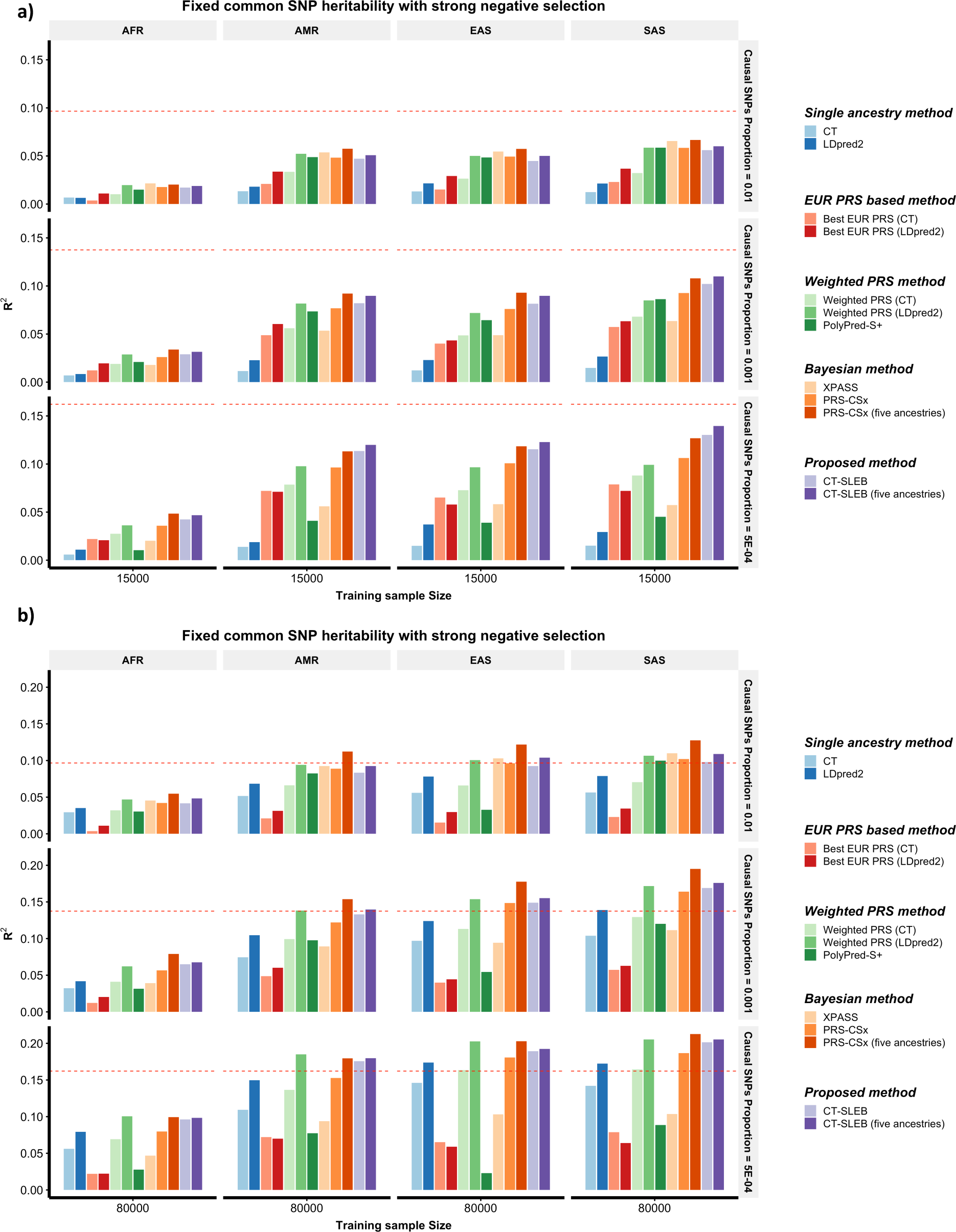
Simulation results of various PRSs methods in multi-ancestry settings. Each of the four non-EUR populations has a training sample size of **15,000 (**Figure 2a**) or 80,000 (**Figure 2b). For the EUR population, the size of the training sample is set at 100,000. The tuning dataset includes 10,000 samples per population. Prediction *R*\* values are reported based on an independent validation dataset with 10,000 subjects per population. Common SNP heritability is assumed to be 0.4 across all populations, and effect-size correlation is assumed to be 0.8 across all pairs of populations. The proportion of causal SNPs varies across 0.01 (top panel), 0.001 (medium panel), 5 × 10^−4^ (bottom panel), and effect sizes for causal variants are assumed to be related to allele frequency under **<u>a strong negative selection model</u>**. Data are generated based on ∼19 million common SNPs across the five populations, but analyses are restricted to ∼2.0 million SNPs that are used on Hapmap3 + Multi-Ethnic Genotyping Arrays chip. PolyPred-S+ and PRS-CSx analyses are further restricted to ∼1.3 million HM3 SNPs. All approaches are trained using data from the EUR and the target population. CT-SLEB and PRS-CSx are also evaluated using five ancestries data as training data. The red dashed line shows the prediction performance of EUR PRS generated using the single ancestry method (best of CT or LDpred2) in the EUR population.

When using only EUR and target population data, both CT-SLEB and PRS-CSx can lead to improvements over other candidate methods in most settings. When the target population sample size is large, weighted LDpred2 performs comparably to CT-SLEB and PRS-CSx. Between CT-SLEB and PRS-CSx, neither is uniformly superior across all scenarios. With a smaller target population sample size (N = 15K), PRS-CSx often outperforms CT-SLEB at the highest degree of polygenicity (*p_causal_* = 0.01), while CT-SLEB excels at the lowest polygenicity (*p_causal_* = 5 × 10^−4^). The difference between the two methods narrows with larger sample sizes (N=45K-100K). When using data from all five ancestries simultaneously, CT-SLEB and PRS-CSx improves by 6.8% and 23.7% on average, respectively, compared to only using EUR and target population data. PRS-CSx outperforms CT-SLEB in many settings (**Fig. 1b, Supplementary Fig. 1a-b, 2-5 b-d)**. Under different simulation settings, the number of SNPs used by CT-SLEB ranged from 549K to 933K, while PRS-CSx retained all HM3 SNPs (**Supplementary Table 1**).

Comparing runtime for constructing AFR PRS on chromosome 22 data (**Methods, Supplementary Table 2**), CT-SLEB is on average almost 25 times faster than PRS-CSx (4.35 vs. 109.11 mins) in two ancestries analyses, and 91 times faster than that of PRS-CSx in five ancestries setting (4.62 vs. 420.96 mins) using a single core with Intel E5-26840v4 CPU. Sensitivity analyses evaluate the required tuning and validation sample size for CT-SLEB (**Extended Data Fig. 2**), showing increased prediction performance with larger sizes. Meanwhile, CT-SLEB’s performance remains robust when the tuning and validation sample size is around 2000. Additional sensitivity analyses compare CT-SLEB and PRS-CSx performance when both methods use HM3 SNPs. CT-SLEB maintains an advantage in low polygenicity setting with training sample size of 15K or 45K (**Supplementary Fig. 6a-b**). However, with training sample size above 80K, both methods show similar performance in low polygenicity setting (**Supplementary Fig. 6c-d**).

Unequal predictive performance of PRS across populations presents an ethnical barrier for implementing this technology in healthcare. We examine the required training GWAS sample size for minority populations to bridge the performance gap compared to the EUR population. Results indicate that when effect sizes for shared causal SNPs are similar across populations (genetic correlation=0.8), the gap is mostly eliminated for all populations except AFR when the sample size reaches between 45% to 80% of the EUR population (**Fig. 3, Supplementary Fig. 7**). However, for the AFR population, sample size requirements can vary dramatically depending on the genetic architecture of traits. When we assume equal common SNP heritability for AFR and other populations, the AFR sample size requirement seems dauntingly large because of smaller per-SNP heritability (**Fig. 3a-b, Supplementary Fig. 7a-b**). If per-SNP heritability remains the same across populations, but heritability varies proportionately to the number of common variants, the AFR sample size requirement aligns with those of other minority populations (**Fig. 3c-d, Supplementary Fig. 7c)**.

**Figure 3:**
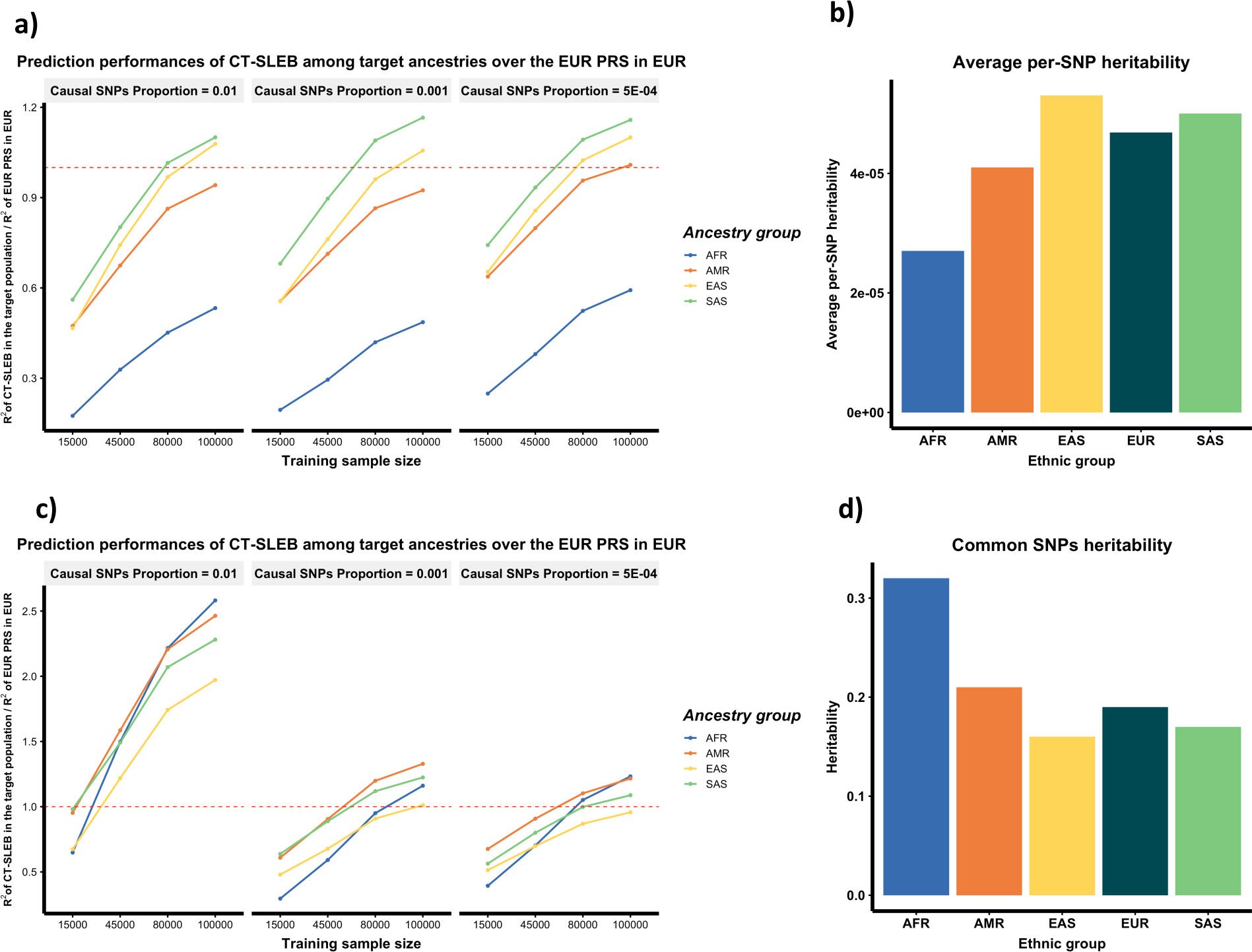
Comparison of CT-SLEB PRS across different ancestries to single ancestry EUR PRS in the EUR population. The training sample size for each of the four non-EUR populations is 15,000, 45000, 80,000, or 100,000. The training sample size for the EUR population is fixed at 100,000, and PRS performance is evaluated using single ancestry CT or LDpred2, whichever performs the best in each setting. Two different genetic architectures are considered: either the common SNP heritability is fixed (at 0.4) (**Figure 3a and 3b**) or the per-SNP heritability is fixed (**Figure 3c and 3d**) across the five populations (**Figure 3c and 3d**). The effect-size correlation is assumed to be 0.8 across all pairs of populations. The effect sizes for causal variants are assumed to be related to allele frequency under a strong negative selection model. CT-SLEB uses the summary statistics from all five ancestries.

CT-SLEB has a major advantage over PRS-CSx in computational scalability, allowing it to handle a much larger number of SNPs. We use CT-SLEB to study the effect of SNP density on PRS performance by considering three SNP sets for PRS building: (1) ∼1.3 million SNPs in HM3^49^ (2) ∼2.0 million SNPs including all HM3 SNPs and additional SNPs in the MEGA array (3) All ∼19 million common SNPs included in the 1000 Genomes Project (Phase 3)^51^ which were used to generate the traits in our simulation studies. We observe that PRS performance in various US minority populations can be substantially enhanced by including SNPs in denser panels. This benefit due to denser panels is more enhanced when the target population sample size is larger and in settings with fewer causal SNPs (**Fig. 4 and Supplementary Fig. 8**).

**Figure 4:**
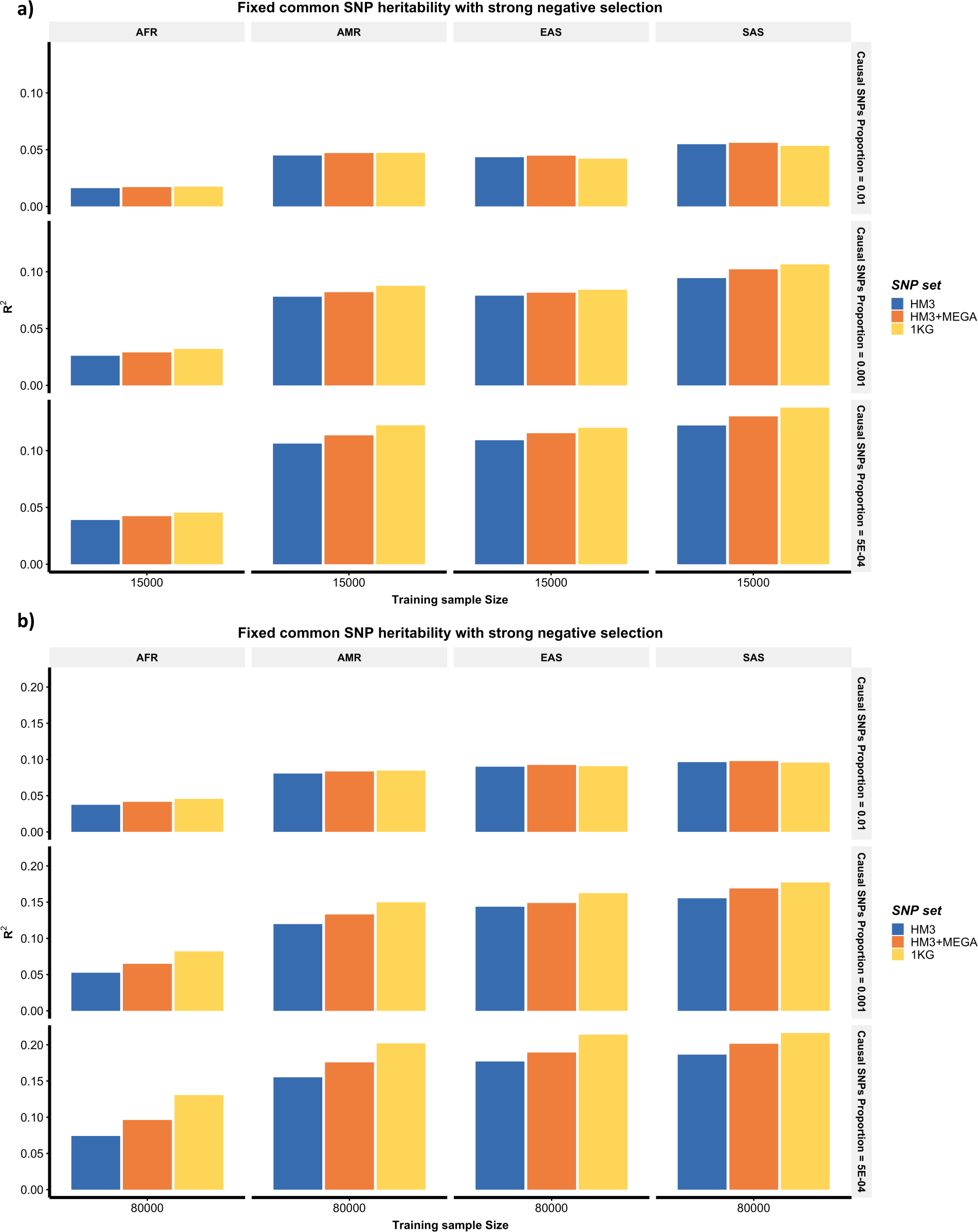
Prediction performance of CT-SLEB PRS under varying SNP densities. The analysis of each simulated data based on ∼19 million SNPs is restricted to three different SNP sets: Hapmap3 (∼1.3 million SNPs), Hapmap3 + Multi-Ethnic Genotyping Arrays (∼2.0 million SNPs), 1000 Genomes Project (∼19 million SNPs). The training sample size for each of the four non-EUR populations is 15,000 (Figure 4a) or 80,000 (Figure 4b). The training sample size for the EUR population is fixed at 100,000. Prediction *R*\* values are reported based on independent validation dataset with 10,000 subjects per population. <u>Common SNP heritability is assumed to be 0.</u>4 across all populations, and effect-size correlation is assumed to be <u>0.8</u> across all pairs of populations. The proportion of causal SNPs proportion vary across 0.01 (top panel), 0.001 (medium panel), 5 × 10^−4^ (bottom panel), and effect sizes for causal variants are assumed to be related to allele frequency under <u>a strong negative selection model</u>.

### 23andMe data analysis results

We develop and validate population-specific PRS for seven complex traits using GWAS data from 23andMe, Inc. (**Methods, Supplementary Table 3-4, Supplementary Data**). We conduct GWAS using a training dataset for each population adjusting for principal components (PC) 1-5, sex and age (**Methods**). The Manhattan plots and QQ plots for GWAS are in **Supplementary Fig. 9-15**, and no inflation is observed (**Supplementary Table 5**). We estimate heritability for the seven traits in the EUR population using LD-score regression^52^ (**Supplementary Table 6, Methods**).

Results for heart metabolic disease burden and height (**Fig. 5, Supplementary Table 7**) show a similar pattern as our simulation studies. The PRS-CSx using five ancestries generally leads to the best performing PRS across different populations. With only EUR and target population data, both CT-SLEB and PRS-CSx perform well across different populations. The relative gain is often large, especially for the AA population, compared to the best performing EUR or single ancestry PRS. Weighted methods don’t perform well for the AA population, but substantially improve performance compared to each component PRS (EUR and single ancestry) for other populations. PolyPred-S+ has comparable performance as PRS-CSx and CT-SLEB on EAS and SAS population, but is notably worse on AA population. We also observe that even with the best performing method and large sample, a significant gap remains for PRS performance in non-EUR populations compared to the EUR population (**Fig. 5**).

**Figure 5:**
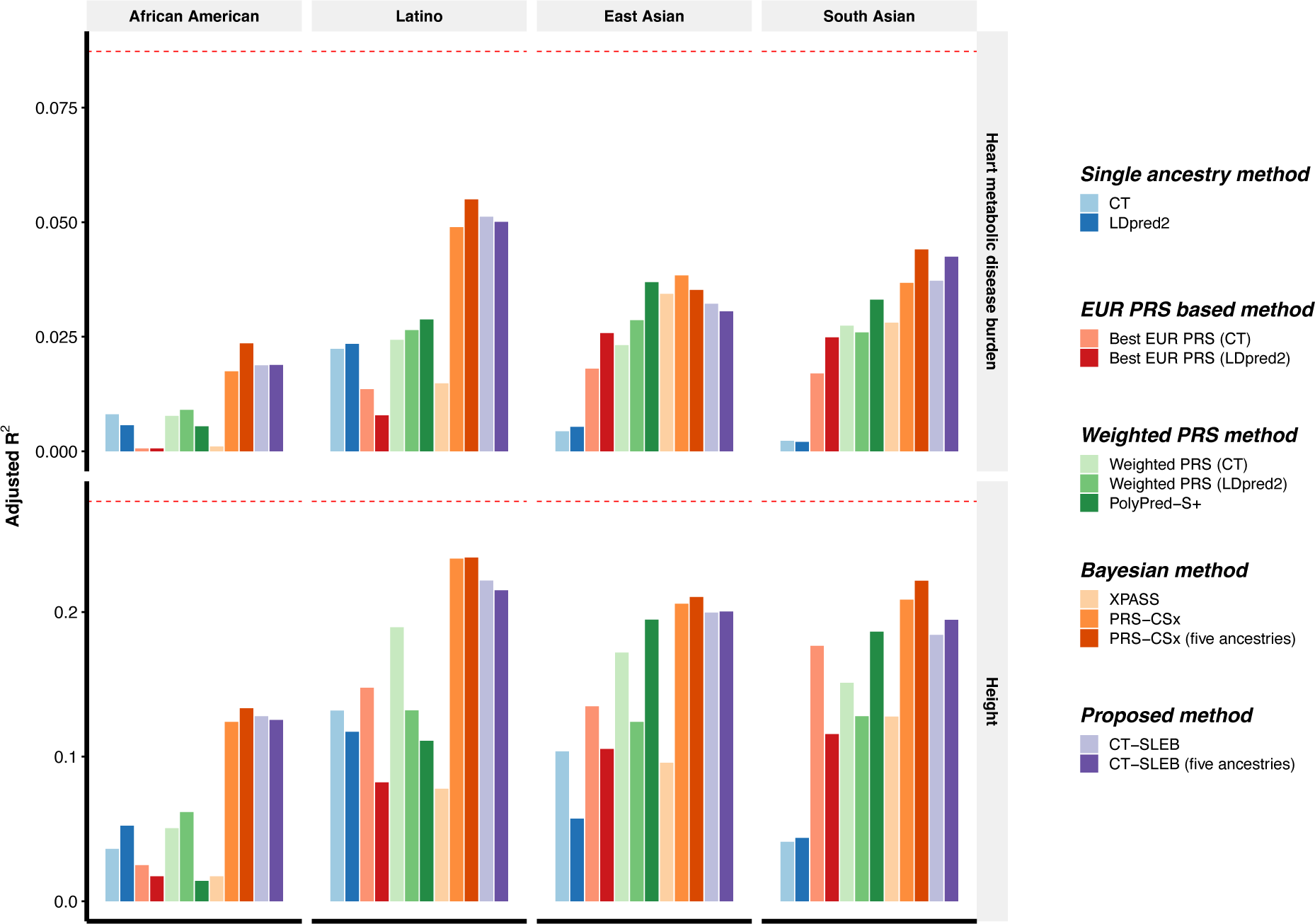
Prediction accuracy of PRSs for heart metabolic disease burden and height in 23andMe, Inc. datasets. The total sample size for heart metabolic disease burden and height is, respectively, 2.46 million and 2.93 million for European, 131K and 141K for African American, 375K and 509K for Latino, 110K and 121K for East Asian, and 29K and 32K for South Asian. The dataset is randomly split into 70%, 20%, 10% for training, tuning and validation dataset, respectively. The adjusted R^2^ values are reported based on the performance of the PRS in the validation dataset, accounting for PC1-5, sex and age. The red dashed line represents the prediction performance of EUR PRS generated using single ancestry method (best of CT or LDpred2) in the EUR population. Analyses are restricted to ∼2.0 million SNPs that are included in Hapmap3, or the Multi-Ethnic Genotyping Arrays chips array or both. PolyPred-S+ and PRS-CSx analyses are further restricted to ∼1.3 million HM3 SNPs. All approaches are trained using data from the EUR and the target population. CT-SLEB and PRS-CSx are also evaluated using training data from five ancestries. From top to bottom panel, two continuous traits are displayed in the following order: 1. heart metabolic disease burden; 2. height.

We observe similar trends in 23andMe data analysis for five binary traits: any cardiovascular disease (any CVD), depression, migraine diagnosis, morning person, and sing back musical note (SBMN) (**Fig. 6, Supplementary Table 7**). In most settings, CT-SLEB, PRS-CSx and PolyPred-S+ often produce superior PRS, improving upon best EUR or single ancestry PRS. For CVD, which is the clinically most relevant trait for risk prediction and preventive intervention, CT-SLEB outperforms PRS-CSx and PolyPred-S+ by a notable margin except for the EAS population. For the AFR population, particularly underrepresented in genetic research, CT-SLEB outperforms PRS-CSx and PolyPred-S+ by a notable margin for several traits (e.g., CVD and morning person). Conversely, PRS-CSx and PolyPred-S+ significantly outperforms CT-SLEB for predicting migraine diagnosis and SBMN in the SAS population. Despite the best performing methods and large GWAS in non-EUR populations, a major gap remains for PRS performance compared to the EUR population.

**Figure 6:**
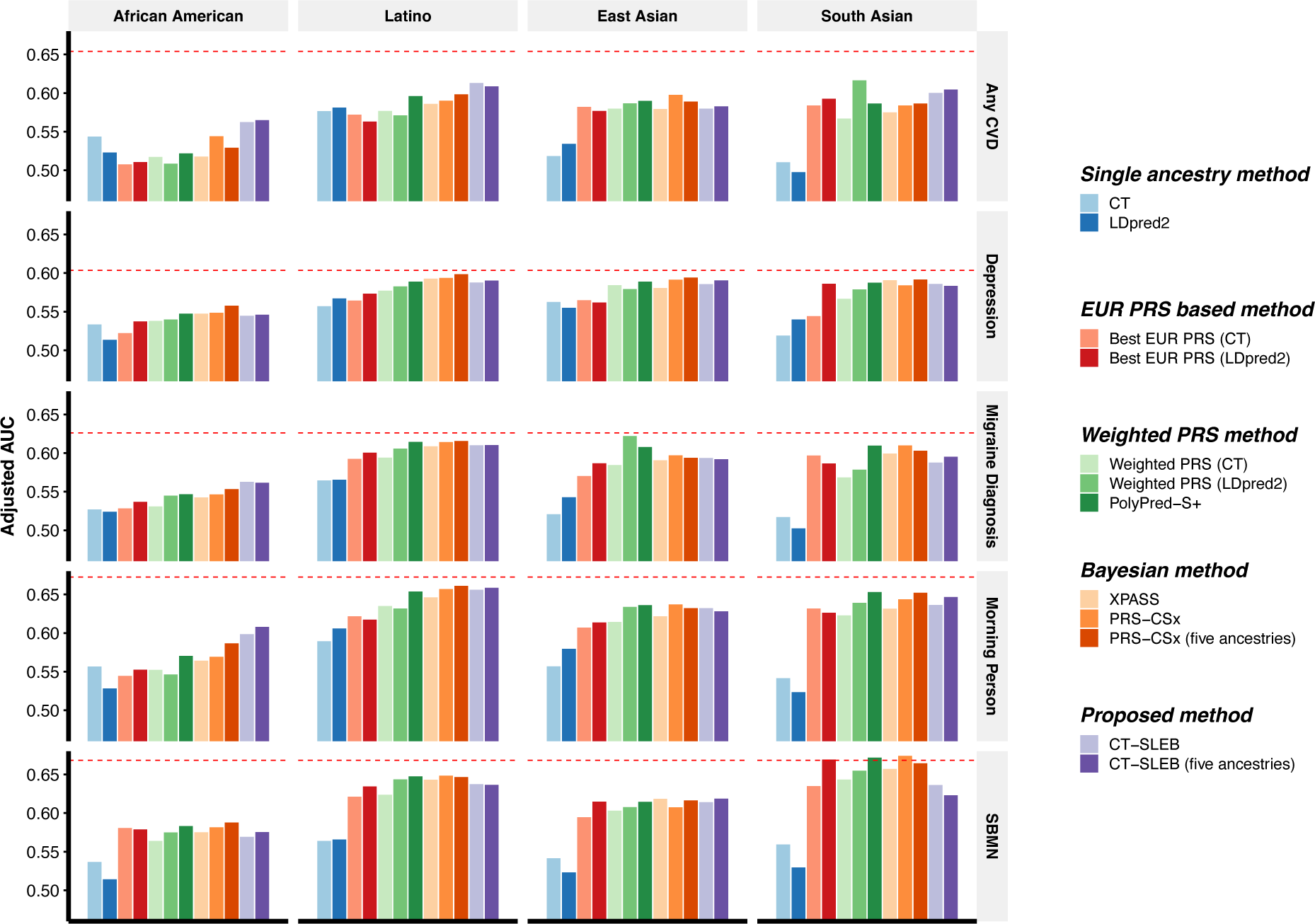
Prediction accuracy of five binary traits in 23andMe, Inc datasets. The data are from five populations: European (averaged N ≈ 2.37 million), African American (averaged N ≈ 109K), Latino (averaged N ≈ 401K), East Asian (averaged N≈ 86K), South Asian (averaged N≈ 24K). The datasets are randomly split into 70%, 20%, 10% for training, tuning and validation dataset, respectively. The adjusted AUC values are reported based on the validation dataset accounting for PC1-5, sex and age. The red dashed line represents the prediction performance of EUR PRS generated using single ancestry method (best of CT or LDpred2) in the EUR population. Analyses are restricted to ∼2.0 million SNPs that are included in Hapmap3, or the Multi-Ethnic Genotyping Arrays chips array or both. PolyPred-S+ and PRS-CSx analyses are further restricted to ∼1.3 million HM3 SNPs as implemented in the provided software. All approaches are trained using data from the EUR and the target population. CT-SLEB and PRS-CSx are also evaluated using training data from five ancestries. From top to bottom panel, five binary traits are displayed in the following order: 1.any cardiovascular disease (any CVD); 2. depression; 3. migraine diagnosis; 4. sing back musical note (SBMN); 5. morning person.

### GLGC & AoU analysis results with UKBB as validation dataset

We develop and validate population-specific PRS using GWAS summary data for four traits from GLGC: high-density lipoprotein cholesterol (HDL), low-density lipoprotein cholesterol (LDL), log triglycerides (logTG) and total cholesterol (TC), and two traits from AoU: height and body mass index (BMI) (**Methods, Supplementary Table 4, 8**). We evaluate the methods using individual level data from UKBB (**Supplementary Table 9**). The Manhattan plots and QQ plots for GWAS are shown in **Supplementary Fig. 16-21**, with no inflation observed given the genomic inflation factor for most ancestries, except height for Latino population in AoU (*λ*_1000_ = 1.04, **Supplementary Table 5**). We estimate heritability for the four traits in the EUR population using LD-score regression^52^ (**Supplementary Table 6, Methods**).

In GLGC data analyses, CT-SLEB, PRS-CSx and PolyPred-S+ outperform the other approaches (**Fig. 7**). CT-SLEB demonstrates superior performance in the AFR population, improving adjusted R^2^ for log-TG and LDL 140% and 66.4%, respectively, compared to PRS-CSx. This highlights the advantage of model-free approach CT-SLEB in handling biomarker traits, with some SNPs having unusually large effects. Conversely, PRS-CSx outperforms CT-SLEB in the EAS population. For log-TG and LDL, the adjusted R^2^ ratio between CT-SLEB and PRS-CSx is only 89.8% and 52.9%, respectively. Interestingly, for LDL and TC, PRS generated by CT-SLEB and five ancestries PRS-CSx demonstrate superior performance in the AFR population compared to EUR-PRSs in the European population (**Fig. 7**). Finally, in AoU data analyses, CT-SLEB performs better in predicting BMI, while PRS-CSx predicts height more accurately (**Fig. 8**). These results highlight the importance of generating PRS using multiple alternative methods in multi-ancestry settings.

**Figure 7:**
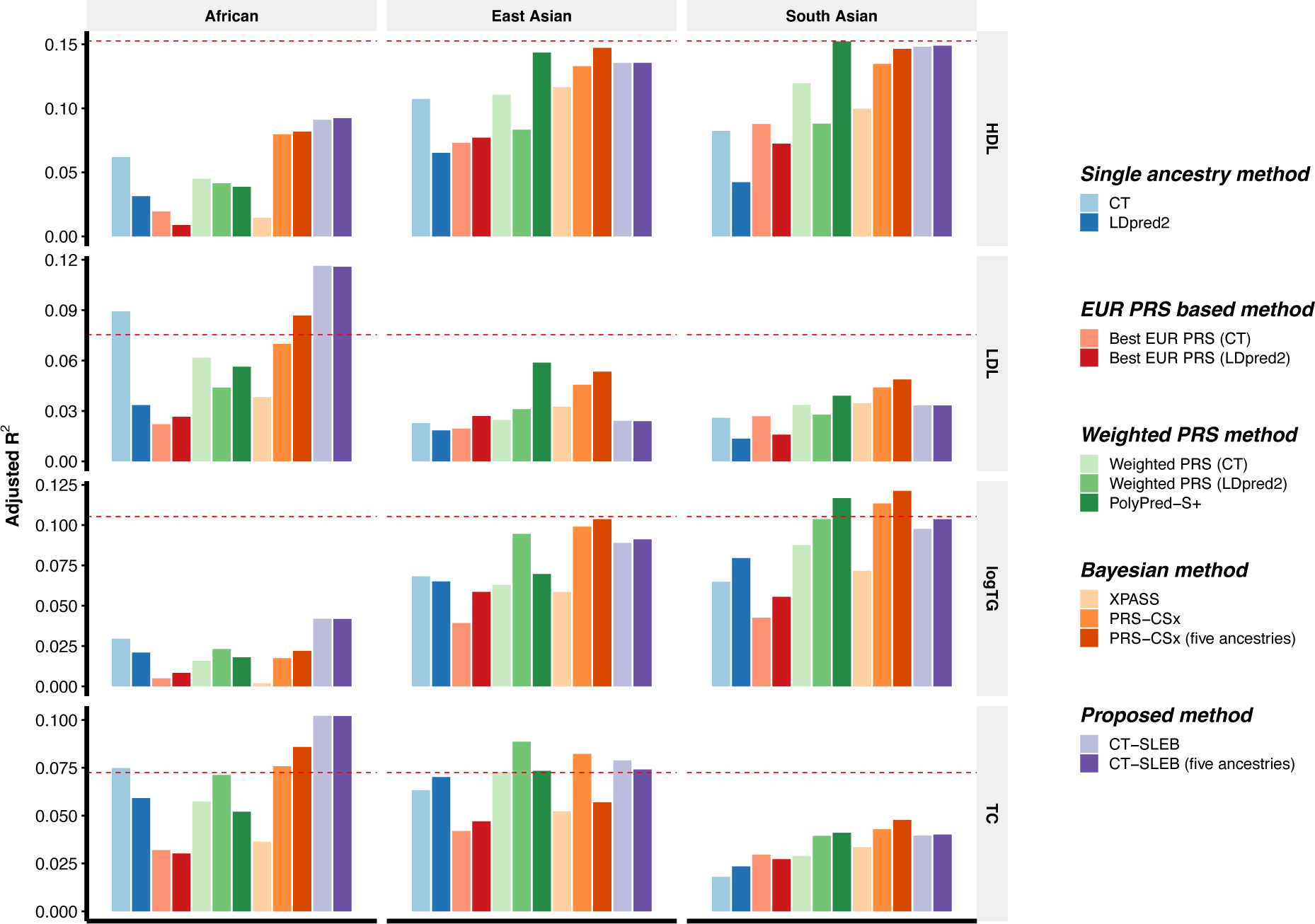
Prediction accuracy of four blood lipids traits from the Global Lipids Genetics Consortium. We used the GWAS summary statistics from five populations as the training data: European (N ≈ 931K), African (primarily African American, N ≈ 93K), Latino (N≈ 50K), East Asian (N ≈ 146*K*), South Asian (N ≈ 34*K*). The tuning and validation dataset are from UK Biobank data with three different ancestries African (N = 9,042), East Asian (N = 2,009), South Asian (N = 10,615). The tuning and validation are split with half and half. The adjusted R^2^ values are reported based on the performance of the PRS in the validation dataset, while accounting for PC1-10, sex and age. The red dashed line represents the prediction performance of EUR PRS generated using single ancestry method (best of CT or LDpred2) in the EUR population. Analyses are restricted to ∼2.0 million SNPs that are included in Hapmap3, or the Multi-Ethnic Genotyping Arrays chips array or both. PolyPred-S+ and PRS-CSx analyses are further restricted to ∼1.3 million HM3 SNPs as has been implemented in the provided software. All approaches are trained using data from the EUR and the target population. CT-SLEB and PRS-CSx are also evaluated using training data from five ancestries. From top to bottom panel, four traits are displayed in the following order: 1. high-density lipoprotein cholesterol (HDL), 2. low-density lipoprotein cholesterol (LDL), 3. log triglycerides (logTG) and 4. total cholesterol (TC).

**Figure 8:**
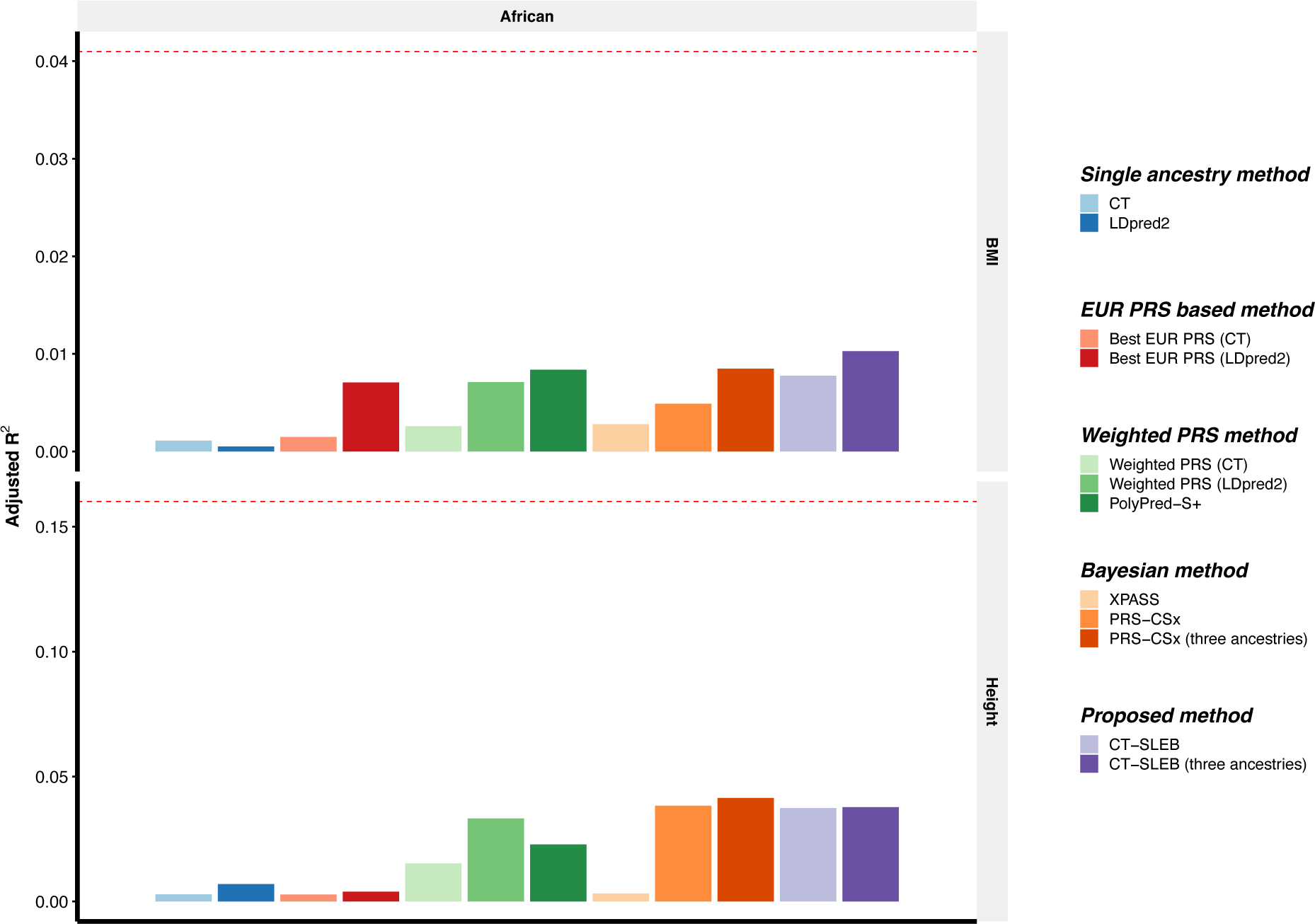
Prediction accuracy of two traits from All of Us dataset. We used the GWAS summary statistics from three populations as the training data: European (N ≈ 48K), African (N ≈ 22K), Latino (averaged N ≈ 15K). The tuning and validation dataset are from UK Biobank data with African (N = 9,042). The tuning and validation are split with half and half. The adjusted R^2^ values are reported based on the performance of the PRS in the validation dataset, while accounting for PC1-10, sex and age. The red dashed line represents the prediction performance of EUR PRS generated using single ancestry method (best of CT or LDpred2) in the European population. Analyses are restricted to around 800K SNPs that are genotyped in All of Us dataset for different ancestries. All approaches are trained using data from the European and African population. CT-SLEB and PRS-CSx are further evaluated using training data from three ancestries: African, European and Latino. From top to bottom panel, two traits are displayed in the following order: 1. body mass index (BMI), and 2. height.

To directly compare CT-SLEB and PRS-CSx, we report the prediction performance R^2^ (continuous traits) or logit-scale variance (converted from Area under the ROC curve (AUC) for binary traits, **Supplementary Note**) between CT-SLEB and PRS-CSx, averaging over the thirteen traits within each ancestry in 23andMe, GLGC and AoU data analyses (**Supplementary Table 10**). When only EUR and target population data are used for PRS construction, the averaged performance ratio between CT-SLEB and PRS-CSx is 144%, 88.5%, 103% and 92.1% for AFR (primarily AA), EAS, Latino, SAS, respectively. When data from five ancestries or three ancestries (AoU) are used for PRS construction, the averaged performance ratio is 143%, 91.7%, 94.5%, 89.6% for AFR (primarily AA), EAS, Latino, SAS, respectively.

## Discussion

In summary, we propose CT-SLEB as a powerful and computationally scalable method to generate optimal PRS across ancestrally distinct groups using GWAS across diverse populations. We compare CT-SLEB’s performance with various simple and complex methods, in large-scale simulation studies and datasets. Results show that no single method is uniformly best across all scenarios, and it’s important to generate PRS using alternative methods across multi-ancestries. Encouragingly, CT-SLEB leads to marked improvement in PRS performance compared to alternatives for AFR origin populations, where polygenic prediction has been most challenging. Computationally, CT-SLEB is an order of magnitude faster than a recently proposed Bayesian method, PRS-CSx^30^, and can more easily handle much larger SNP contents and additional populations.

A unique contribution of our study is the evaluation of a variety of PRS methodologies in the unprecedented large and diverse datasets from the 23andMe, Inc., GLGC, AoU and UKBB GWAS. Our findings offer crucial insights into the future yield of emerging large multi-ancestry GWAS. Adult height is often used as a model trait to explore the genetic architecture of complex traits and the potential for polygenic prediction. The standard CT method, when trained in ∼2 million EUR individuals from 23andMe data, leads to a PRS with a prediction *R*\* of approximately 0.276. Using LD-score regression, we estimated GWAS heritability of height using the 23andMe data to be 0.395, indicating that the PRS has achieved about 69.8% (0.276/0.395) of its maximum potential in the 23andMe EUR population. However, even with the best method and large 23andMe GWAS sample (N_Latino_∼350K and N_AFR_ ∼100K), the highest prediction accuracy of height PRS for non-EUR populations is substantially lower compared to that of the EUR population (Relative *R*\*∼0.48 (0.133/0.276) for AFR, 0.76 (0.210/0.276) for EAS, ∼ 0.86 (0.237/0.276) for Latino, and ∼ 0.80 (0.222/0.276) for SAS compared to that of EUR. The average relative R^2^ (continuous traits) or logit-scale variance (binary traits) for AFR, EAS, Latino, and SAS across the thirteen evaluated traits is 0.50, 0.76, 0.77, and 0.79, respectively (**Supplementary Table 11**).

We observe similar patterns for other traits, including disease outcomes for which risk prediction is of most interest. For CVD, for example, the CT method, trained in a sample of ∼700K cases and ∼1.3 million controls from the EUR population, produces a PRS with a prediction accuracy of AUC 0.65. For other populations with considerable but smaller sample sizes than the EUR population (N_case_/N_control =_32K/66K for AA and N_case_/N_control =_84K/270K for Latino populations), the AUCs for the best performing PRSs are close to 60% or lower. Further, sample size isn’t the only factor for differential PRS performances across populations. For example, the performance of the best CVD-PRS for Latino and South Asian populations are similar, despite a much smaller sample size for the latter population. Collectively, these findings and additional simulation study results, indicate that bridging PRS performance gap across populations requires greater parity in GWAS sample sizes.

Our simulation studies and data analyses show that no single PRS method is uniformly most powerful in all settings. In the analysis of GLGC data, for example, CT-SLEB greatly improves PRS performance compared to PRS-CSx for the AFR population, but the opposite is true for the EAS population. The optimal method for generating PRS depends on the underlying multivariate effect-size distribution of the traits across different populations. While Bayesian methods, in principle, can generate optimal PRS under correct specification of underlying effect-size distribution^19,26,30,34^, modeling this distribution in multi-ancestry settings can be challenging. In the analysis of lipid traits in GLGC, for example, we found the Bayesian methods can’t account well for existence of large-effect SNPs in the AFR population due to inadequacy of the underlying model for the effect-size distribution. Conversely, the CT method and their extensions, while do not require strong modeling assumptions about effect-size distribution, cannot optimally incorporate LD among SNPs. Our analysis reveals the advantages of alternative methods in different settings by comparing results across various complex traits with distinct architecture in terms of heritability, polygenicity, and number of clusters of distinct effect-sizes^53,54^. We thus advocate that in future applications, researchers consider generating and evaluating a variety of PRS obtained from complementary methods. As different PRS may contain some orthogonal information, the best strategy could be to combine them using a final super learning step, rather than choosing one best PRS.

Our study has several limitations. While 23andMe datasets have extremely large sample size, the power of genetic risk prediction is likely to be blunted in this population, compared to other settings, due to higher environmental heterogeneity. For example, a recent study^55^ reported achieving prediction R^2^ for height of ∼41% for the EUR individuals within the UKBB using a PRS developed using ∼1.1 million individuals from the UKBB (N=400K) and 23andMe (N=700K). In comparison, the PRS prediction R^2^ for height we achieve within 23andMe EUR population is only ∼30% despite doubling the sample size of the training dataset. However, we also note that heritability estimate in 23andMe (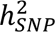 = 0.395) is substantially smaller than those previously reported^56,57^ based on the UKBB (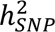 ∼0.5 − 0.7). Comparing the results across the two studies using prediction R^2^ relative to the underlying heritability of the respective populations, we observe a significant gain in performance due to the increased sample size of the current study. Thus, while caution is needed to extrapolate 23andMe study results to other populations, the relative performance of PRSs across different methods and different ancestry groups within this population is likely generalizable to other settings.

While we have conducted large-scale simulation across various scenarios, the simulated genotype data from HapGen2 may not fully reflect the levels of differentiation within and across ancestries due to limited haplotype data within 1000 Genomes Project. Furthermore, our proposed method, as well as many existing methods, primarily focus on generating PRS across ancestrally distinct populations. However, highly admixed populations like the AFR and Latino origin in the US could benefit from methods that explicitly account for individual-level estimates of admixture proportions^58^. Additionally, our method assumes that individual-level data is available for model tuning and validation, but this is not a fundamental limitation as summary-statistics-based methods^59,60^ could also be used in these steps. Moreover, while we observe a consistent increase of five ancestries CT-SLEB over two ancestries CT-SLEB in simulations, real data analyses don’t show the same consistent pattern (**Fig. 5-7**), potentially due to the complexity of underlying effect-sizes across different ancestries compared to simulations.

In conclusion, we have proposed a novel and computationally scalable method for generating powerful PRS using data from GWAS studies in diverse populations. Further, our simulation studies and data analysis across multiple traits involving large 23andMe Inc., GLGC, AoU and UKBB studies provide unique insight into the potential outcomes of future GWAS in diverse populations for years to come.

## Acknowledgements

We would like to thank the research participants and employees of 23andMe, Inc for making this work possible. We want to thank Liz Noblin, Melissa J. Francis and Emily Voeglein for helping with the research collaboration agreement with Harvard T.H. Chan School of Public Health, Johns Hopkins Bloomberg School of Public Health and 23andMe, Inc. The analysis utilized the high-performance computation Biowulf cluster at National Institutes of Health, USA, Faculty of Arts and Sciences Research Computing Cluster at Harvard University, and the Joint High Performance Computing Exchange at Johns Hopkins Bloomberg School of Public Health. The UK Biobank data was obtained under the UK Biobank resource application 17712. This work was funded by NIH grants: K99 CA256513-01 (H. Z.), R00 HG012223 (J.J.), NHLBI 5T32HL007604-37 (Z.Y.), R35-CA197449, U19-CA203654, R01-HL163560, U01-HG009088 U01-HG012064 (X. L.), R01 HG010480-01 (N. C. and J. Z.) and U01HG011724 (N. C.). The AoU Research Program is supported by the National Institutes of Health, Office of the Director: Regional Medical Centers: 1 OT2 OD026549; 1 OT2 OD026554; 1 OT2 OD026557; 1 OT2 OD026556; 1 OT2 OD026550; 1 OT2 OD 026552; 1 OT2 OD026553; 1 OT2 OD026548; 1 OT2 OD026551; 1 OT2 OD026555; IAA #: AOD 16037; Federally Qualified Health Centers: HHSN 263201600085U; Data and Research Center: 5 U2C OD023196; Biobank: 1 U24 OD023121; The Participant Center: U24 OD023176; Participant Technology Systems Center: 1 U24 OD023163; Communications and Engagement: 3 OT2 OD023205; 3 OT2 OD023206; and Community Partners: 1 OT2 OD025277; 3 OT2 OD025315; 1 OT2 OD025337; 1 OT2 OD025276. In addition, the AoU Research Program would not be possible without the partnership of its participants.

## Author Contributions

H.Z. and N.C. conceived the project. H.Z., J.Z. (Jianan Zhan), J.J., Jn.Z. (Jingning Zhang), W.L. and R.Z. carried out all data analyses with supervision from N.C.; J.Z., J.O.C., Y.J. ran GWAS for training data from 23andMe Inc. with the supervision from B.L.K.; R.Z. ran GWAS for training data from AoU with the supervision from N.C. and H.Z.; H.Z., T.C., and D.O. developed the software and online resources for data sharing; H.Z., J.Z., J.J., Jn.Z., W.L., R.Z. and N.C. drafted the manuscript, and X.L., M.G.C. and T.U.A. provided comments. All co-authors reviewed and approved the final version of the manuscript.

---

**Extended Data Figure 1:**
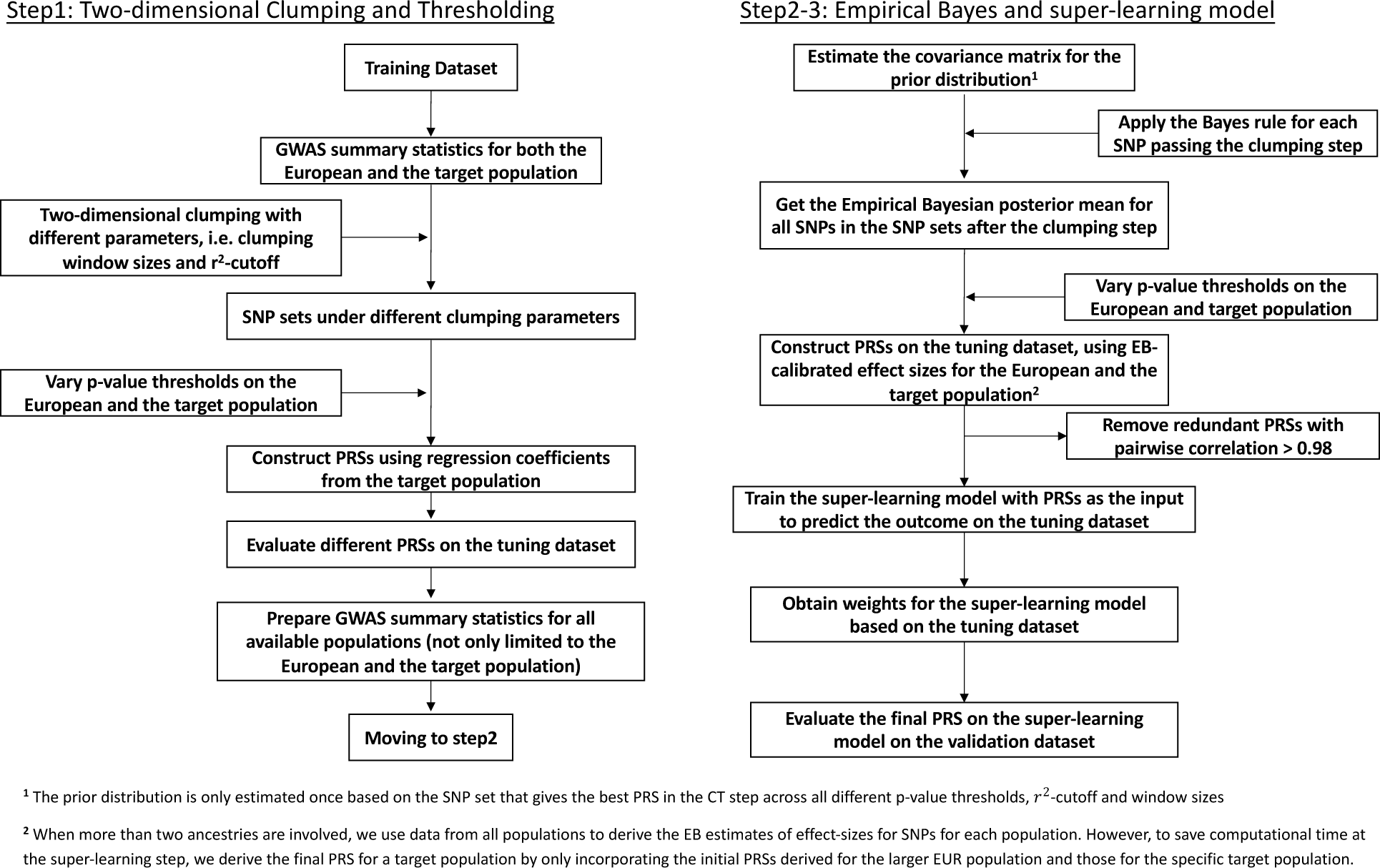
CT-SLEB detailed flowchart. The method contains three major steps: 1. Two-dimensional clumping and thresholding; 2. Empirical-Bayes procedure for utilizing genetic correlations of effect sizes across populations; 3. Super-learning model for combining PRSs under different tuning parameters. The tuning dataset is used to train the super learning model. The final prediction performance is evaluated based on an independent validation dataset. For continuous traits, the prediction is evaluated using R^2^ obtained from the linear regression between outcome and PRS after adjusting for covariates (**Methods**). For binary traits, the prediction is evaluated using area under the ROC curve (AUC).

**Extended Data Figure 2:**
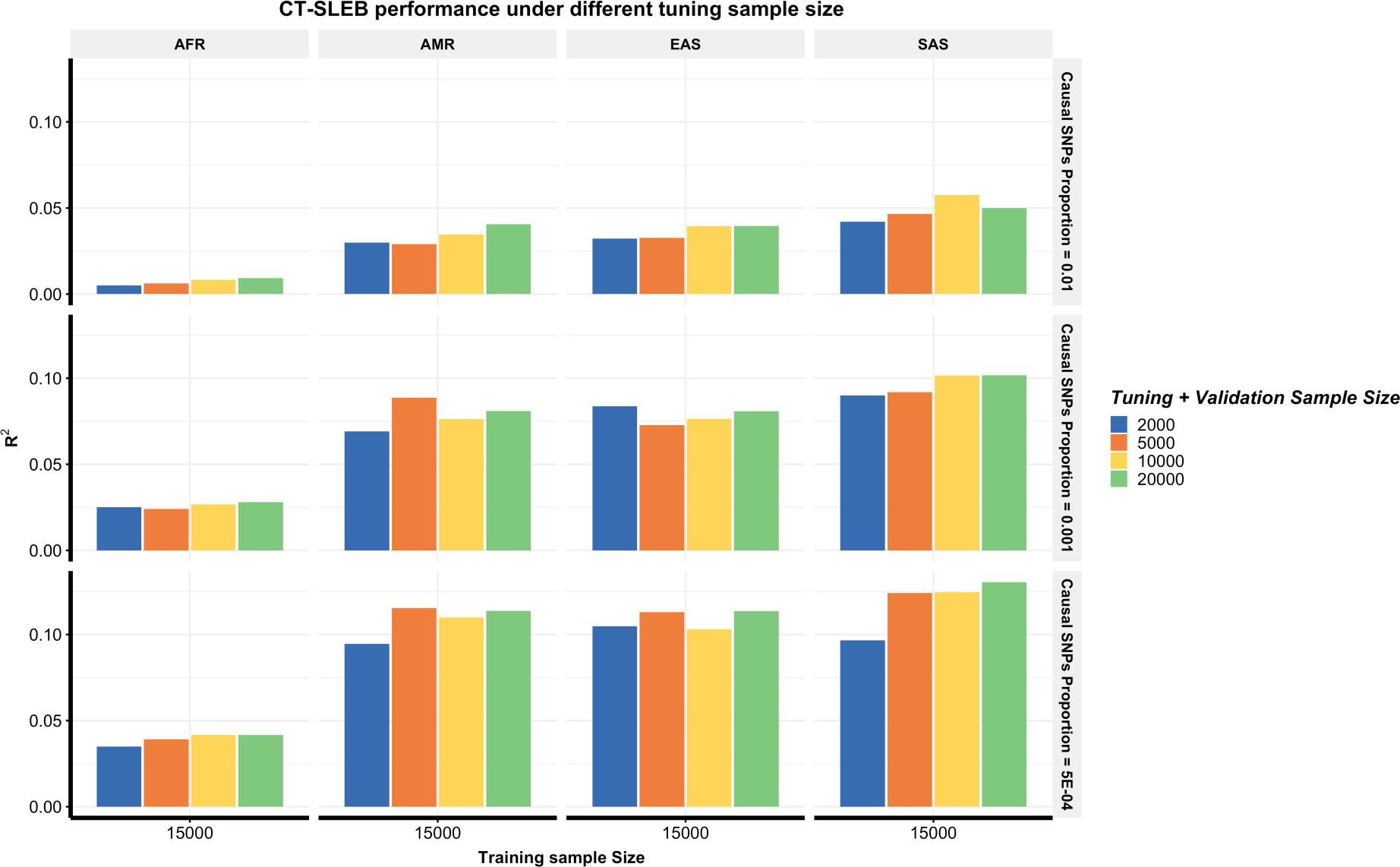
Performance of CT-SLEB with different tuning and validation sample sizes. The total tuning and validation sample size is set as 2000, 5000, 100,000 and 200,000 with half for tuning and half for validation. Analyses are conducted in the multi-ancestry setting under a strong negative selection model. The training sample size for AFR population is 15,000. The training sample size for EUR is 100,000. The sample size for the tuning dataset and validation for each population is fixed at 10,000, respectively. Common SNP heritability is assumed to be 0.4 across all populations and effect-size correlation is assumed to be 0.8 across populations. The causal SNPs proportion are varied across 0.01 (top panel), 0.001 (medium panel), or 5×10^−4^ (bottom panel). The final prediction R^2^ is reported as the average of ten independent simulation replicates.

## Methods

We assume there are *l* = 1, … *L* populations, with *l* = 1 indexing the EUR population. We assume that for each population, summary-statistics data from underlying GWAS are available in the form of (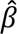_*k*l_, 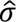_*k*l_, *p*_*k*l_) for *k* = 1,2, …, *K*_/_ SNPs, where 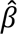, 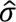 and *p* denote effect-size estimates, standard errors and p-values for individual SNPs, respectively. We also assume additional datasets are available for each target population, which can be split into tuning and validation sets. Our proposed CT-SLEB method contains three steps: 1. Two-dimensional clumping and thresholding (CT); 2. EB procedure; 3. Super-learning algorithm, detailed in the following subsections.

**CT.** In this step, we extend the traditional CT to a two-dimensional setting so that PRS for a target population can be built using approximately independent SNPs that show significant association in at least one of the two populations (majority population and target population). The CT method has two components, Clumping and Thresholding. In the two-dimensional setting where the lead SNPs might be informed by GWAS of either the EUR or target population, it is unclear which reference sample is the most suited for LD clumping. After initial exploration of alternative approaches through simulation studies, we find the most informative approach is to split the SNPs into two sets depending on which population they shows stronger signals and then perform LD clumping for each set separately based on the reference sample from respective populations. For the thresholding step, we select SNPs based on two distinct thresholds for their respective p-values in the two populations. As the optimal threshold for p-value selection depends on sample size^18,35,36^, and sample sizes for GWAS across EUR and minority populations are highly differential, we anticipate (and confirm through simulation studies) that a two-dimensional approach for threshold selection is more optimal than using a single p-value threshold across both populations. The CT step details are as follows:

1. The clumping *r*^2^-cutoff and base size of the clumping window size *w*_0_ vary across (0.01, 0.05, 0.1, 0.2, 0.5, 0.8) and (50kb, 100kb), respectively. The clumping window size *w_s_* is defined as *w_b_*/*r*^2^ because LD is inversely proportional to the genetic distance between variants^20,61^.
2. Select all variants with smaller p-values in EUR (*p*_*k*1_ < *p*_*k*2_), and clump based on *p*_*k*1_ using LD estimates from EUR reference samples with selected *r*^2^ and *w_s_*.
3. Select all variants with smaller p-values in the target population (*p_k2_* < *p*_*k*1_) and the population-specific SNPs, and then, clump based on *p*_*k*2_ using LD estimates from the target population’s reference samples with the same *r*^2^-cutoff and *w_b_*.
4. Combine the post-clumping variants from the step 2 and 3 as the candidate variants set.
5. Define two different p-value cutoffs (*p*_*t*1_, *p*_*t*2_) for EUR and the target population. A variant is selected if *p*_*k*1_ < *p*_*t*1_ or *p*_*t*1_ < *p*_*t*2_. We allow *p*_*t*1_ and *p*_*t*2_ to vary in the set: (5 × 10^−8^, 5 × 10^−7^, 5 × 10^−6^, …, 5 × 10^−1^, 1.0). With the cross combination of *p_t1_* and *p*_1*_, a total of 81 different p-value cutoffs are applied.
6. With the cross combination of *p*_*t*1_, *p*_*t*2_, *r*^2^ and *w_b_*, a total of 972 PRSs (6 *r*^2^ thresholds * 2 *w_b_* windows * 81 value thresholds) are evaluated on the tuning dataset using estimated regression coefficients (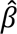_*k*2_) from the target population’s GWAS.

### EB to calibrate regression coefficients

In the CT step, we use 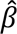_*k*2_ from the target population to calculate PRS. However, 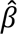_*k*2_ can be noisy with small target population GWAS sample sizes. Meanwhile, given high genetic correlations across ancestries^38,39^, effect sizes from other populations can calibrate regression coefficients for PRS. Although we only use p-values from GWAS for the EUR and the target population for selecting SNPs in the CT step, the EB step leverages GWAS from multiple populations. Suppose (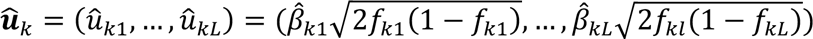), is the vector of the standardized effect-size for the *k*th SNP in L different populations, with 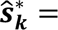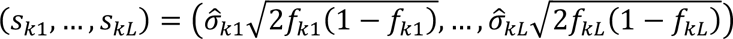 being the vector of the corresponding standard errors of *û_k_*. We assume that 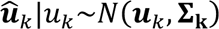, where 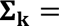 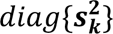 given that the GWAS for different populations are independent. Additionally, we assume that the prior distribution of the mean of *û_k_* is *u_k_* ∼ *N*(**0, Σ**_**0**_), which assumes the effect-size follows the strong negative selection model. By integrating the conditional and prior distribution, we obtain the marginal distribution of *û_k_* as *N*(**0, Σ**_**0**_ + **Σ**_**k**_). Suppose the SNP set selected from the CT step has *K*^∗^ variants overlapped across all the populations. We estimate the prior covariance matrix **Σ**_**0**_ using the *K*^*^ overlapped variants shared across all populations as:

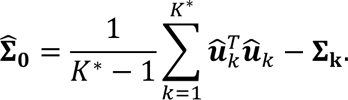

We note that we ignore potential correlation across selected SNPs in this step, but the estimate is still expected to be consistent for **Σ**_**0**_ which represents marginal variance-covariance matrices for effect sizes associated with an individual SNP across populations. Applying the Bayes formula, the posterior distribution of *u_k_* becomes

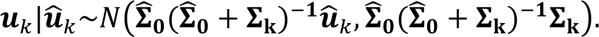

The EB coefficients for the *k*th SNP are defined as:

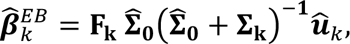

where 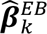 is a *L* × 1 vector for the posterior effect-size of the kth SNP for in each of the L populations, and 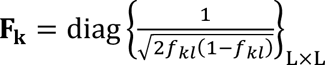 is the scaling matrix to scale effect sizes from the standardized scale back to the original scale. Preliminary simulation studies indicate that EB step of effect-size calibration leads to distinct improvement in PRS performance (compared to using effect-size estimates from the target population) irrespective of all other steps.

To save computational time, we estimate 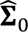), based on the SNP set that gives the best PRS in the CT step, and apply the same 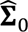) to derive the EB-calibrated effect sizes for all PRSs corresponding to cross combination of *p*_*t*1_, *p*_*t*2_, *r*^2^-cutoff and *w_b_*. Using EB-calibrated effect sizes, we compute 972 PRSs for each population (6 *r*^2^ thresholds * 2 *w_b_* windows * 81 value thresholds). In all analyses, we compute the 1944 PRSs using EB-calibrated effect sizes of the target population and EUR. When more than two ancestries are involved, we use data from all populations to derive EB estimates (**Supplementary Note**). However, to save computational time at the super-learning step, we derive the final PRS for the target population by only incorporating the 1944 PRSs derived for the larger EUR population and the target population. All 1944 PRSs are used as input for the super-learning step to predict the outcome for the target population. Because many PRSs are highly correlated, we filter out redundant ones with pairwise correlations higher than 0.98. In the simulations, 369 of the 1994 PRSs are kept on average after the filtering.

### Super learning

We combine all PRSs generated from the previous steps into an input dataset and train them on the tuning dataset to predict the outcome *Y*. The super-learning algorithm generates an optimally weighted combination from a set of distinct prediction algorithms^40–42,62^ (**Supplementary Note**). The set of prediction algorithms can be self-designed or chosen from classical prediction algorithms e.g., Lasso^44^, ridge regression^45^, neural networks^46^, etc. We use three different prediction algorithms implemented in the SuperLearner package^43^ to generate the super learning estimate: Lasso^44^, ridge regression^45^ and neural networks^46^. For binary traits, since ridge regression is currently not supported by the SuperLearner package, we use Lasso and neural network in data analysis. To use AUC as the objective function, we use the flag “method = method.AUC” in the SupearLearner package.

### Simulation

Large-scale multi-ancestry genotype data are generated using HAPGEN2 (version 2.1.2)^63^ mimicking the LD of EUR, AFR, Americas (AMR), East Asia (EAS) and South Asia (SAS). The 1000 Genomes Project (Phase 3)^51^ serves as the reference panel, including 503 EUR, 661 AFR, 347 AMR, 504 EAS and 489 SAS subjects. Biallelic SNPs with MAF more than 0.01 in any of the populations are kept, resulting in ∼8.6 million SNPs for EUR, ∼14.8 million SNPs for AFR, ∼9.8 million SNPs for AMR, ∼7.6 million SNPs for EAS, and ∼9.0 million SNPs for SAS. The genotype data are generated with a total of ∼19.2 million SNPs. The set of simulated variants for all five ancestries is same. Population-specific SNP proportions range from 2.92% for AMR to 43.84% for AFR, respectively (**Supplementary Fig. 22**). SNPs with MAF below 0.01 in a population are excluded from PRS calculation due to unstable LD estimation. A total of 120,000 independent subjects are simulated for each of the population.

Trait values are generated by selecting causal SNPs randomly across the genome, with the causal SNP proportion set to 0.01, 0.001, or 5 × 10^−4^. We consider two models for heritability distribution: (A) Constant common-SNP heritability. (B) Constant per-SNP heritability that implies the total heritability is proportional to the number of common SNPs. We also consider three negative selection patterns: strong, mild and no negative selection.

We denote *u_kl_* as the standardized effect-size for *k*th causal SNP for the *l*th population. Under strong negative selection and constant heritability model, standardized effect-sizes are drawn from a multivariate normal distribution of the form:

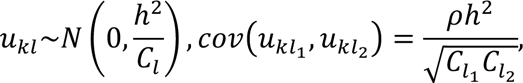

where *C_l_* is the number of causal SNPs with MAF > 0.01 in the *l*th population, the heritability *h*^2^ associated with common SNPs for each population is set to 0.4, and the genetic correlation *ρ* is set to 0.8. We then generate the phenotype using linear model of the form 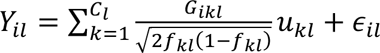 for the *i*th subject in the *l*th population, where *f_kl_*is the effect allele frequency for the *k*th causal SNP in *l*th population. The error terms are generated as *ε_il_* ∼ *N*(0,1 − *h^2^*). We also consider mild negative selection (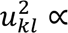 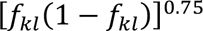) and no negative selection (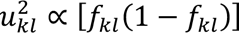) scenarios (**Supplemental Notes**). Finally, we assume total heritability of all ∼19 million SNPs to be 0.4 across all populations, but common SNP heritability varying proportionately to their number within each the populations. The model assumes equal per SNP heritability across populations, leading to the common SNP heritability values of 0.32, 0.21, 0.16, 0.19 and 0.17 for AFR, AMR, EAS, EUR and SAS, respectively. Genetic correlation is set to 0.8 or 0.6.

We set the training sample sizes for each target population to 15,000, 45,000, 80,000, or 100,000. GWAS summary statistics for each population are generated based on the training samples using PLINK version 1.90 with the command “--linear”. We fixed the EUR sample size at 100,000, and simulate the tuning and validation dataset of 10,000 for each target population. The final prediction *R*^2^ is the average of ten independent simulation replicates. For CT-SLEB and PRS-CSx, incorporating data across all five ancestries, we assume non-EUR training sample sizes to be equal to the target population’s.

### Existing PRS methods

The CT method selects clumped SNPs with varying p-value thresholds and chooses an optimal PRS based on its performance on the tuning dataset. We implement CT using PLINK version 1.90^64^ with the clumping step command “--clump --clump-r2 0.1 --clump-kb 500”. We estimate LD based on 3,000 randomly selected unrelated subjects from the training dataset for each population. We set candidate p-value thresholds to be (5 × 10^−8^, 1 × 10^−7^, 5 × 10^−7^, 1 × 10^−6^, …, 5 × 10^−1^, 1.0) and we use the PLINK command “--score no-sum no-mean-imputation” for computing PRS. The optimal p-value threshold is determined based prediction R^2^ on the tuning dataset.

The LDpred2 method infers SNP effect sizes by a shrinkage estimator, combining GWAS summary statistics with a prior on effect sizes while leveraging LD information from an external reference panel. LDpred2 is implemented using the R package “bigsnpr”^26^. The tuning parameters are: (1) the proportion of causal SNPs, with candidate values set to a sequence of length 17 that are evenly spaced on a logarithmic scale from 10^−4^ to 1; (2) per-SNP heritability, with candidate values set to 0.7, 1, or 1.4 times the total heritability estimated by LD score regression divided by the number of causal SNPs; (3) “sparse” option, which is set to “yes” or “no” (the “sparse” option sets some weak effects to zero). The method selects tuning parameters based on the performance on the tuning dataset.

The EUR PRS based on CT or LDpred2 are built using EUR training dataset from the EUR population and estimates tuning parameters based on tne EUR tuning sample. When evaluating the EUR PRSs in the target population, we exclude the SNPs that do not exist in the target population.

Weighted-PRS linearly combines the CT or LDpred2 PRS generated from the EUR and target populations. The weights for EUR PRS and for target population PRS are estimated using the target population’s tuning dataset through a linear regression. We implement weighted-PRS using R version 4.0.0.

PolyPred-S+ consists of a PolyFun-pred predictor trained on the EUR population and two SBayesR predictors using training data from the EUR and target populations respectively. On a target population tuning dataset, PolyPred-S+ performs non-negative least squares regression to compute the mixture weights and linearly combines the predictors. PolyFun-pred leverages genome-wise functional annotations for prior causal probabilities, fed into the SuSiE fine-mapping method for the posterior causal effect estimation. SBayesR estimates posterior tagging effects with a finite normal mixture prior on effect sizes. For PolyFun-pred, we use precomputed prior causal probabilities provided by the authors, extract LD information using EUR population in 1000 Genomes Project (Phase 3), and assume 10 causal SNPs per locus. Using GCTB (2.03 beta version), we train SBayesR with the sparse shrunk LD matrix for HapMap3 variants published by the SBayesR authors. Currently, the shrunk LD matrix are only available for EUR populations. Therefore, both SBayesR predictors for EUR and target populations use the shrunk LD matrix for EUR. Model parameters and MCMC settings followthe the same way as the PolyPred-S+ authors’ UKBB simulations.

XPASS leverages the genetic correlation between the target and EUR populations, assuming a bivariate normal distribution with non-zero covariance for effect sizes pairs corresponding to the same SNP in both populations. It can incorporate population-specific covariates as fixed effects to improve weight estimation accuracy. We extract the top 20 PCs from the reference genome for each population as the covariates. When estimating LD matrices, ancestry-specific reference data is inputted. LD matrices are estimated based on EUR LD blocks for all datasets, as XPASS package only offers EUR and EAS options.

PRS-CSx estimates population-specific SNP effect sizes using a Bayesian framework using continuous shrinkage priors to jointly model GWAS summary statistics from multiple populations. Additionally, PRS-CSx conducts a step similar to weighted-PRS, linearly combining PRS based on the posterior effect-sizes from EUR and target populations, with weights estimated based on the target population’s tuning dataset. We implemented PRS-CSx following https://github.com/getian107/PRScsx. We set the gamma-gamma prior hyperparameters *a* and *b* to default values of 1 and 0.5, respectively. Further, the parameter *ϕ* is varied over the default set of values 10^−6^, 10^−4^, 10^−2^, and 1, with optimal *ϕ* is determined based on tuning dataset performances.

### Runtimes and memory usage comparison

We compare the computation time and memory usage of CT-SLEB (two ancestries and five ancestries), and PRS-CSx (two ancestries and five ancestries) based on their performance on chromosome 22, assuming AFR as the target population. All analyses use a single core with Intel E5-26840v4 CPU. Performance is averaged over 100 replicates. The training dataset consists of GWAS summary statistics for AFR (N_GWAS_=15,000) and EUR (N_GWAS_=100,000) populations. Tuning dataset and validation datasets each contain 10,000 subjects. For five ancestries analyses, training GWAS sample sizes for AMR, EAS and SAS are set to 15,000 each.

### 23andMe Data analysis

The individuals in our analyses are part of the 23andMe participant cohort. All participants provided informed consent and answered surveys online according to our human subject protocol reviewed and approved by Ethical & Independent Review Services, a private institutional review board (http://www.eandireview.com). Detailed information about genotyping, quality control, imputation, removing related individuals, and ancestry determination is provided in **Supplementary Note.** Participants were included in the analysis based on consent status as checked when data analyses were initiated.

Our analysis involves five ancestries (AA, EAS, EUR, Latino, SAS), and include two continuous and five binary traits: 1. Heart metabolic disease burden 2. Height 3. Any CVD 4. Depression 5. Migraine Diagnosis 6. Morning Person 7. SBMN. Data for each population is randomly split into training, tuning, and validation datasets (70%, 20%, and 10%, respectively), with detailed sample size in **Supplementary Table 3**. We perform GWAS for the seven traits using each population’s training dataset, adjusting for PC 1-5, sex, and age with standard quality control procedures (**Supplementary Note**). SNPs with MAF > 0.01 in at least one population are kept in the analyses. We further restrict analyses to SNPs that are on HM3 + MEGA chips with ∼2.0 million SNPs. LDSC version 1.01^52^ is used to estimate the heritability with EUR population GWAS summary statistics for the seven traits. LD scores are estimated using the 503 unrelated EUR samples from the 1000 Genomes Project. Heritability analyses are limited to EUR populations due to insufficient sample size in non-EUR populations for stable LD-score regression estimates.

We compare PRS prediction performance for ten methods: CT, LDPred2, best EUR PRS based on CT and LDpred2, weighted-PRS based on CT and LDpred2, PolyPred-S+, XPASS, PRS-CSx (using EUR and target population data or using all five populations), CT-SLEB (using EUR and target population data or using all five populations). As individual-level data is unavailable in the training step, we use 1000 Genomes Project (Phase 3) reference data to estimate LD for each population. Specifically, AFR and AMR from the 1000 Genomes Project serves as references for the AA and Latino population in 23andMe, respectively. PRS prediction performance is reported based on the independent validation dataset, separate from training and tuning datasets. To calculate the adjusted R^2^ for continuous traits, we first regress the traits on covariates and then evaluate PRS performance by predicting residualized trait values. Adjusted AUC for binary traits is calculated using roc.binary function in the R package RISCA version 1.01^65^.

### GLGC Data analysis

We obtain GWAS summary statistics of four blood lipids traits are obtained from public available GLGC database (http://csg.sph.umich.edu/willer/public/glgc-lipids2021/results/ancestry_specific/). UKBB data are removed from the GWAS summary statistics. The details of study design of study, genotyping, quality control and GWAS are described elsewhere^37^. Training data are available for four blood lipid traits: LDL, HDL, logTG, TC from five different ancestries: EUR, AFR (primarily AA), Latino, EAS, and SAS (**Supplementary Table 8**). Tuning + validation data from the UKBB dataset are from EUR, AFR, EAS and SAS ancestries (**Supplementary Table 9**). Details of ancestry prediction of UKBB are described in the **Supplementary Note.** Due to poor ancestry classification and low sample size, the Latino population is not evaluated using UKBB data. The implementation of the ten different PRSs approaches follow the same steps as in the 23andMe data analyses. We use the 1000 Genomes Project (Phase 3) reference data to estimate the LD for each population. The adjusted R^2^ are adjusted by sex, age and genetic PC1-10.

### AoU Data analysis

The individuals included in our analyses are part of the AoU participant cohort. All these individuals’ information have been collected according to the AoU Research Program Operational Protocol (https://allofus.nih.gov/sites/default/files/aou_operational_protocol_v1.7_mar_2018.pdf). Detailed information about genotyping, ancestry determination, quality control, removing related individuals can be found in AoU Research Program Genomic Research Data Quality Report (https://www.researchallofus.org/wp-content/themes/research-hub-wordpress-theme/media/2022/06/All%20Of%20Us%20Q2%202022%20Release%20Genomic%20Quality%20Report.pdf).

We analyze three ancestries (EUR, AFR, and Latino/Admixed American) and two continuous traits (height and BMI). GWAS for these traits are performed using unrelated samples for each population, adjusting for PC 1-16, sex, and age, with quality control steps provided by the AoU platform.

The AoU platform provides whole genome sequencing (WGS) data and array data. While the WGS data has fewer samples than Array data (98,590 WGS and 165,127 array samples, June 22, 2022, version), quality control information and relatedness of samples are only provided within the WGS data. For GWAS with respect to each population, we perform sample-level quality control within WGS data. Due to computation burden, analyses are conducted using array SNPs with subjects passing the WGS data quality control. SNPs with MAF > 0.01 in at least one of the three populations are kept, while analyses are restricted to SNPs available on HM3 + MEGA chips. Since the analyses are constrained to array data, all analyses involve up to 800K SNPs (**Supplementary Table 4**). We use the reference data from the 1000 Genomes Project (Phase 3) to estimate the LD for each population.

Tuning + validation data from the UKBB dataset are from EUR and AFR ancestries. Latino population is not evaluated on UKBB for the same reason as GLGC analyses. The implementation of the ten different PRSs approaches follow the same steps as the 23andMe data analyses. The adjusted R^2^ are adjusted by sex, age and PC1-10.

## Data availability

Simulated genotype data for 600K subjects from five ancestries: https://dataverse.harvard.edu/dataset.xhtml?persistentId=doi:10.7910/DVN/COXHAP GWAS summary level statistics for five ancestries from GLGC: http://csg.sph.umich.edu/willer/public/glgc-lipids2021/results/ancestry_specific/ GWAS summary statistics for three ancestries from AoU: https://dataverse.harvard.edu/dataset.xhtml?persistentId=doi:10.7910/DVN/FAWEQK The PRS developed for six traits for GLGC and AoU are released through PGS Catalog (https://www.pgscatalog.org/) with publication ID: PGP000489 and score IDs: PGS003767-PGS003848.

The 23andMe GWAS summary statistics for top 10,000 genetic markers associated with three traits (height, morning person, and sing back musical note) across five diverse ancestries has been made available as **Supplementary Data** and also available: https://dataverse.harvard.edu/dataset.xhtml?persistentId=doi:10.7910/DVN/3NBNCV. The full GWAS summary statistics and the final PRS for these three traits (height, morning person, and sing back musical note) are available through 23andMe to qualified researchers under an agreement with 23andMe that protects the privacy of the 23andMe participants. Please visit research.23andme.com/dataset-access/ for more information and to apply to access the data. The summary statistics for the four other traits used in the paper (any CVD, heart metabolic disease burden, depression, and migraine) will not be made available because of 23andMe’s business requirements.

Participants provided informed consent and participated in the research online, under a protocol approved by the external AAHRPP-accredited IRB, Ethical & Independent Review Services.

## Code Availability

Simulation and data analyses code is available at: GitHub (https://github.com/andrewhaoyu/multi_ethnic^66^

Software implementing CT-SLEB is available at: GitHub (https://github.com/andrewhaoyu/CTSLEB^67^

P + T: https://www.cog-genomics.org/plink/1.9/

SCT and LDpred2: https://github.com/privefl/bigsnpr.

XPASS: https://github.com/YangLabHKUST/XPASS

PolyPred-S+: https://github.com/omerwe/polyfun

PRS-CSx: https://github.com/getian107/PRScsx

LDSC: https://github.com/bulik/ldsc

PLINK: https://www.cog-genomics.org/plink/1.9/

The majority of our statistical analyses are performed using the following R packages: ggplot2 version 3.3.3, dplyr version 1.0.4, data.table version 1.13.6, bigsnpr version 1.6.1, SuperLearner version 2.0.26, caret version 6.0.86, ranger version 0.12.1, glmnet version 4.1, RISCA version 1.01, XPASS version 0.1.0, xgboost version 1.7.5.1, randomForest version

