## Supplementary Figure and Note for "A new method for multi-ancestry polygenic prediction improves performance across diverse populations"

**Supplementary Figure 1: Simulation performances of different PRSs methods in multi-ancestry settings under <u>a strong negative selection model</u>.** Each of the four non-EUR populations has a training sample size of 45,000 (Sup. Fig. 2a) or 100,000 (Sup. Fig. 2b). For the EUR population, the size of the training sample is set at 100,000. The number of samples for each population in the tuning dataset is always 10,000. Prediction $R^2$ values are reported based on an independent validation dataset with 10,000 subjects for each population. Common SNP heritability is assumed to be 0.4 across all populations, and effect-size correlation is assumed to be 0.8 across all pairs of populations. The causal SNPs proportion (degree of polygenicity) is varied across 0.01 (top panel), 0.001 (medium panel), $5 \times 10^{-4}$ (bottom panel). All data are generated based on ~19 million common SNPs across the five populations, but analyses are restricted to ~2.0 million SNPs that are used on Hapmap3 + Multi-Ethnic Genotyping Arrays chip. PolyPred-S+ and PRS-CSx analyses are further restricted to ~1.3 million HM3 SNPs. All approaches are trained using data from the EUR and the target population. CT-SLEB and PRS-CSx are further evaluated using training data from five ancestries. The red dashed line represents the prediction performance of EUR PRS generated using single ancestry method (best of CT or LDpred2) in the EUR population.

**a)**

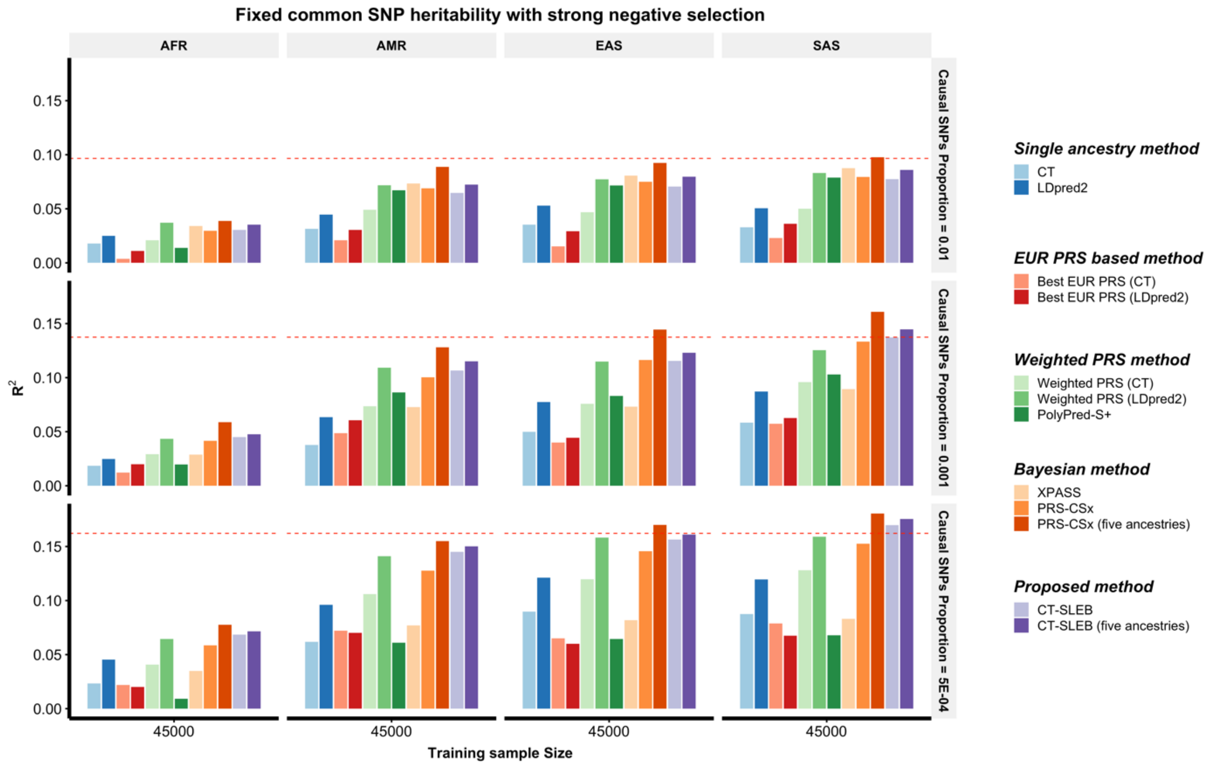

**Supplementary Figure 1 continued - Panel B: Simulation with training sample size of 100,000 for each of the four non-EUR populations.**

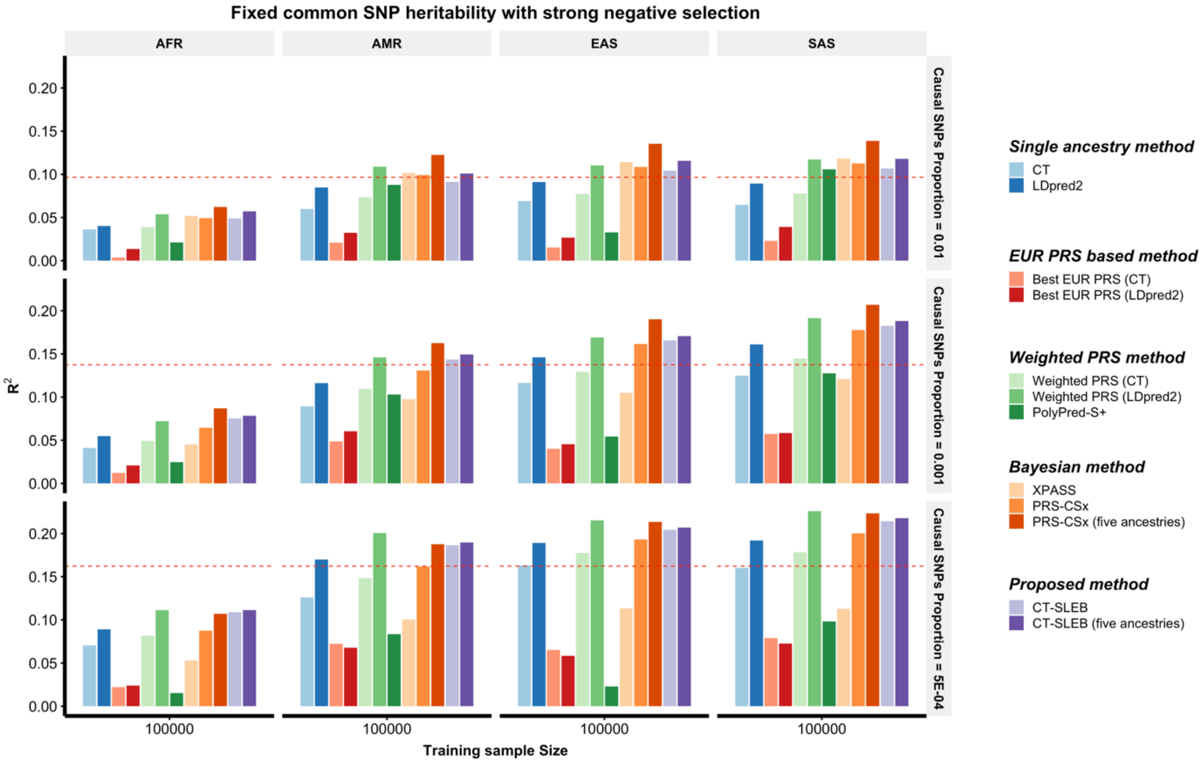

**Supplementary Figure 2: Simulation performances of different PRSs methods in multi-ancestry settings under a mild negative selection model.** Each of the four non-EUR populations has a training sample size of 15,000 (Sup. Fig. 3a), 45,000 (Sup. Fig. 3b), 80,000 (Sup. Fig. 3c) or 100,000 (Sup. Fig. 3d). For the EUR population, the size of the training sample is set at 100,000. The number of samples for each population in the tuning dataset is always 10,000. Prediction $R^2$ values are reported based on an independent validation dataset with 10,000 subjects for each population. Common SNP heritability is assumed to be 0.4 across all populations, and effect-size correlation is assumed to be 0.8 across all pairs of populations. The causal SNPs proportion (degree of polygenicity) is varied across 0.01 (top panel), 0.001 (medium panel), $5 \times 10^{-4}$ (bottom panel). All data are generated based on ~19 million common SNPs across the five populations, but analyses are restricted to ~2.0 million SNPs that are used on Hapmap3 + Multi-Ethnic Genotyping Arrays chip. PolyPred-S+ and PRS-CSx analyses are further restricted to ~1.3 million HM3 SNPs. All approaches are trained using data from the EUR and the target population. CT-SLEB and PRS-CSx are further evaluated using training data from five ancestries. The red dashed line shows the prediction performance of EUR PRS generated using single ancestry method (best of CT or LDpred2) in the EUR population.

**a)**

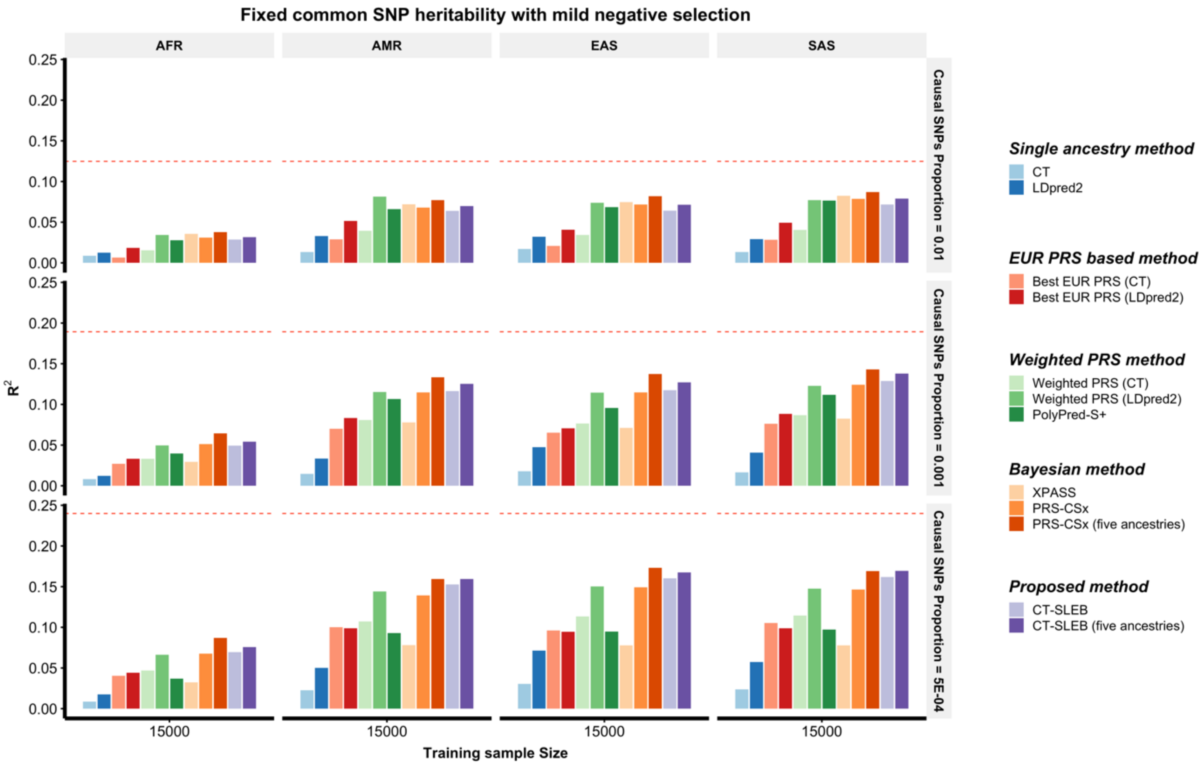

**Supplementary Figure 2 continued - Panel b: Simulation with training sample size of 45,000 for each of the four non-EUR populations.**

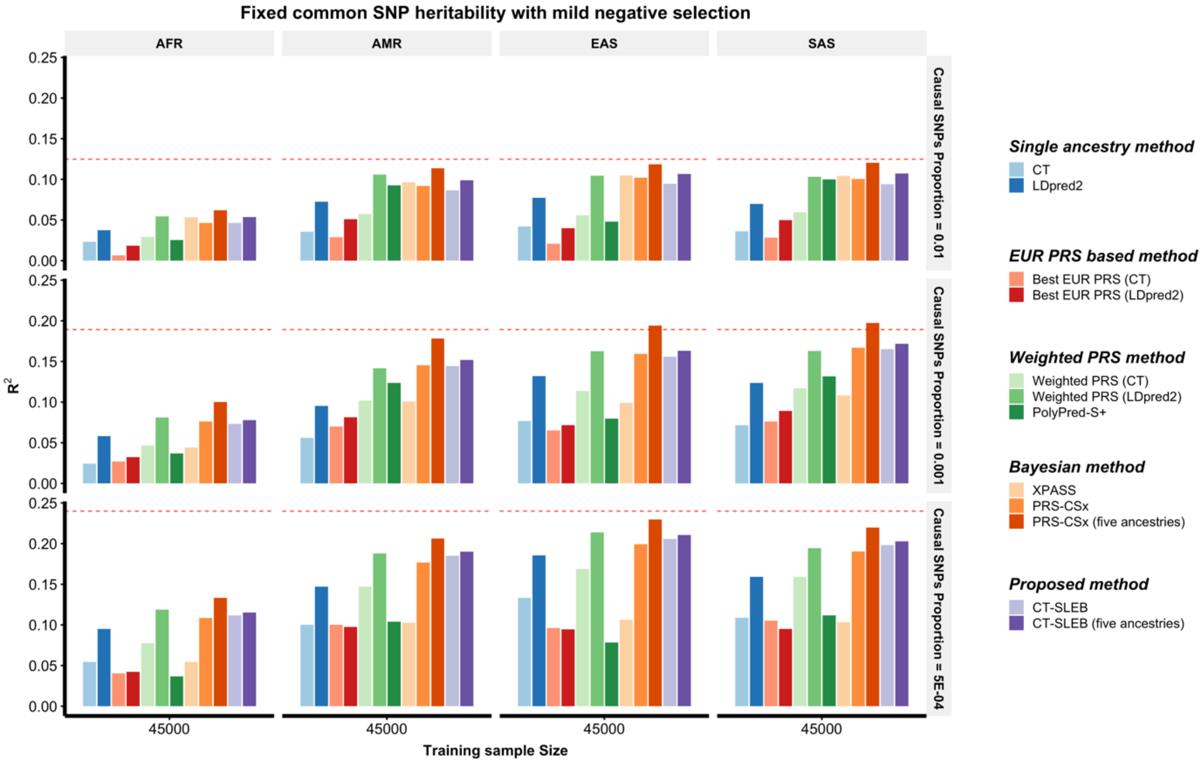

**Supplementary Figure 2 continued - Panel c: Simulation with training sample size of 80,000 for each of the four non-EUR populations.**

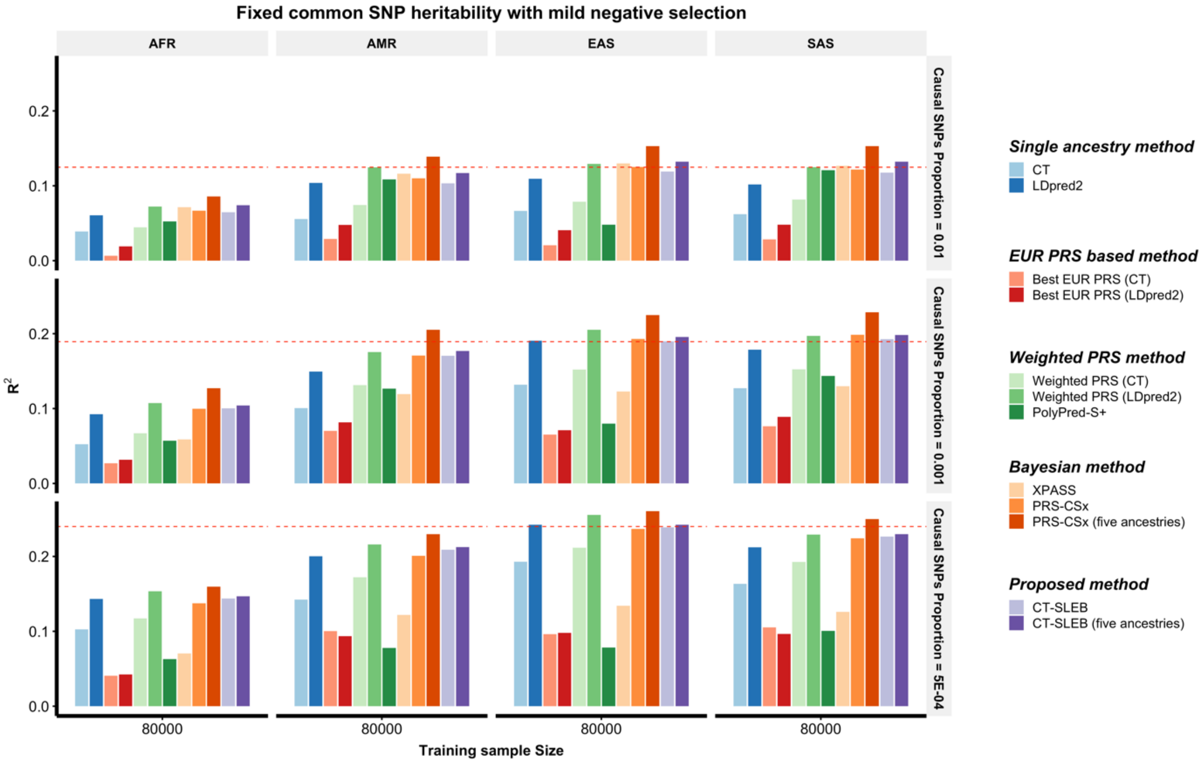

**Supplementary Figure 2 continued - Panel d: Simulation with training sample size of 100,000 for each of the four non-EUR populations.**

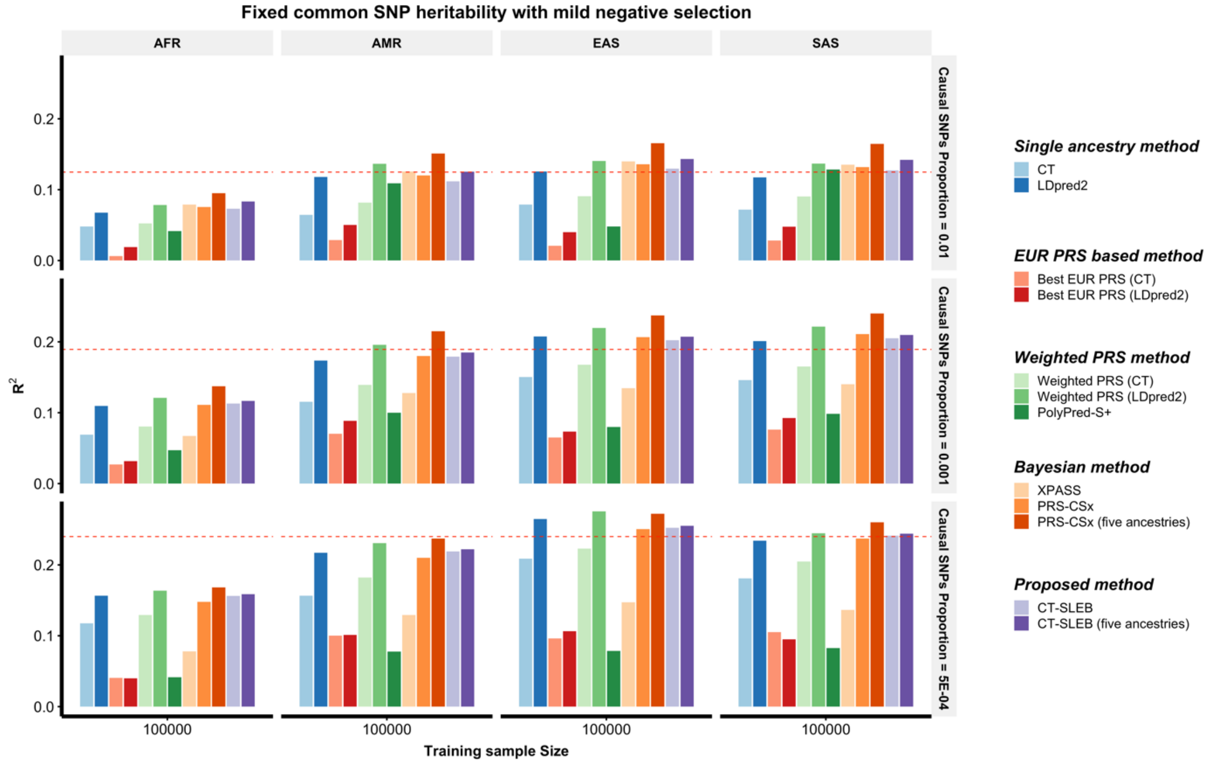

**Supplementary Figure 3: Simulation performances of different PRSs methods in multi-ancestry settings under <u>a no negative selection model</u>.** Each of the four non-EUR populations has a training sample size of 15,000 (Sup. Fig. 4a), 45,000 (Sup. Fig. 4b), 80,000 (Sup. Fig. 4c) or 100,000 (Sup. Fig. 4d). For the EUR population, the size of the training sample is set at 100,000. The number of samples for each population in the tuning dataset is always 10,000. Prediction $R^2$ values are reported based on an independent validation dataset with 10,000 subjects for each population. Common SNP heritability is assumed to be 0.4 across all populations, and effect-size correlation is assumed to be 0.8 across all pairs of populations. The causal SNPs proportion (degree of polygenicity) is varied across 0.01 (top panel), 0.001 (medium panel), $5 \times 10^{-4}$ (bottom panel). All data are generated based on ~19 million common SNPs across the five populations, but analyses are restricted to ~2.0 million SNPs that are used on Hapmap3 + Multi-Ethnic Genotyping Arrays chip. PolyPred-S+ and PRS-CSx analysis is further restricted to ~1.3 million HM3 SNPs. All approaches are trained using data from the EUR and the target population. CT-SLEB and PRS-CSx are further evaluated using training data from five ancestries. The red dashed line shows the prediction performance of EUR PRS generated using single ancestry method (best of CT or LDpred2) in the EUR population.

**a)**

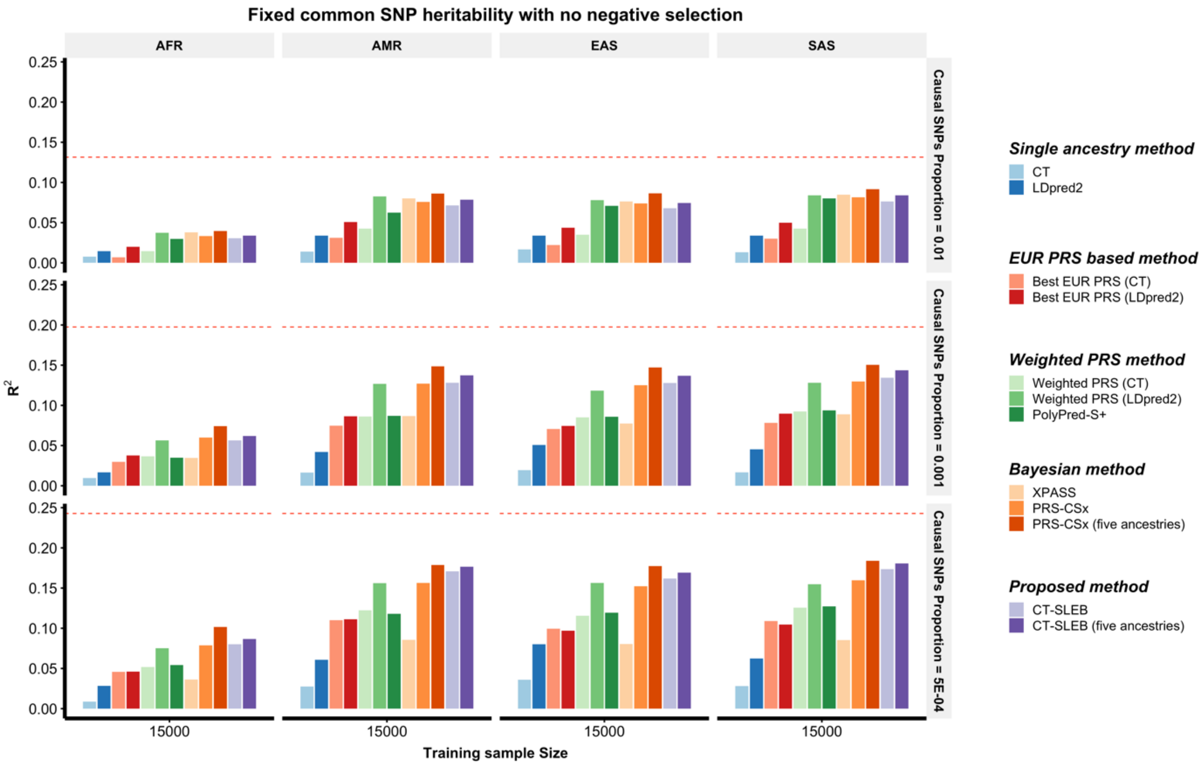

**Supplementary Figure 3 continued – Panel b: Simulation with training sample size of 45,000 for each of the four non-EUR populations.**

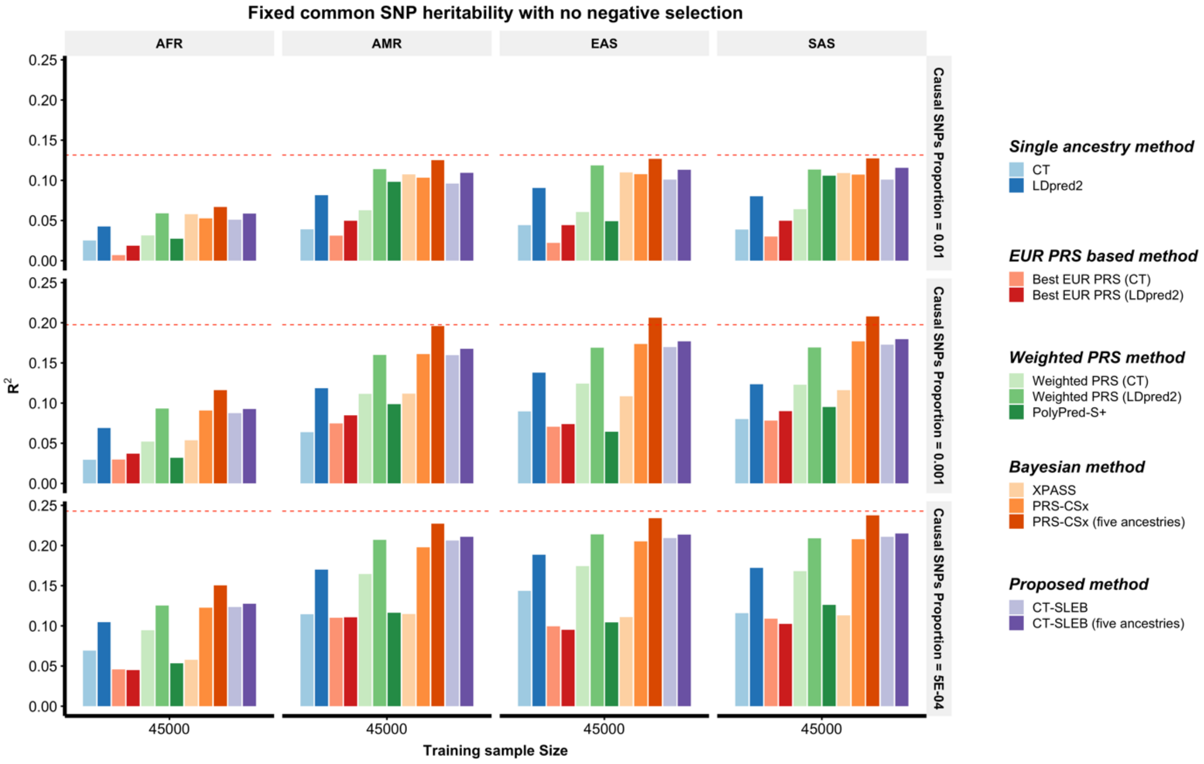

**Supplementary Figure 3 continued – Panel c: Simulation with training sample size of 80,000 for each of the four non-EUR populations.**

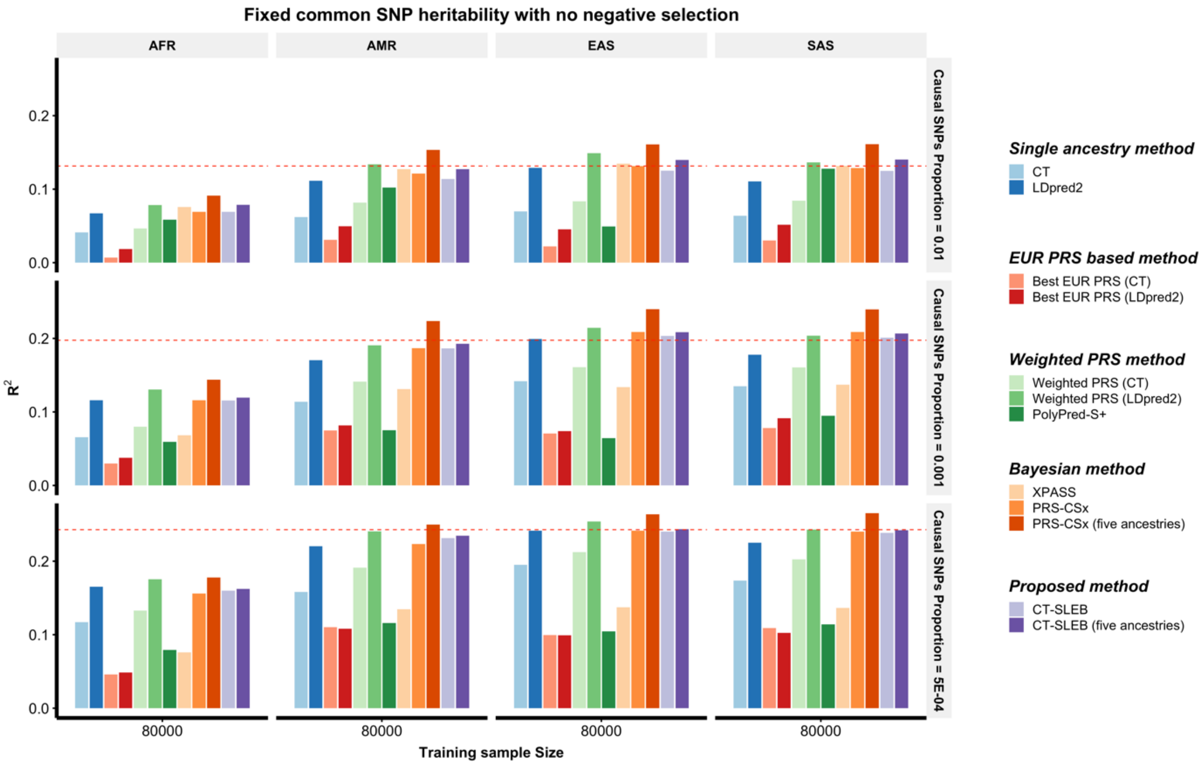

**Supplementary Figure 3 continued – Panel d: Simulation with training sample size of 100,000 for each of the four non-EUR populations.**

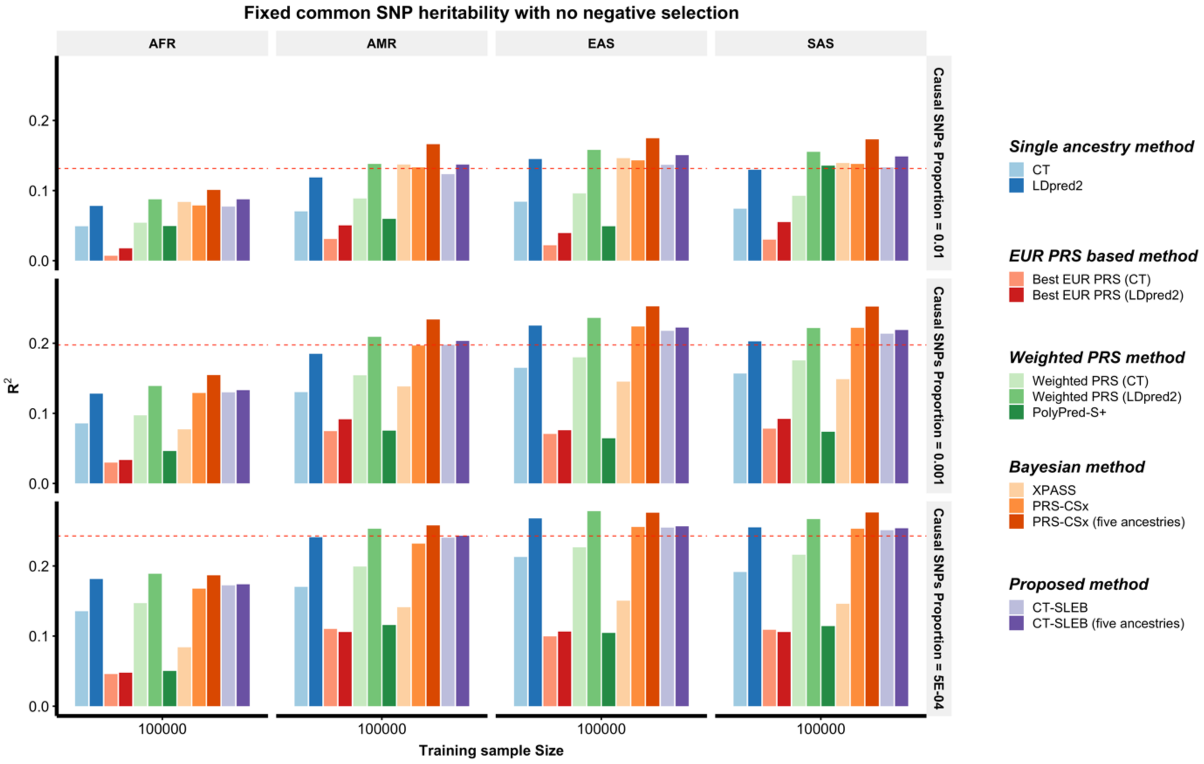

**Supplementary Figure 4: Simulation performances of different PRSs methods in multi-ancestry settings under a strong negative selection model with per-SNP heritability fixed across all populations.** Each of the four non-EUR populations has a training sample size of 15,000 (Sup. Fig. 5a), 45,000 (Sup. Fig. 5b), 80,000 (Sup. Fig. 5c) or 100,000 (Sup. Fig. 5d). For the EUR population, the size of the training sample is set at 100,000. The number of samples for each population in the tuning dataset is always 10,000. Prediction $R^2$ values are reported based on an independent validation dataset with 10,000 subjects for each population. Common SNP heritability is assumed to be 0.4 across all populations, and effect-size correlation is assumed to be 0.8 across all pairs of populations. The causal SNPs proportion (degree of polygenicity) is varied across 0.01 (top panel), 0.001 (medium panel), $5 \times 10^{-4}$ (bottom panel). All data are generated based on ~19 million common SNPs across the five populations, but analyses are restricted to ~2.0 million SNPs that are used on Hapmap3 + Multi-Ethnic Genotyping Arrays chip. PolyPred-S+ and PRS-CSx analysis is further restricted to ~1.3 million HM3 SNPs. All approaches are trained using data from the EUR and the target population. CT-SLEB and PRS-CSx are further evaluated using training data from five ancestries. The red dashed line shows the prediction performance of EUR PRS generated using single ancestry method (best of CT or LDpred2) in the EUR population.

**a)**

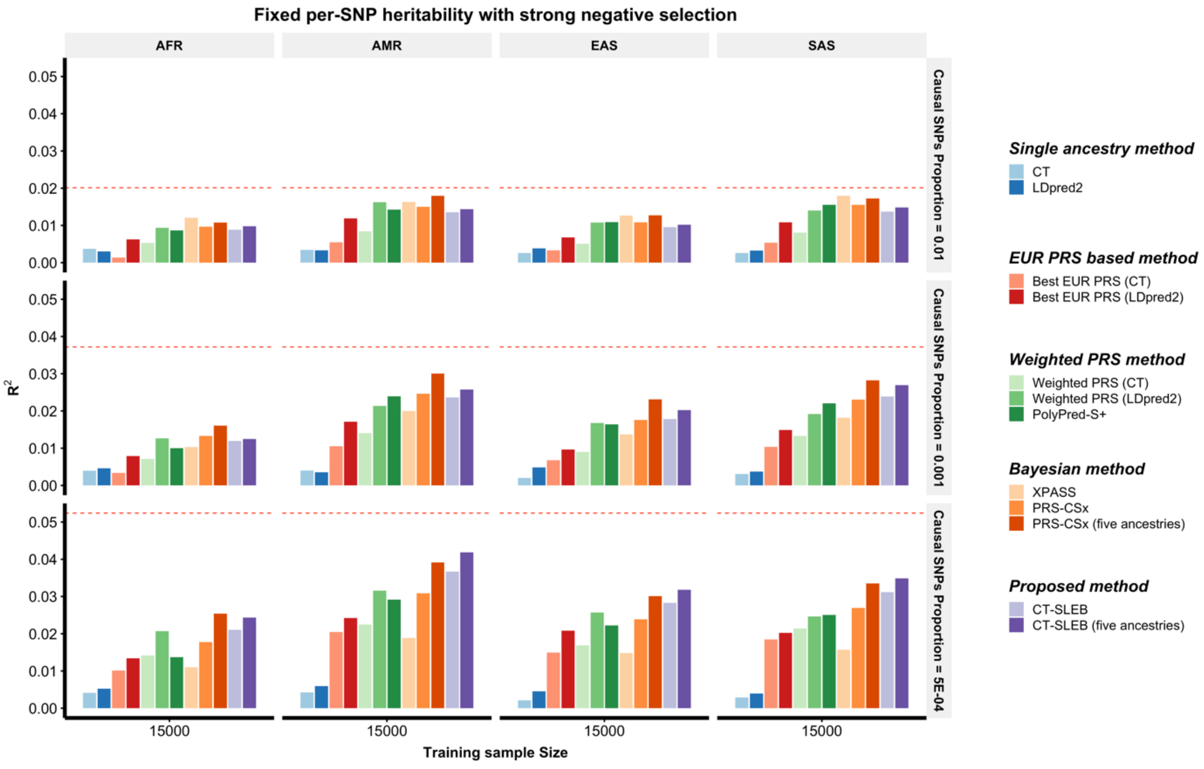

**Supplementary Figure 4 continued – Panel b: Simulation with training sample size of 45,000 for each of the four non-EUR populations.**

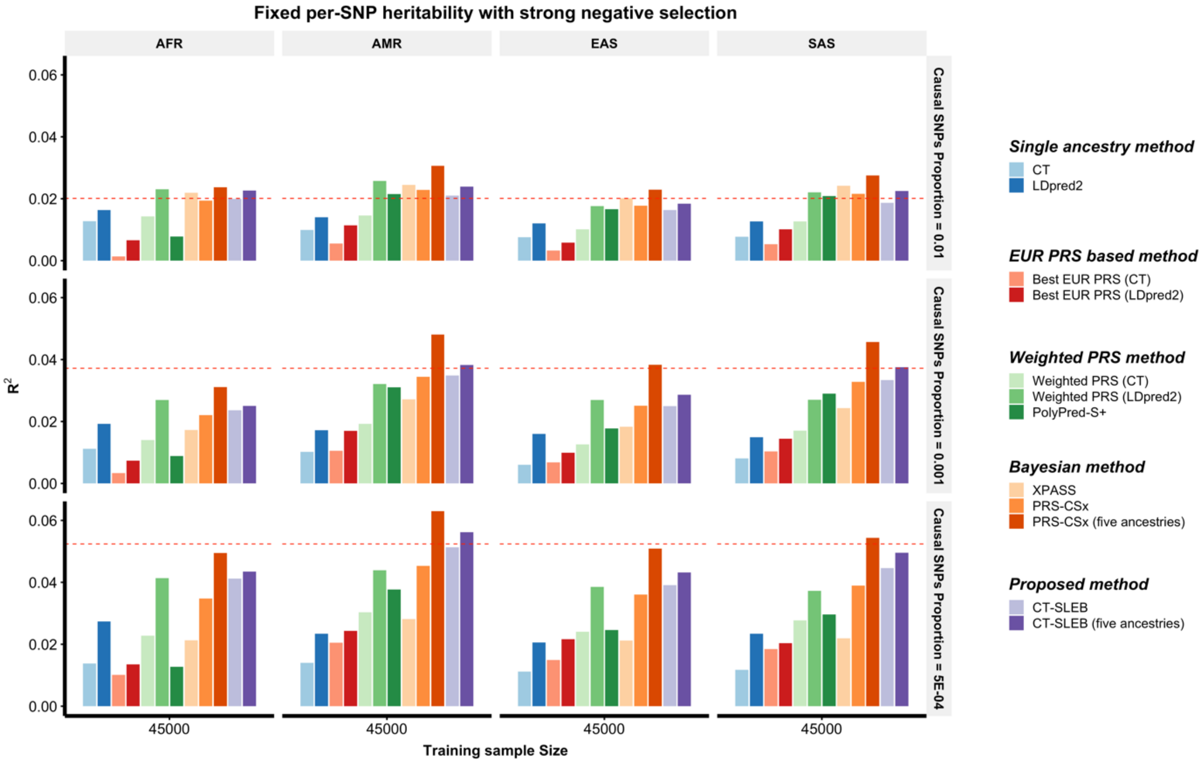

**Supplementary Figure 4 continued – Panel c: Simulation with training sample size of 80,000 for each of the four non-EUR populations.**

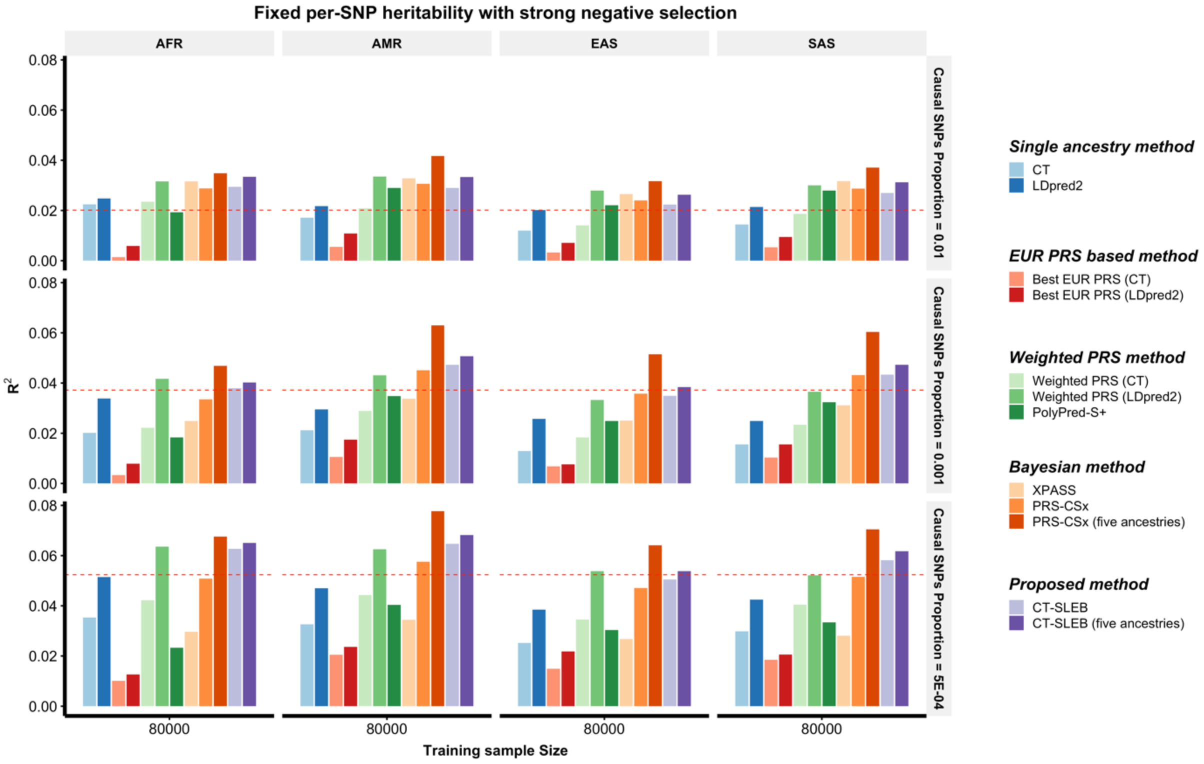

**Supplementary Figure 4 continued – Panel d: Simulation with training sample size of 100,000 for each of the four non-EUR populations.**

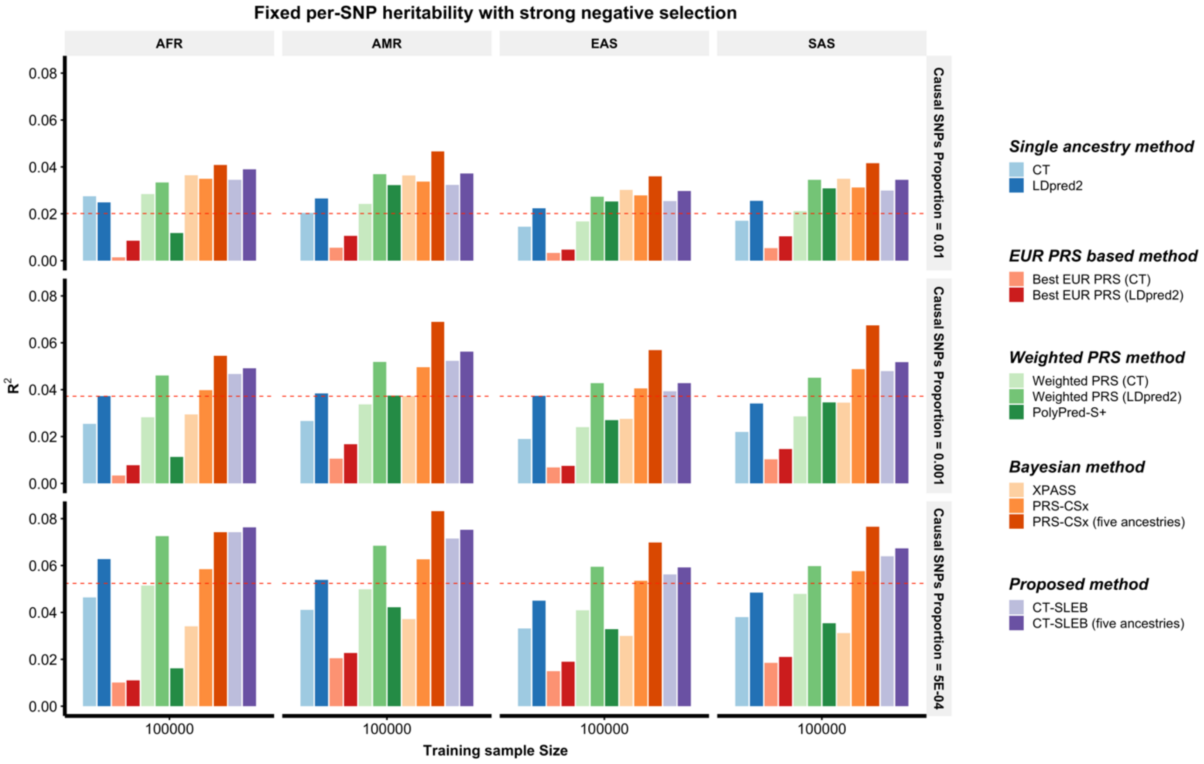

**Supplementary Figure 5: Simulation performances of different PRSs methods in multi-ancestry settings under a strong negative selection model with per-SNP heritability fixed across all populations (genetic correlation = 0.6).** Each of the four non-EUR populations has a training sample size of 15,000 (Sup. Fig. 6a), 45,000 (Sup. Fig. 6b), 80,000 (Sup. Fig. 6c) or 100,000 (Sup. Fig. 6d). For the EUR population, the size of the training sample is set at 100,000. The number of samples for each population in the tuning dataset is always 10,000. Prediction $R^2$ values are reported based on an independent validation dataset with 10,000 subjects for each population. Common SNP heritability is assumed to be 0.4 across all populations, and effect-size correlation is assumed to be 0.8 across all pairs of populations. The causal SNPs proportion (degree of polygenicity) is varied across 0.01 (top panel), 0.001 (medium panel), $5 \times 10^{-4}$ (bottom panel). All data are generated based on ~19 million common SNPs across the five populations, but analyses are restricted to ~2.0 million SNPs that are used on Hapmap3 + Multi-Ethnic Genotyping Arrays chip. PolyPred-S+ and PRS-CSx analysis is further restricted to ~1.3 million HM3 SNPs. All approaches are trained using data from the EUR and the target population. CT-SLEB and PRS-CSx are further evaluated using training data from five ancestries. The red dashed line shows the prediction performance of EUR PRS generated using single ancestry method (best of CT or LDpred2) in the EUR population.

**a)**

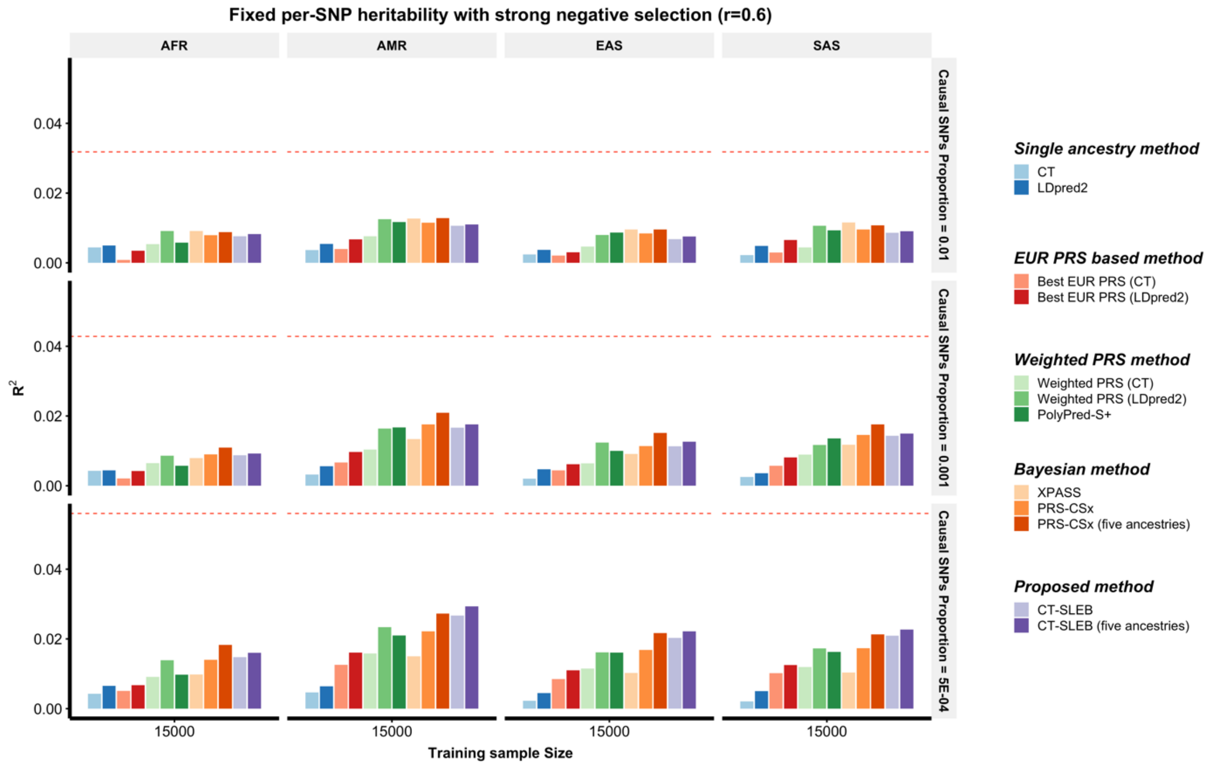

**Supplementary Figure 5 continued – Panel b: Simulation with training sample size of 45,000 for each of the four non-EUR populations.**

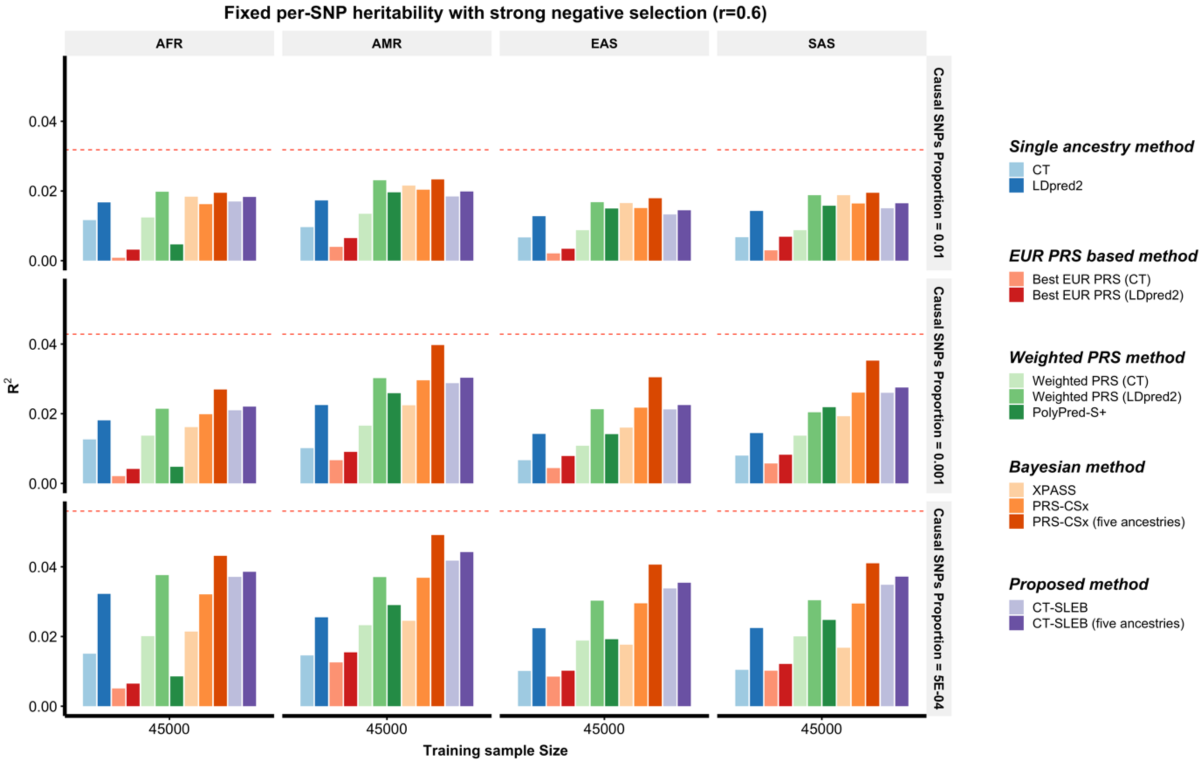

### Supplementary Figure 5 continued – Panel c: Simulation with training sample size of 80,000 for each of the four non-EUR populations

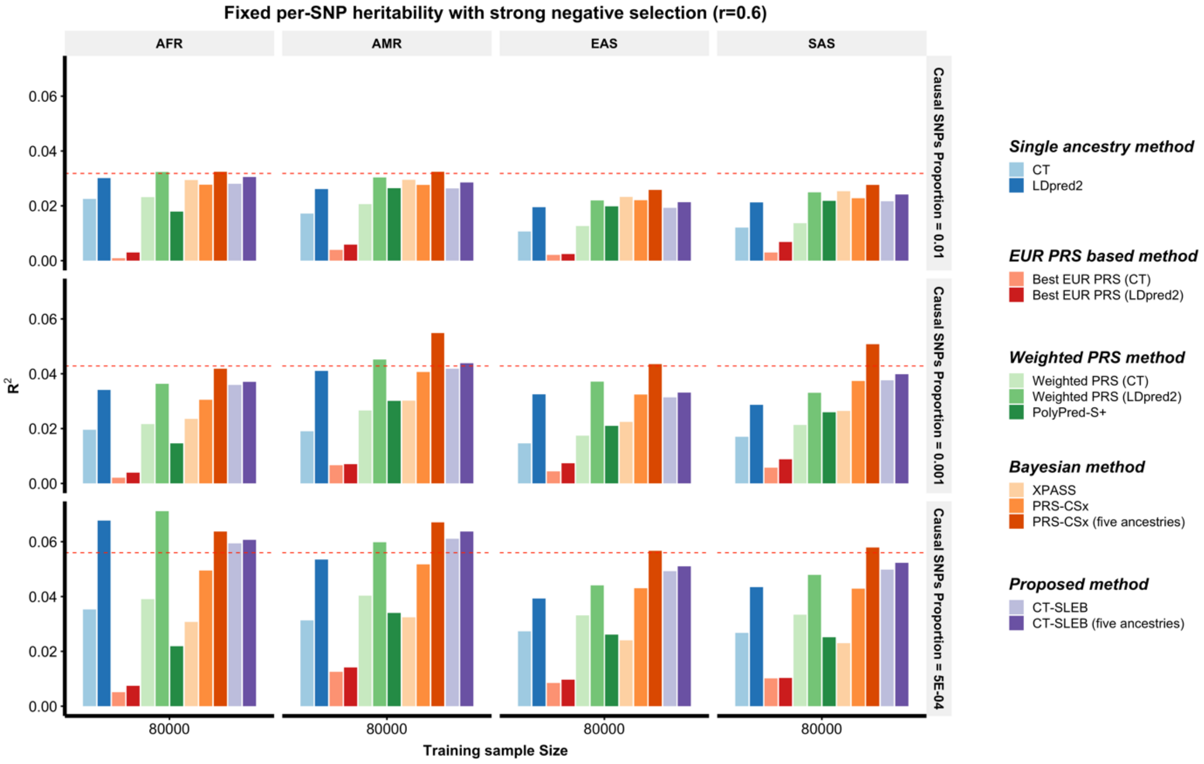

**Supplementary Figure 5 continued – Panel d: Simulation with training sample size of 100,000 for each of the four non-EUR populations.**

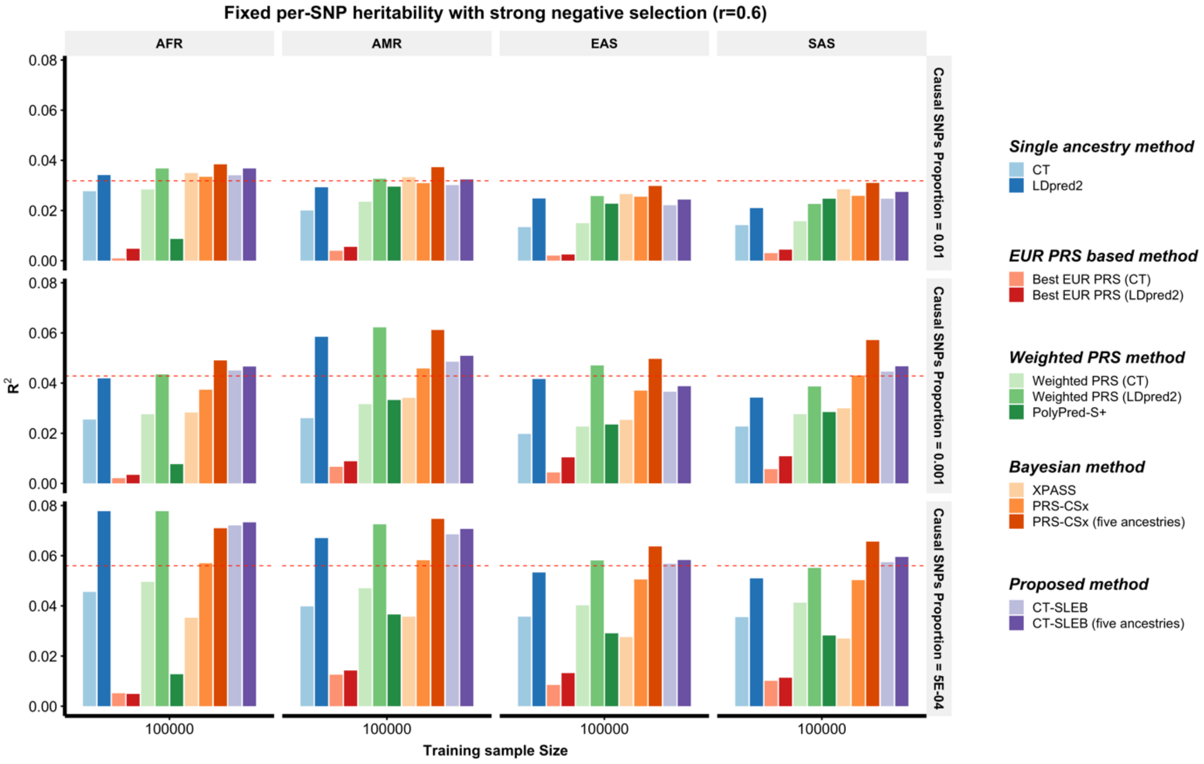

**Supplementary Figure 6: Prediction performance of CT-SLEB and PRS-CSx on HapMap3 array (HM3).** The training sample sizes are **15,000 (Sup. Fig. 6a)**, **45,000 (Sup. Fig. 6b), 80,000 (Sup. Fig. 8c) or 100,000** (**Sup. Fig. 8d**) for each of the four non-EUR populations. The training sample size for the EUR population is fixed at 100,000. Prediction $R^2$ values are reported based on independent validation dataset with 10,000 subjects for each population. <u>Common SNP heritability is assumed to be 0.</u>4 across all populations and effect-size correlation is assumed to be <u>0.8</u> across all pairs of populations. The causal SNPs proportion are varied across 0.01 (top panel), 0.001 (medium panel), $5 \times 10^{-4}$ (bottom panel) and effect sizes for causal variants are assumed to be related to allele frequency under <u>a strong negative selection model</u>.

**a)**

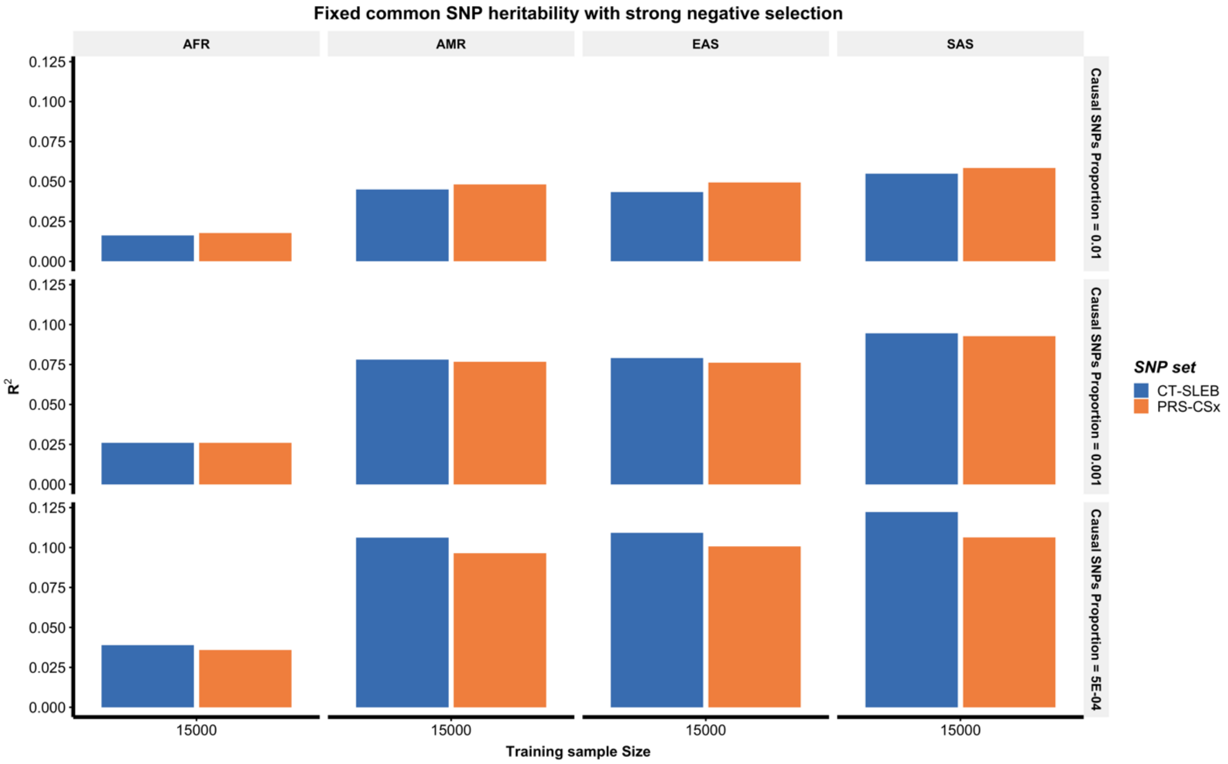

**Supplementary Figure 6 continued – Panel b: Simulation with training sample size of 45,000 for each of the four non-EUR populations.**

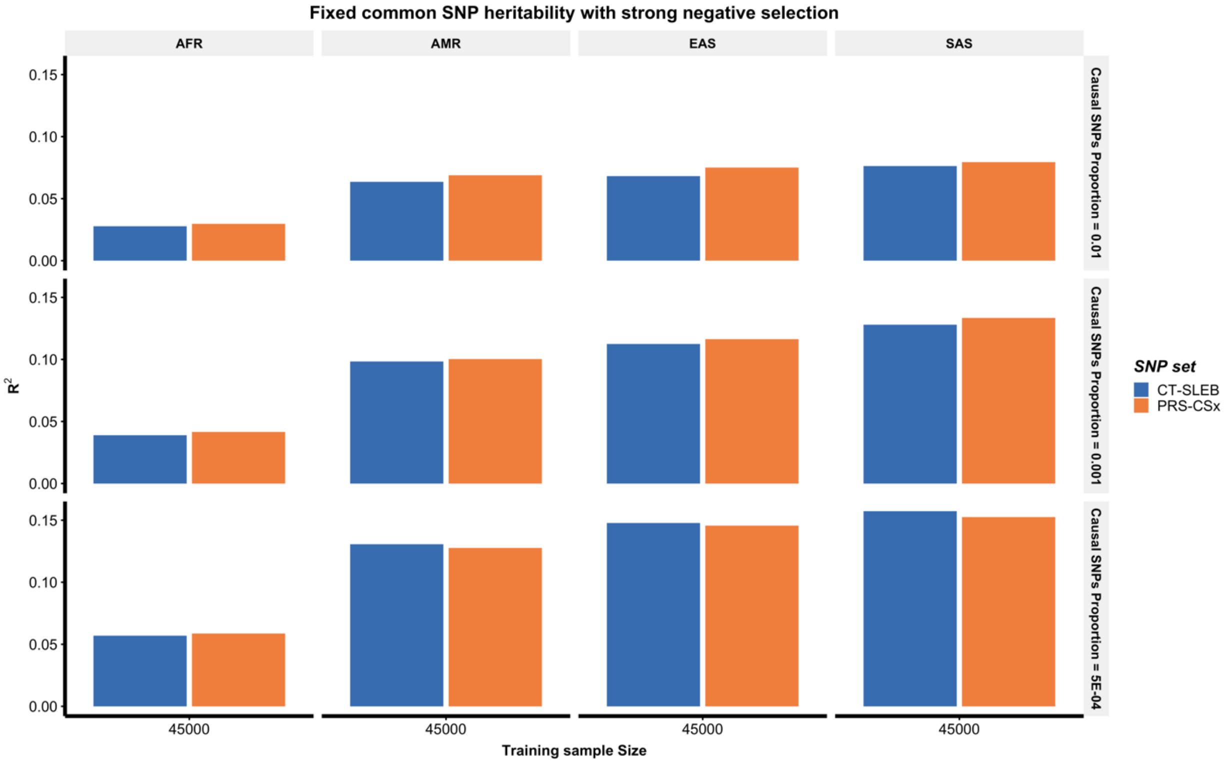

**Supplementary Figure 6 continued – Panel c: Simulation with training sample size of 80,000 for each of the four non-EUR populations.**

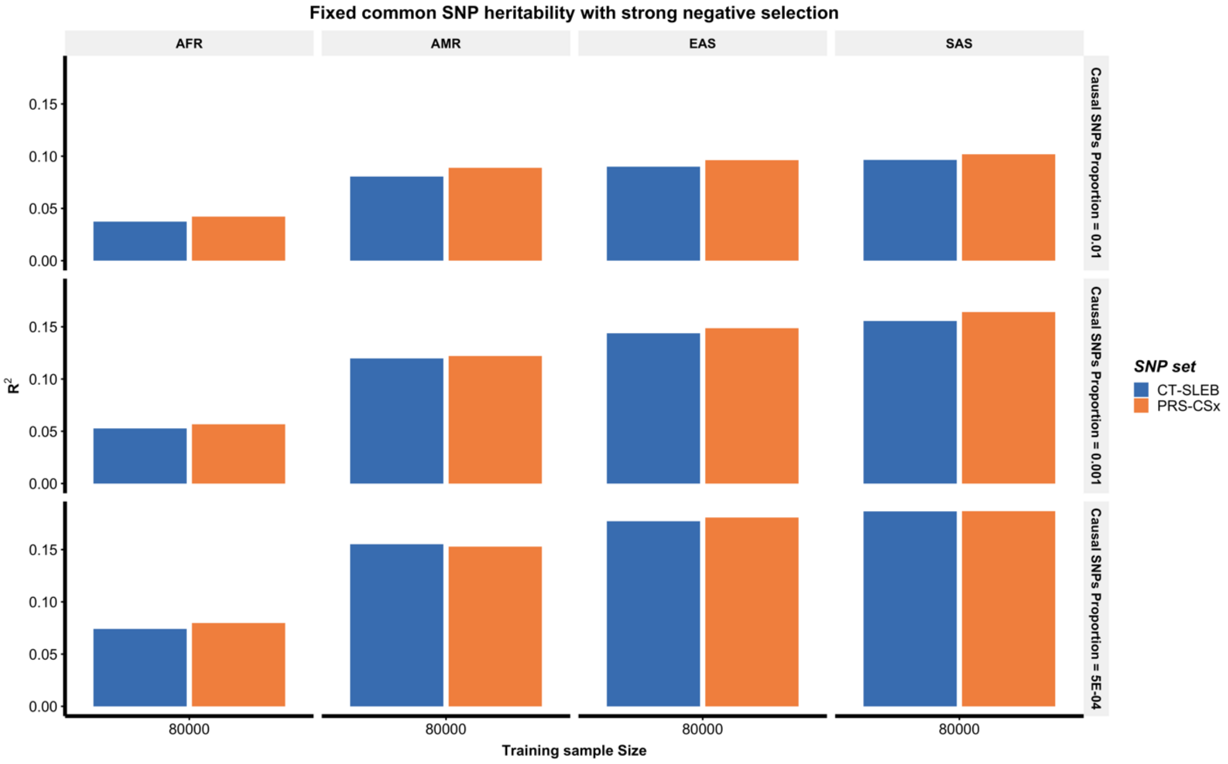

**Supplementary Figure 6 continued – Panel d: Simulation with training sample size of 100,000 for each of the four non-EUR populations.**

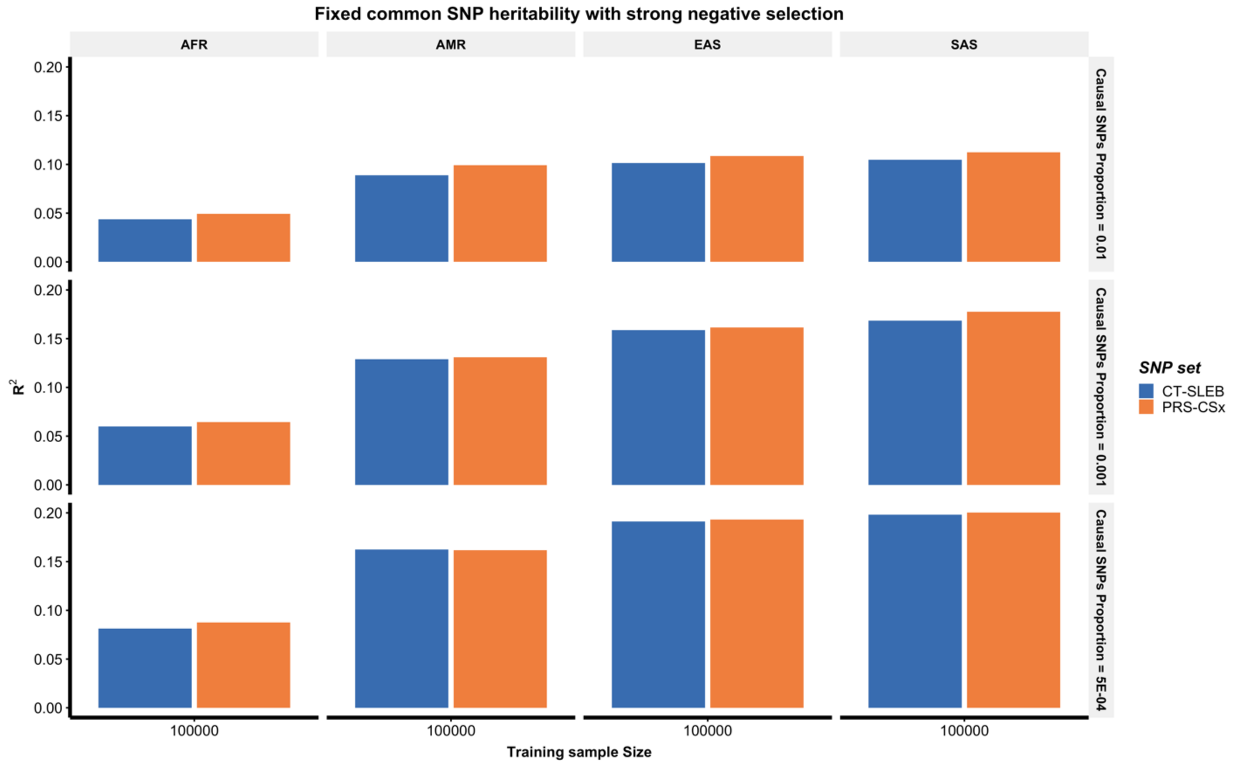

**Supplementary Figure 7: Prediction performance of CT-SLEB PRS across different ancestries relative to single ancestry EUR PRS in the EUR population.** The training sample size for the EUR population is fixed at 100,000, and PRS performance is assessed using single ancestry CT or LDpred2, whichever performs the best in each setting. Three different models for genetic architectures are considered: the common SNP heritability is fixed (at 0.4) with mild negative selection (**Sup. Fig. 7a**) and with no negative selection (**Sup. Fig. 7b**) and fixed per-SNP heritability with strong negative selection (**Sup. Fig. 7c**). The effect-size correlation is assumed to be 0.8 across all pairs of populations for settings in Sup. Fig. 7a and 7b. The effect-size correlation is assumed to be 0.6 for the setting in Sup. Fig. 7c.

a)

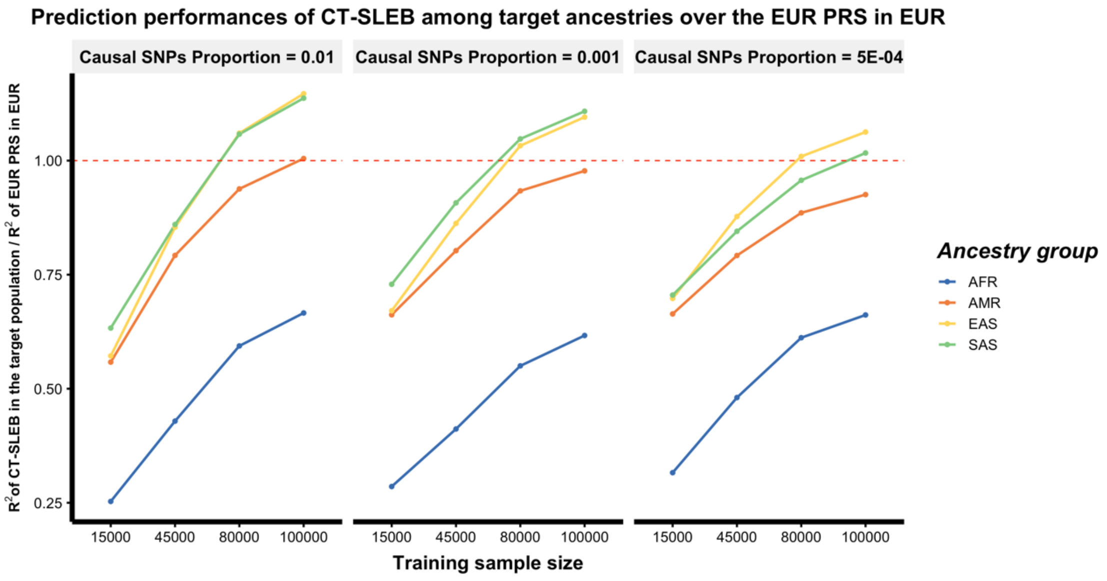


b)

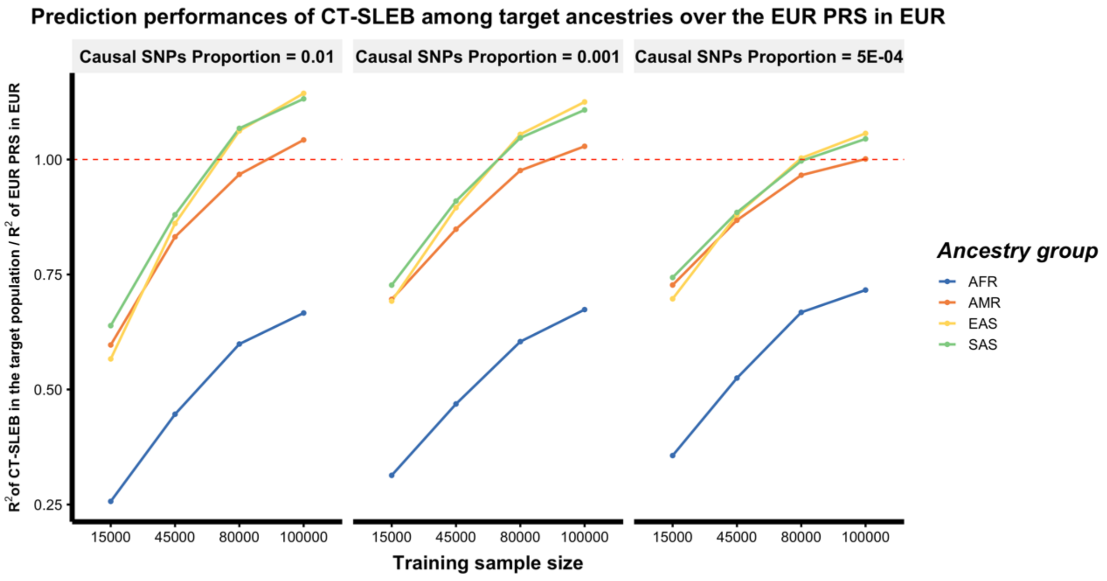

**Supplementary Figure 7 continued - Panel C: Prediction performance of CT-SLEB PRS with fixed per-SNP heritability, strong negative selection, and an effect-size correlation of 0.6.**

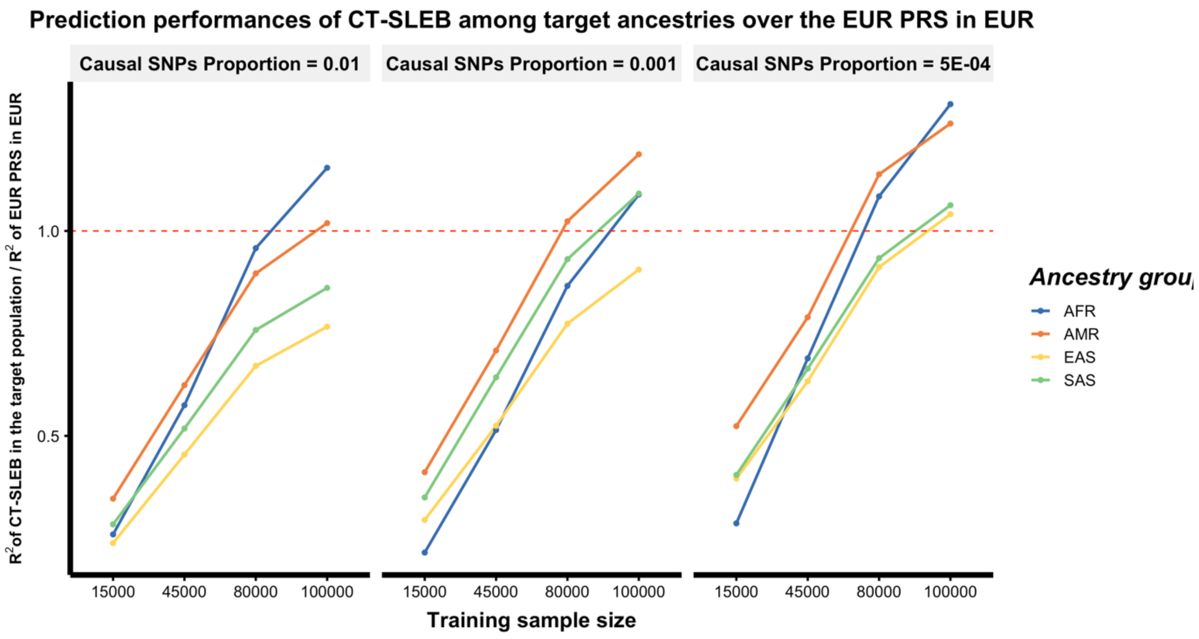

**Supplementary Figure 8: Prediction performance of CT-SLEB under different SNP density.** Analysis of each simulated data based on ~19 million SNPs are restricted to three different SNP sets Hapmap3 (~1.3 million SNPs), Hapmap3 + Multi-Ethnic Genotyping Arrays (~2.0 million SNPs), and 1000 Genomes Project (~19 million SNPs). The training sample size is **45,000 (Sup. Fig. 10a)** or **100,000 (Sup. Fig. 10b)** for each of the four non-EUR populations. The training sample size for the EUR population is fixed at 100,000. Prediction $R^2$ values are reported based on independent validation dataset with 10,000 subjects for each population. Common SNP heritability is assumed to be 0.4 across all populations and effect-size correlation is assumed to be 0.8 across all pairs of populations. The causal SNPs proportion are varied across 0.01 (top panel), 0.001 (medium panel), $5 \times 10^{-4}$ (bottom panel) and effect sizes for causal variants are assumed to be related to allele frequency under a strong negative selection model.

**a)**

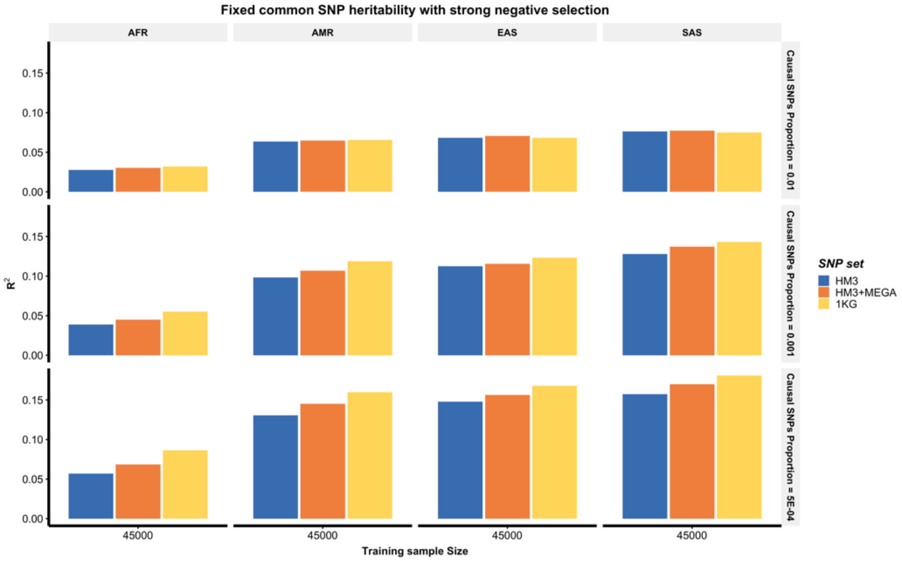


**b)**

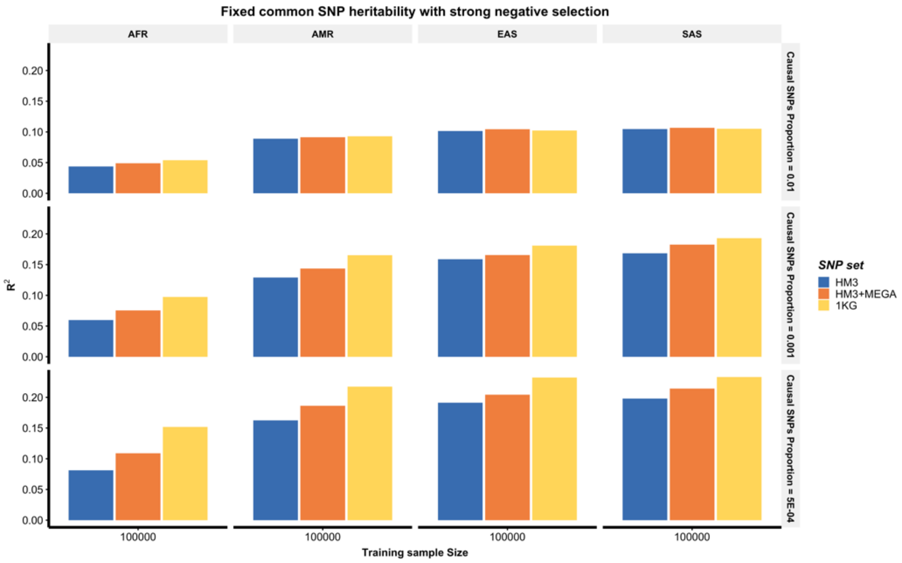

**Supplementary Figure 9:** Manhattan plot and QQ plot[1] based on the 23andMe, Inc. GWAS summary statistics for **any cardiovascular disease (any CVD)** in five populations: European, African American, Latino, East Asian, South Asian (from top panel to bottom panel). The red and blue shaded regions around the diagonal line in the QQ plots indicate the 95% confidence intervals expected under the null hypothesis of no association between genetic markers and the trait of interest, for minor allele frequencies (MAF) within the ranges (0.05, 0.5] and [0.01, 0.05], respectively. Under the null hypothesis, the p-value follows a uniform (0,1) distribution. The jth order statistic follows a Beta (j, N-j+1) distribution, where N is the total number of variants given a specific MAF cutoff.

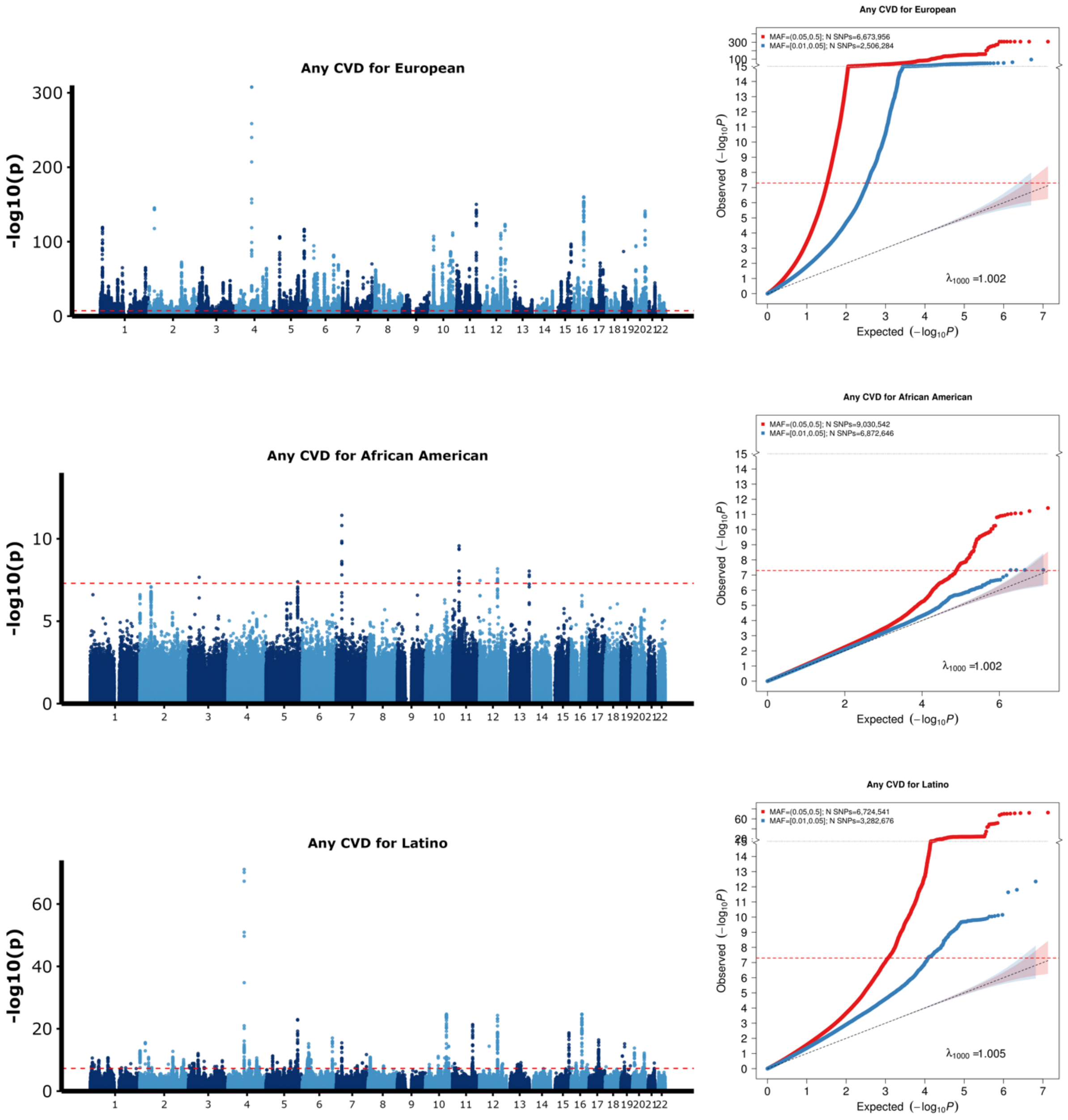


[1] For continuous traits, $\lambda_{1000}$ scales the genomic inflation factor $\lambda$ to a study with 1000 subjects using $\lambda_{1000} = 1 + 1000 * (\lambda - 1)/N$, where N is the total sample size. For binary traits, $\lambda_{1000}$ scales $\lambda$ to a study with 1000 cases and 1000 controls using $\lambda_{1000} = 1 + 1000 * (\lambda - 1) * (\frac{1}{N_{case}} + \frac{1}{N_{control}})$

**Supplementary Figure 9 continued: Manhattan and QQ plots for any CVD in the East Asian and South Asian populations.**

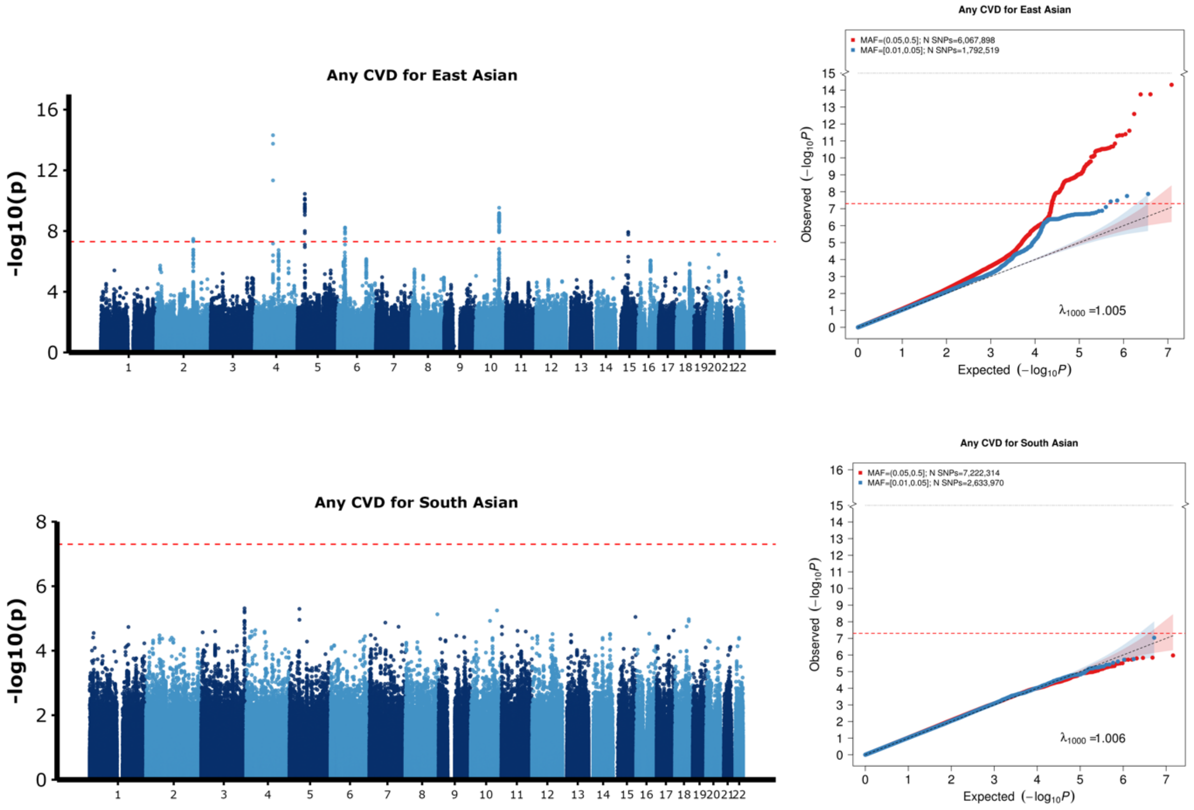


[1] For continuous traits, $\lambda_{1000}$ scales the genomic inflation factor $\lambda$ to a study with 1000 subjects using $\lambda_{1000} = 1 + 1000 * (\lambda - 1)/N$, where N is the total sample size. For binary traits, $\lambda_{1000}$ scales $\lambda$ to a study with 1000 cases and 1000 controls using $\lambda_{1000} = 1 + 1000 * (\lambda - 1) * (\frac{1}{N_{case}} + \frac{1}{N_{control}})$

**Supplementary Figure 10:** Manhattan plot and QQ plot[1] based on the 23andMe, Inc. GWAS summary statistics for **depression** in five populations: European, African American, Latino, East Asian, South Asian (from top panel to bottom panel). The red and blue shaded regions around the diagonal line in the QQ plots indicate the 95% confidence intervals expected under the null hypothesis of no association between genetic markers and the trait of interest, for minor allele frequencies (MAF) within the ranges (0.05, 0.5] and [0.01, 0.05], respectively. Under the null hypothesis, the p-value follows a uniform (0,1) distribution. The jth order statistic follows a Beta (j, N-j+1) distribution, where N is the total number of variants given a specific MAF cutoff.

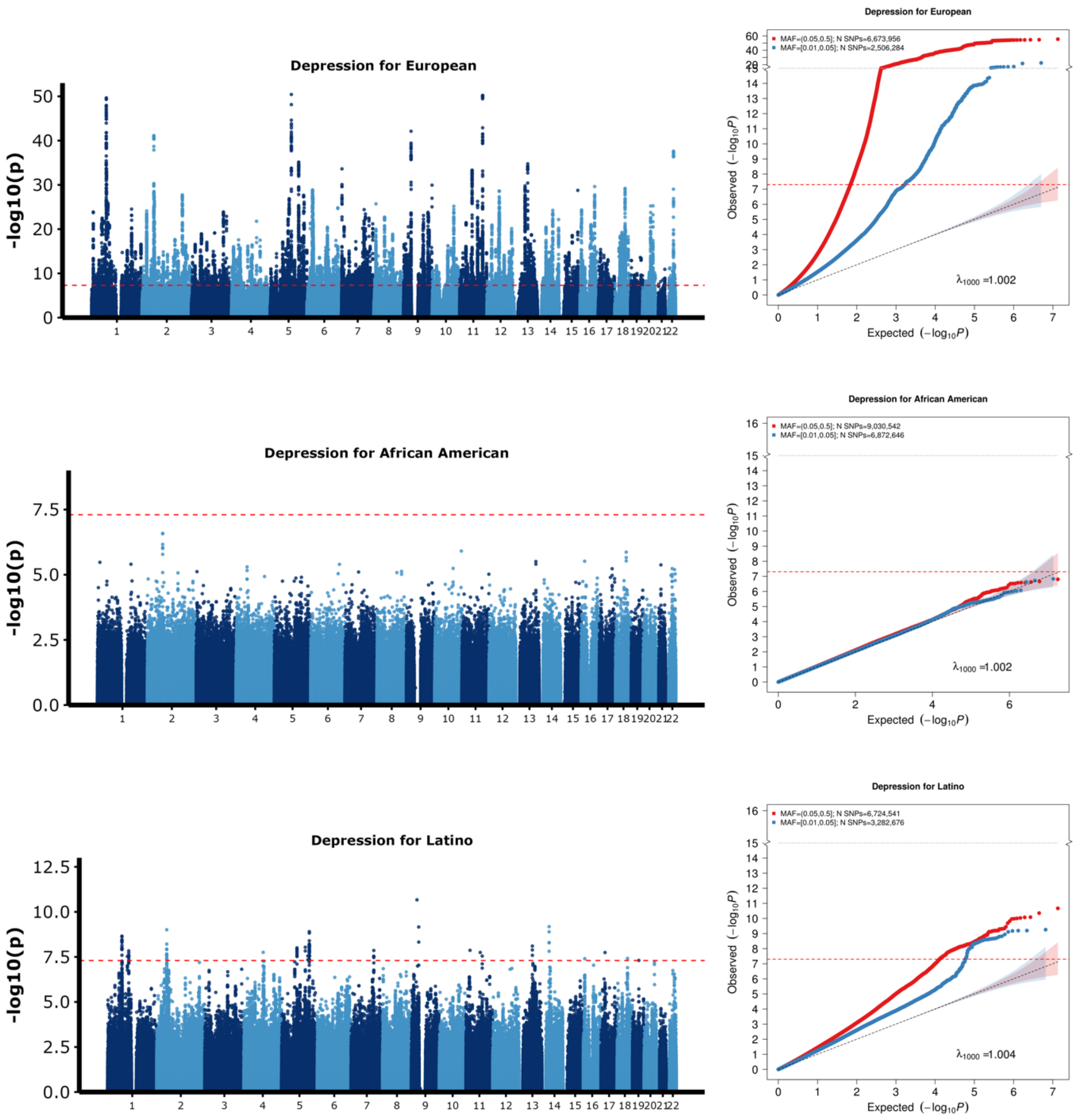


[1] For continuous traits, $\lambda_{1000}$ scales the genomic inflation factor $\lambda$ to a study with 1000 subjects using $\lambda_{1000} = 1 + 1000 * (\lambda - 1)/N$, where N is the total sample size. For binary traits, $\lambda_{1000}$ scales $\lambda$ to a study with 1000 cases and 1000 controls using $\lambda_{1000} = 1 + 1000 * (\lambda - 1) * (\frac{1}{N_{case}} + \frac{1}{N_{control}})$

**Supplementary Figure 10 continued: Manhattan and QQ plots for depression in the East Asian and South Asian populations.**

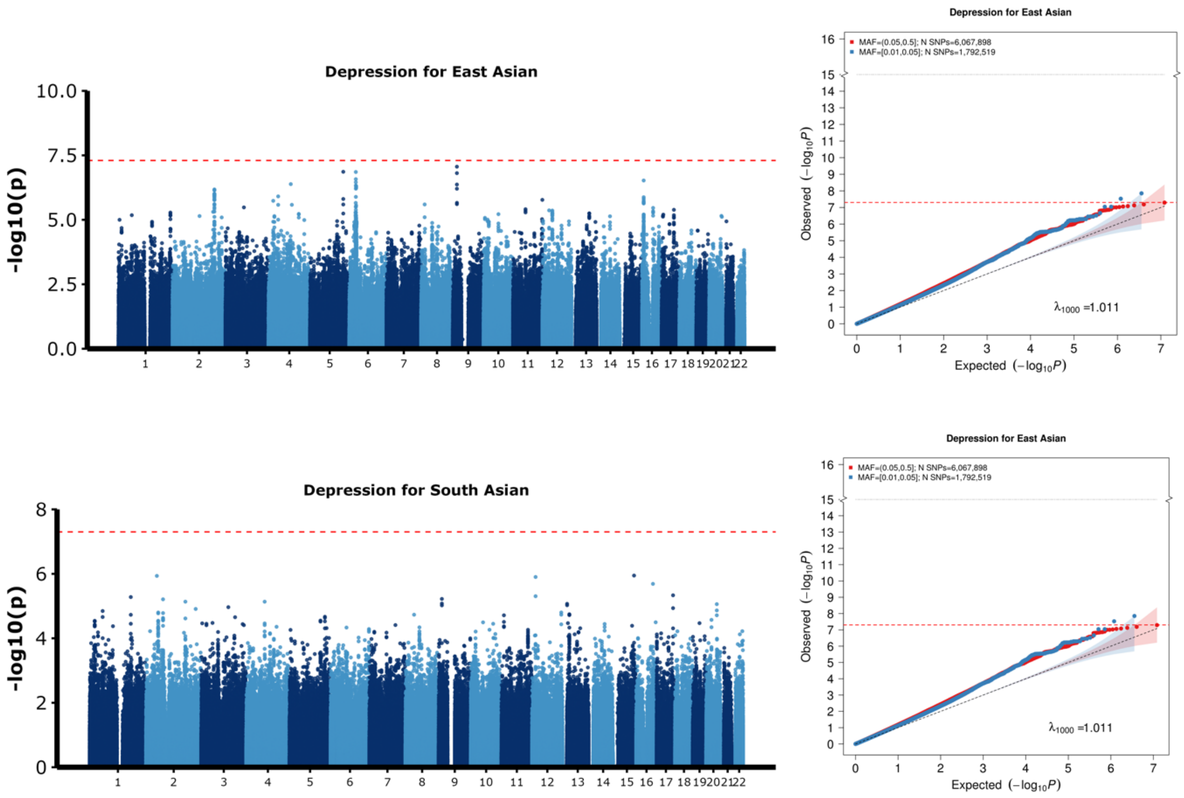


[1] For continuous traits, $\lambda_{1000}$ scales the genomic inflation factor $\lambda$ to a study with 1000 subjects using $\lambda_{1000} = 1 + 1000 * (\lambda - 1)/N$, where N is the total sample size. For binary traits, $\lambda_{1000}$ scales $\lambda$ to a study with 1000 cases and 1000 controls using $\lambda_{1000} = 1 + 1000 * (\lambda - 1) * (\frac{1}{N_{case}} + \frac{1}{N_{control}})$

**Supplementary Figure 11:** Manhattan plot and QQ plot[1] based on the 23andMe, Inc. GWAS summary statistics for **heart metabolic disease burden** in five populations: European, African American, Latino, East Asian, South Asian (from top panel to bottom panel). The red and blue shaded regions around the diagonal line in the QQ plots indicate the 95% confidence intervals expected under the null hypothesis of no association between genetic markers and the trait of interest, for minor allele frequencies (MAF) within the ranges (0.05, 0.5] and [0.01, 0.05], respectively. Under the null hypothesis, the p-value follows a uniform (0,1) distribution. The jth order statistic follows a Beta (j, N-j+1) distribution, where N is the total number of variants given a specific MAF cutoff.

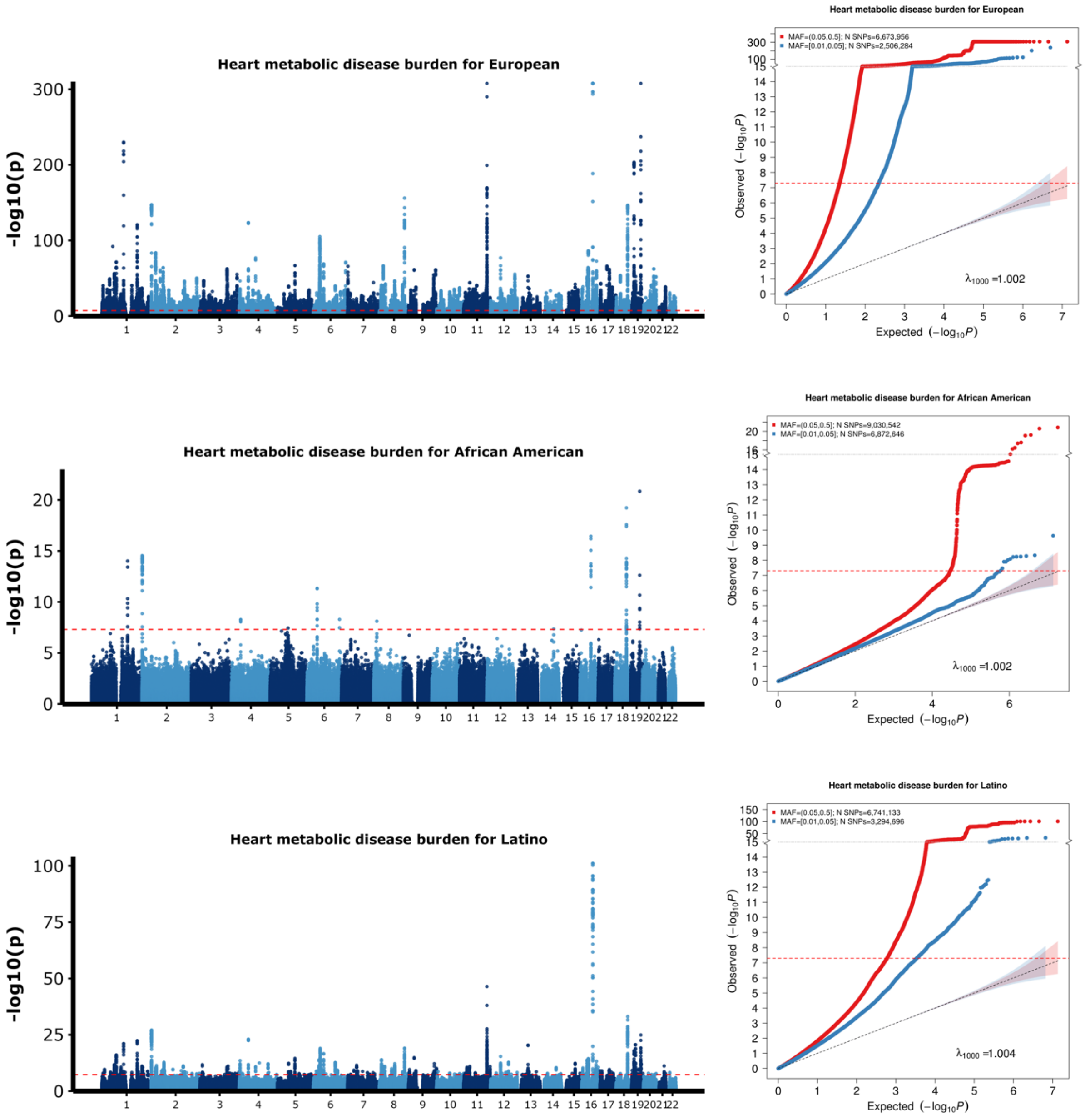


[1] For continuous traits, $\lambda_{1000}$ scales the genomic inflation factor $\lambda$ to a study with 1000 subjects using $\lambda_{1000} = 1 + 1000 * (\lambda - 1)/N$, where N is the total sample size. For binary traits, $\lambda_{1000}$ scales $\lambda$ to a study with 1000 cases and 1000 controls using $\lambda_{1000} = 1 + 1000 * (\lambda - 1) * (\frac{1}{N_{case}} + \frac{1}{N_{control}})$

**Supplementary Figure 11 continued: Manhattan and QQ plots for heart metabolic disease burden in the East Asian and South Asian populations.**

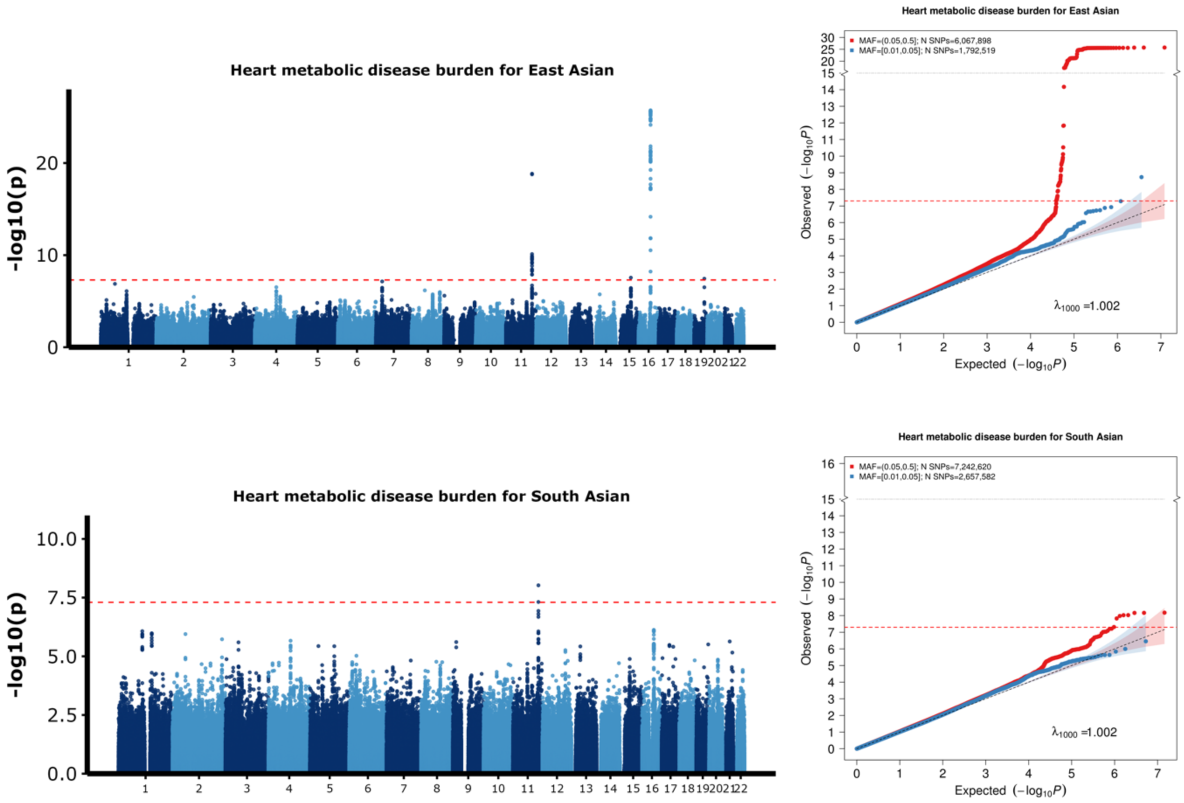


[1] For continuous traits, $\lambda_{1000}$ scales the genomic inflation factor $\lambda$ to a study with 1000 subjects using $\lambda_{1000} = 1 + 1000 * (\lambda - 1)/N$, where N is the total sample size. For binary traits, $\lambda_{1000}$ scales $\lambda$ to a study with 1000 cases and 1000 controls using $\lambda_{1000} = 1 + 1000 * (\lambda - 1) * (\frac{1}{N_{case}} + \frac{1}{N_{control}})$

**Supplementary Figure 12:** Manhattan plot and QQ plot[1] based on the 23andMe, Inc. GWAS summary statistics for **height** in five populations: European, African American, Latino, East Asian, South Asian (from top panel to bottom panel). The red and blue shaded regions around the diagonal line in the QQ plots indicate the 95% confidence intervals expected under the null hypothesis of no association between genetic markers and the trait of interest, for minor allele frequencies (MAF) within the ranges (0.05, 0.5] and [0.01, 0.05], respectively. Under the null hypothesis, the p-value follows a uniform (0,1) distribution. The jth order statistic follows a Beta (j, N-j+1) distribution, where N is the total number of variants given a specific MAF cutoff.

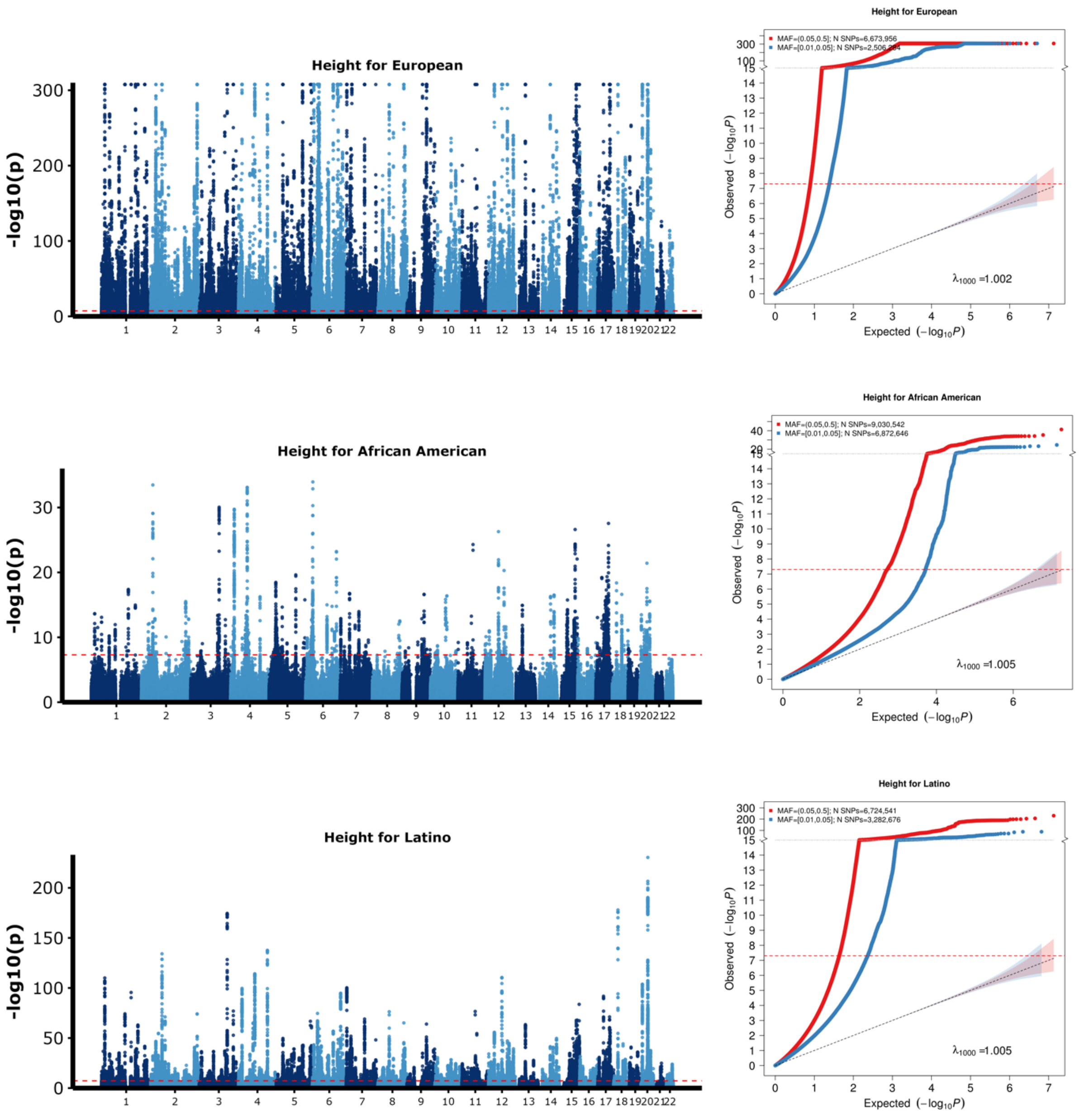


**1** For continuous traits, $\lambda_{1000}$ scales the genomic inflation factor $\lambda$ to a study with 1000 subjects using $\lambda_{1000} = 1 + 1000 * (\lambda - 1)/N$, where N is the total sample size. For binary traits, $\lambda_{1000}$ scales $\lambda$ to a study with 1000 cases and 1000 controls using $\lambda_{1000} = 1 + 1000 * (\lambda - 1) * (\frac{1}{N_{case}} + \frac{1}{N_{control}})$

**Supplementary Figure 12 continued: Manhattan and QQ plots for height in the East Asian and South Asian populations.**

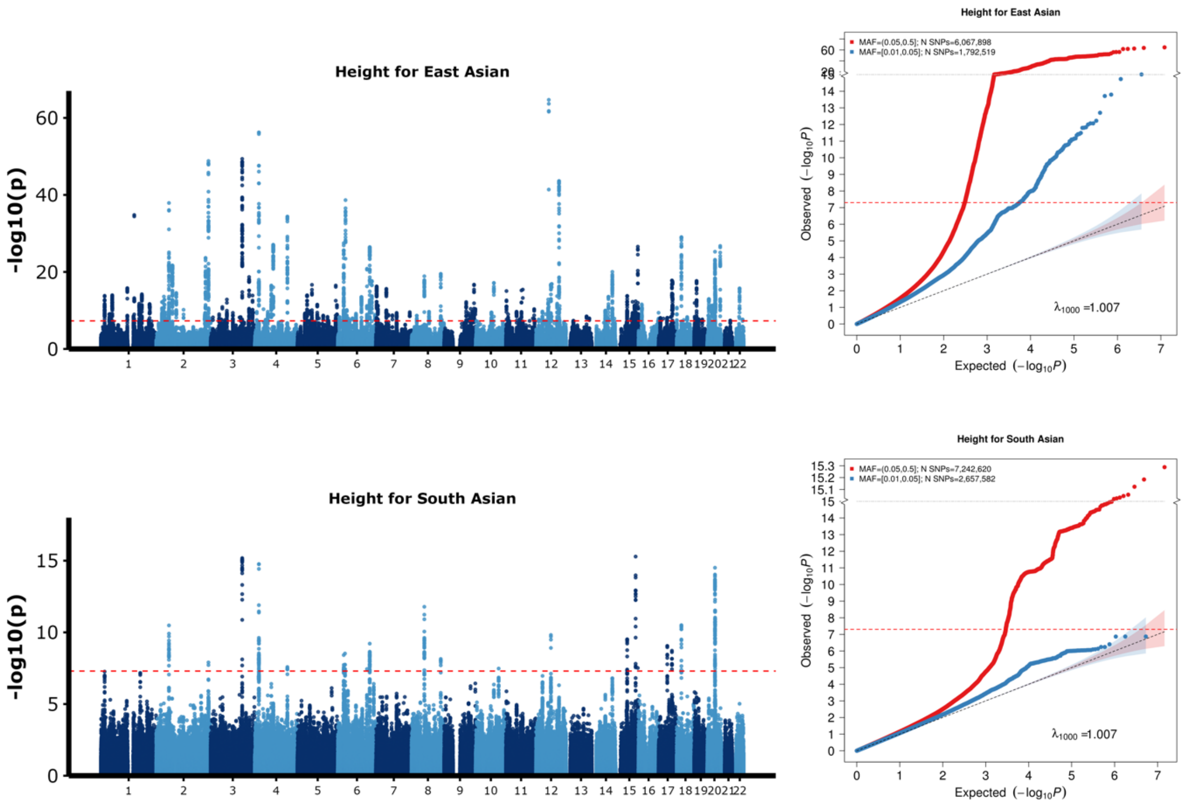


[1] For continuous traits, $\lambda_{1000}$ scales the genomic inflation factor $\lambda$ to a study with 1000 subjects using $\lambda_{1000} = 1 + 1000 * (\lambda - 1)/N$, where N is the total sample size. For binary traits, $\lambda_{1000}$ scales $\lambda$ to a study with 1000 cases and 1000 controls using $\lambda_{1000} = 1 + 1000 * (\lambda - 1) * (\frac{1}{N_{case}} + \frac{1}{N_{control}})$

**Supplementary Figure 13:** Manhattan plot and QQ plot[1] based on the 23andMe, Inc. GWAS summary statistics for **sing back musical note (SBMN)** in five populations: European, African American, Latino, East Asian, South Asian (from top panel to bottom panel). The red and blue shaded regions around the diagonal line in the QQ plots indicate the 95% confidence intervals expected under the null hypothesis of no association between genetic markers and the trait of interest, for minor allele frequencies (MAF) within the ranges (0.05, 0.5] and [0.01, 0.05], respectively. Under the null hypothesis, the p-value follows a uniform (0,1) distribution. The jth order statistic follows a Beta (j, N-j+1) distribution, where N is the total number of variants given a specific MAF cutoff.

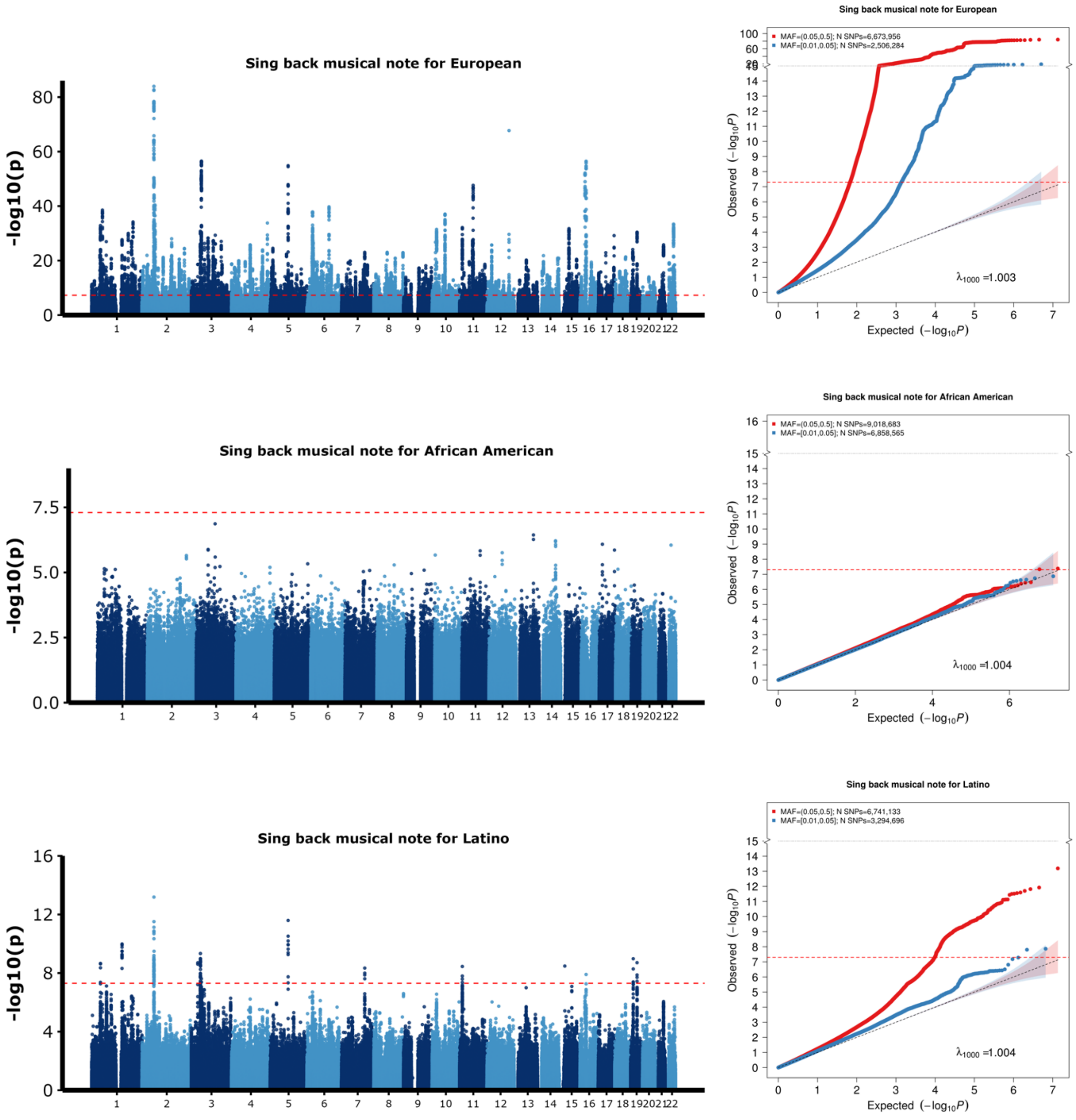


[1] For continuous traits, $\lambda_{1000}$ scales the genomic inflation factor $\lambda$ to a study with 1000 subjects using $\lambda_{1000} = 1 + 1000 * (\lambda - 1)/N$, where N is the total sample size. For binary traits, $\lambda_{1000}$ scales $\lambda$ to a study with 1000 cases and 1000 controls using $\lambda_{1000} = 1 + 1000 * (\lambda - 1) * (\frac{1}{N_{case}} + \frac{1}{N_{control}})$

**Supplementary Figure 13 continued: Manhattan and QQ plots for sing back musical note in the East Asian and South Asian populations.**

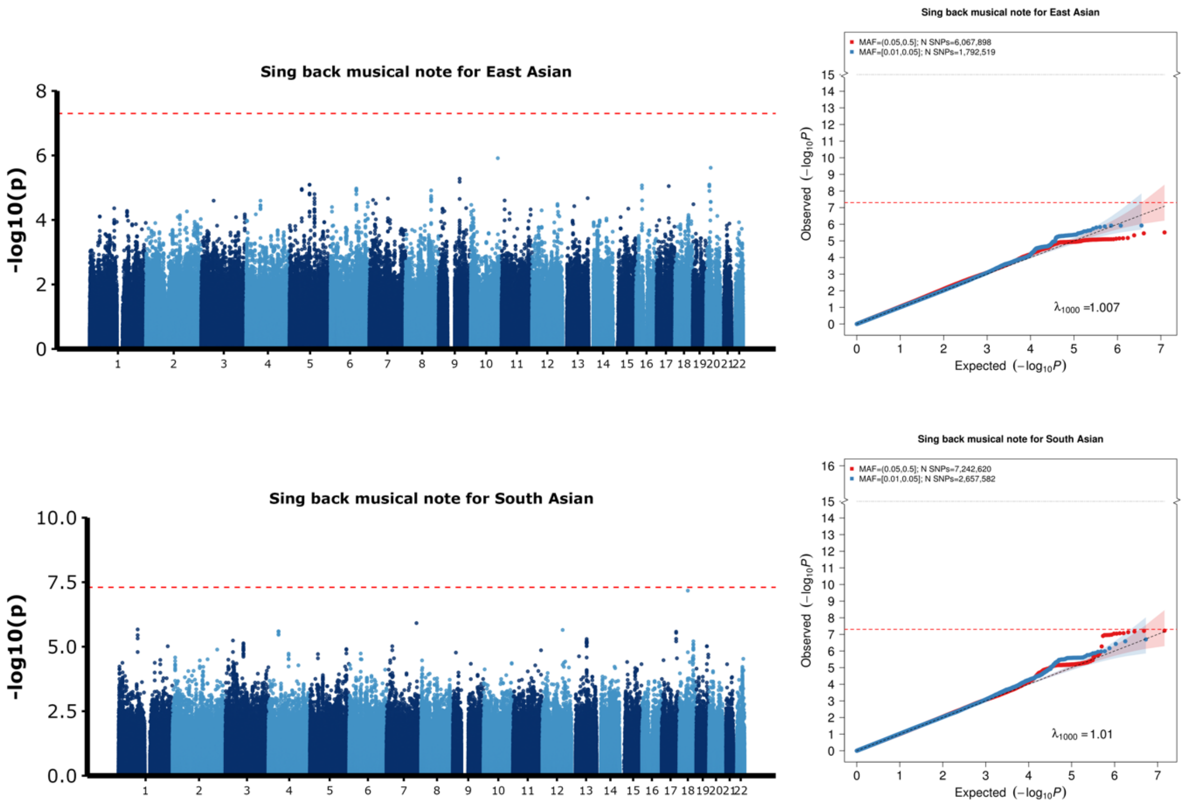


[1] For continuous traits, $\lambda_{1000}$ scales the genomic inflation factor $\lambda$ to a study with 1000 subjects using $\lambda_{1000} = 1 + 1000 * (\lambda - 1)/N$, where N is the total sample size. For binary traits, $\lambda_{1000}$ scales $\lambda$ to a study with 1000 cases and 1000 controls using $\lambda_{1000} = 1 + 1000 * (\lambda - 1) * (\frac{1}{N_{case}} + \frac{1}{N_{control}})$

**Supplementary Figure 14:** Manhattan plot and QQ plot[1] based on the 23andMe, Inc. GWAS summary statistics for **migraine diagnosis** in five populations: European, African American, Latino, East Asian, South Asian (from top panel to bottom panel). The red and blue shaded regions around the diagonal line in the QQ plots indicate the 95% confidence intervals expected under the null hypothesis of no association between genetic markers and the trait of interest, for minor allele frequencies (MAF) within the ranges (0.05, 0.5] and [0.01, 0.05], respectively. Under the null hypothesis, the p-value follows a uniform (0,1) distribution. The jth order statistic follows a Beta (j, N-j+1) distribution, where N is the total number of variants given a specific MAF cutoff.

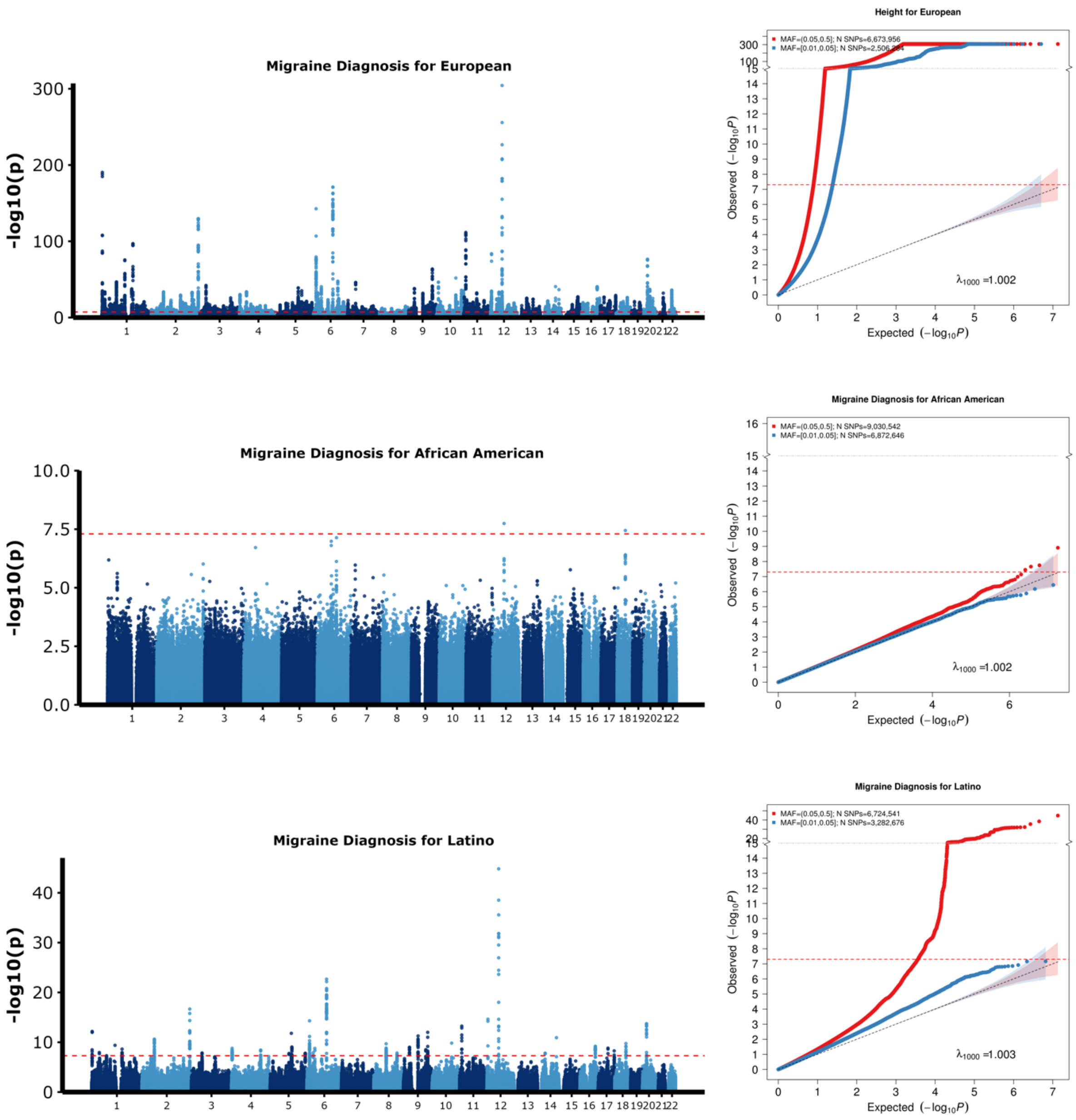


[1] For continuous traits, $\lambda_{1000}$ scales the genomic inflation factor $\lambda$ to a study with 1000 subjects using $\lambda_{1000} = 1 + 1000 * (\lambda - 1)/N$, where N is the total sample size. For binary traits, $\lambda_{1000}$ scales $\lambda$ to a study with 1000 cases and 1000 controls using $\lambda_{1000} = 1 + 1000 * (\lambda - 1) * (\frac{1}{N_{case}} + \frac{1}{N_{control}})$

**Supplementary Figure 14 continued: Manhattan and QQ plots for migraine diagnosis in the East Asian and South Asian populations.**

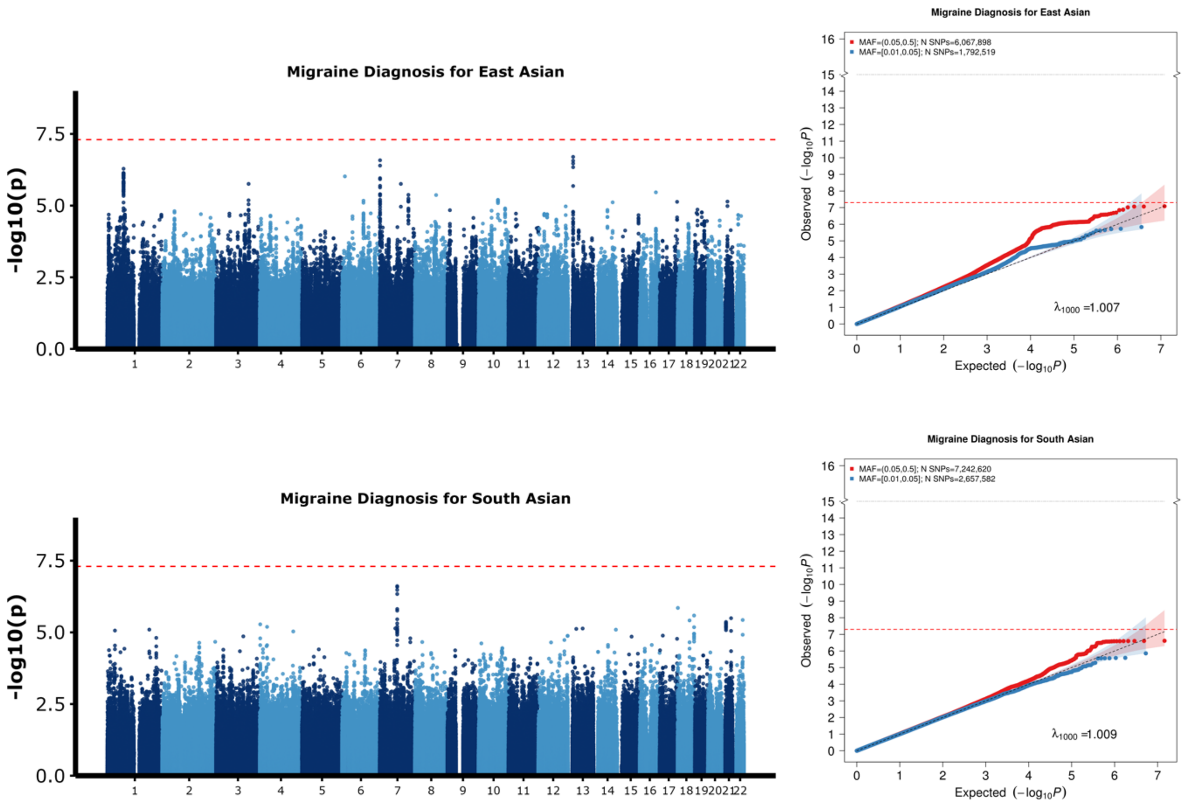


[1] For continuous traits, $\lambda_{1000}$ scales the genomic inflation factor $\lambda$ to a study with 1000 subjects using $\lambda_{1000} = 1 + 1000 * (\lambda - 1)/N$, where N is the total sample size. For binary traits, $\lambda_{1000}$ scales $\lambda$ to a study with 1000 cases and 1000 controls using $\lambda_{1000} = 1 + 1000 * (\lambda - 1) * (\frac{1}{N_{case}} + \frac{1}{N_{control}})$

**Supplementary Figure 15:** Manhattan plot and QQ plot[1] based on the 23andMe, Inc. GWAS summary statistics for **morning person** in five populations: European, African American, Latino, East Asian, South Asian (from top panel to bottom panel). The red and blue shaded regions around the diagonal line in the QQ plots indicate the 95% confidence intervals expected under the null hypothesis of no association between genetic markers and the trait of interest, for minor allele frequencies (MAF) within the ranges (0.05, 0.5] and [0.01, 0.05], respectively. Under the null hypothesis, the p-value follows a uniform (0,1) distribution. The jth order statistic follows a Beta (j, N-j+1) distribution, where N is the total number of variants given a specific MAF cutoff.

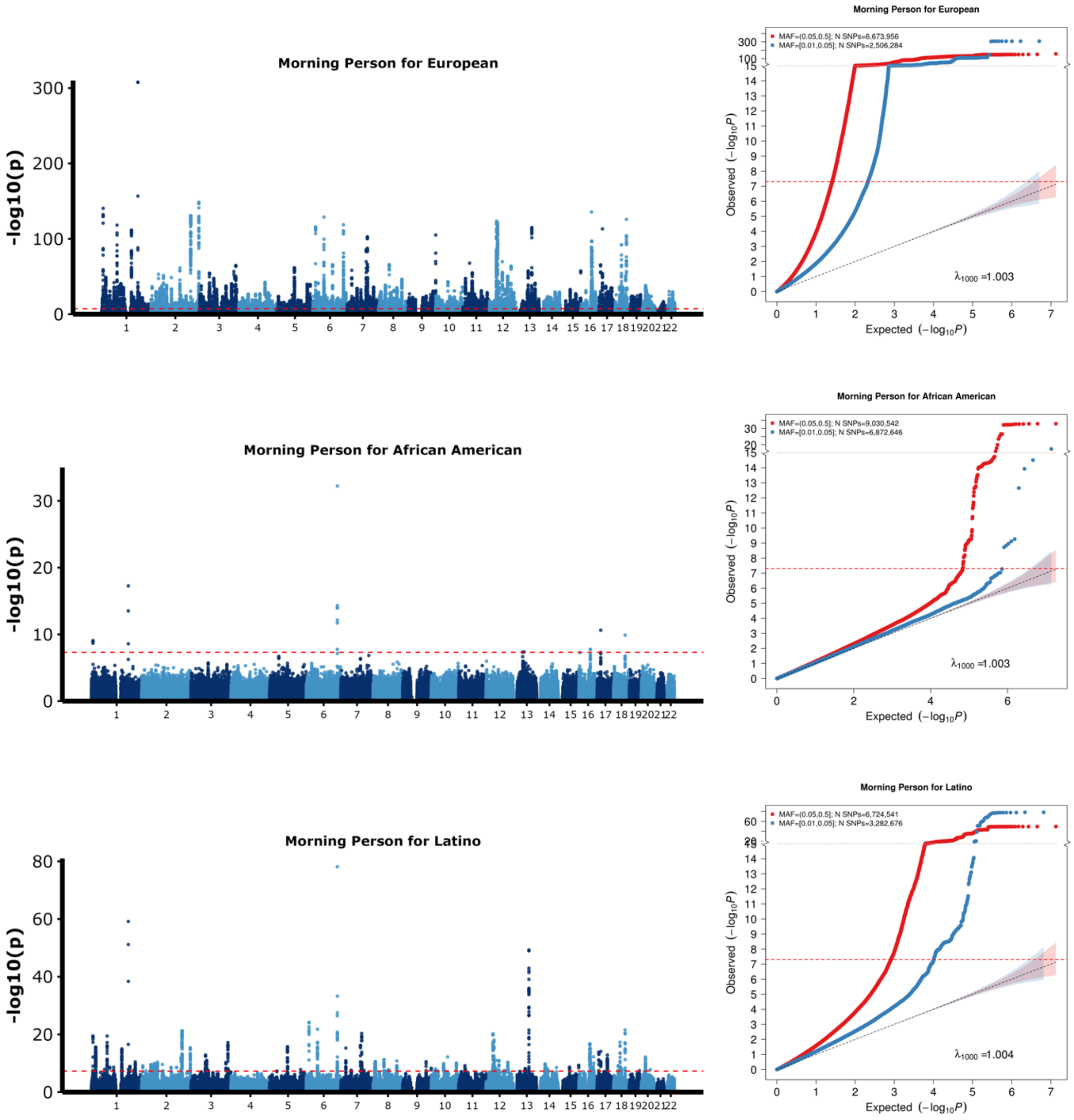


[1] For continuous traits, $\lambda_{1000}$ scales the genomic inflation factor $\lambda$ to a study with 1000 subjects using $\lambda_{1000} = 1 + 1000 * (\lambda - 1)/N$, where N is the total sample size. For binary traits, $\lambda_{1000}$ scales $\lambda$ to a study with 1000 cases and 1000 controls using $\lambda_{1000} = 1 + 1000 * (\lambda - 1) * (\frac{1}{N_{case}} + \frac{1}{N_{control}})$

**Supplementary Figure 15 continued: Manhattan and QQ plots for morning person in the East Asian and South Asian populations.**

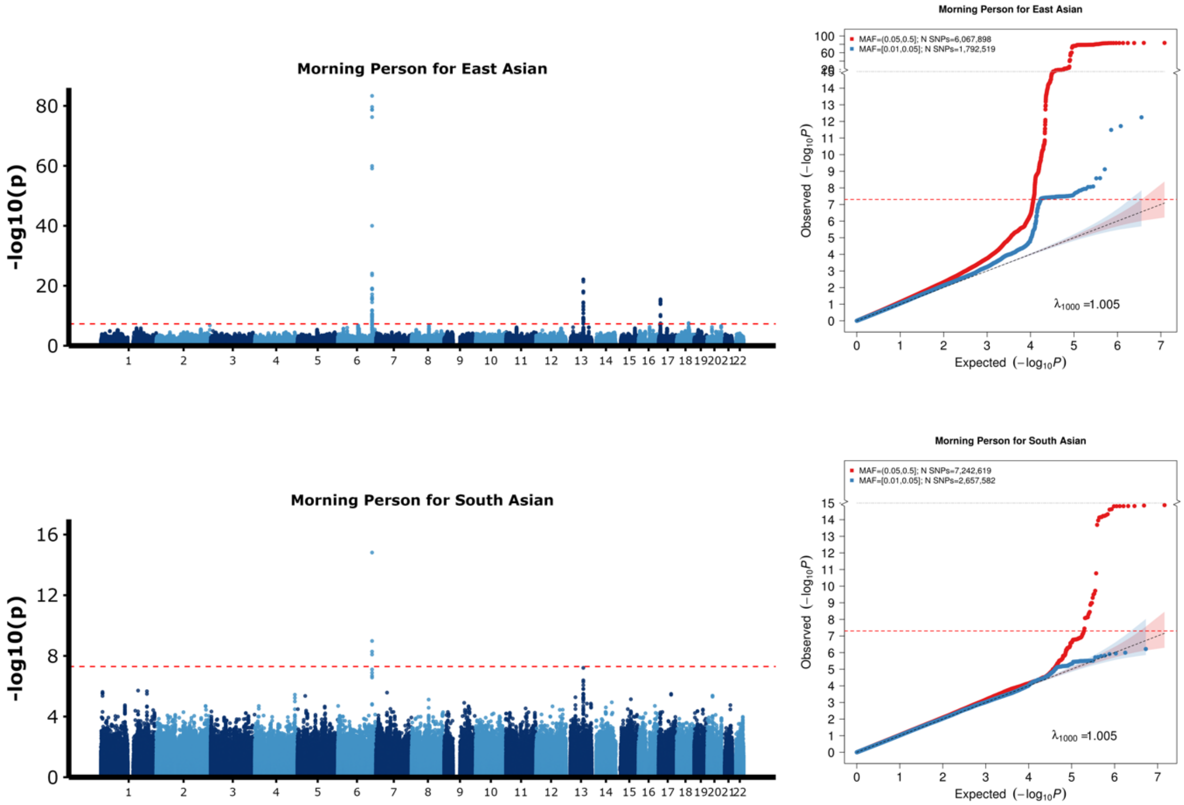


[1] For continuous traits, $\lambda_{1000}$ scales the genomic inflation factor $\lambda$ to a study with 1000 subjects using $\lambda_{1000} = 1 + 1000 * (\lambda - 1)/N$, where N is the total sample size. For binary traits, $\lambda_{1000}$ scales $\lambda$ to a study with 1000 cases and 1000 controls using $\lambda_{1000} = 1 + 1000 * (\lambda - 1) * (\frac{1}{N_{case}} + \frac{1}{N_{control}})$

**Supplementary Figure 16:** Manhattan plot and QQ plot[1] based on the Global Lipids Genetics Consortium (GLGC) GWAS summary statistics for **high-density lipoprotein cholesterol (HDL)** in five populations: European, African (primarily African American), Latino, East Asian, South Asian (from top panel to bottom panel). The red and blue shaded regions around the diagonal line in the QQ plots indicate the 95% confidence intervals expected under the null hypothesis of no association between genetic markers and the trait of interest, for minor allele frequencies (MAF) within the ranges (0.05, 0.5]and [0.01, 0.05], respectively. Under the null hypothesis, the p-value follows a uniform (0,1) distribution. The jth order statistic follows a Beta (j, N-j+1) distribution, where N is the total number of variants given a specific MAF cutoff

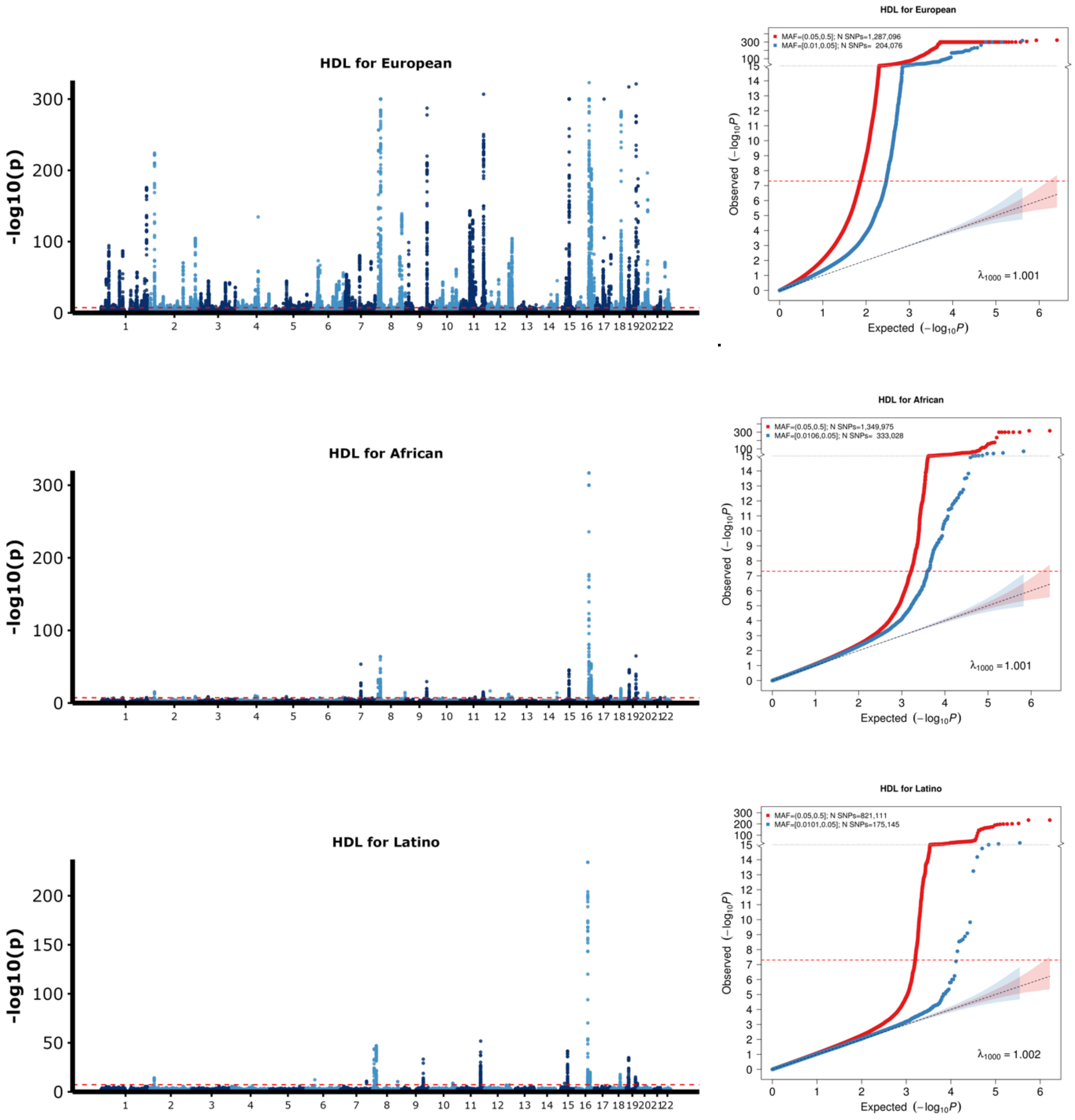


[1] For continuous traits, $\lambda_{1000}$ scales the genomic inflation factor $\lambda$ to a study with 1000 subjects using $\lambda_{1000} = 1 + 1000 * (\lambda - 1)/N$, where N is the total sample size. For binary traits, $\lambda_{1000}$ scales $\lambda$ to a study with 1000 cases and 1000 controls using $\lambda_{1000} = 1 + 1000 * (\lambda - 1) * (\frac{1}{N_{case}} + \frac{1}{N_{control}})$

**Supplementary Figure 16 continued: Manhattan and QQ plots for HDL in the East Asian and South Asian populations.**

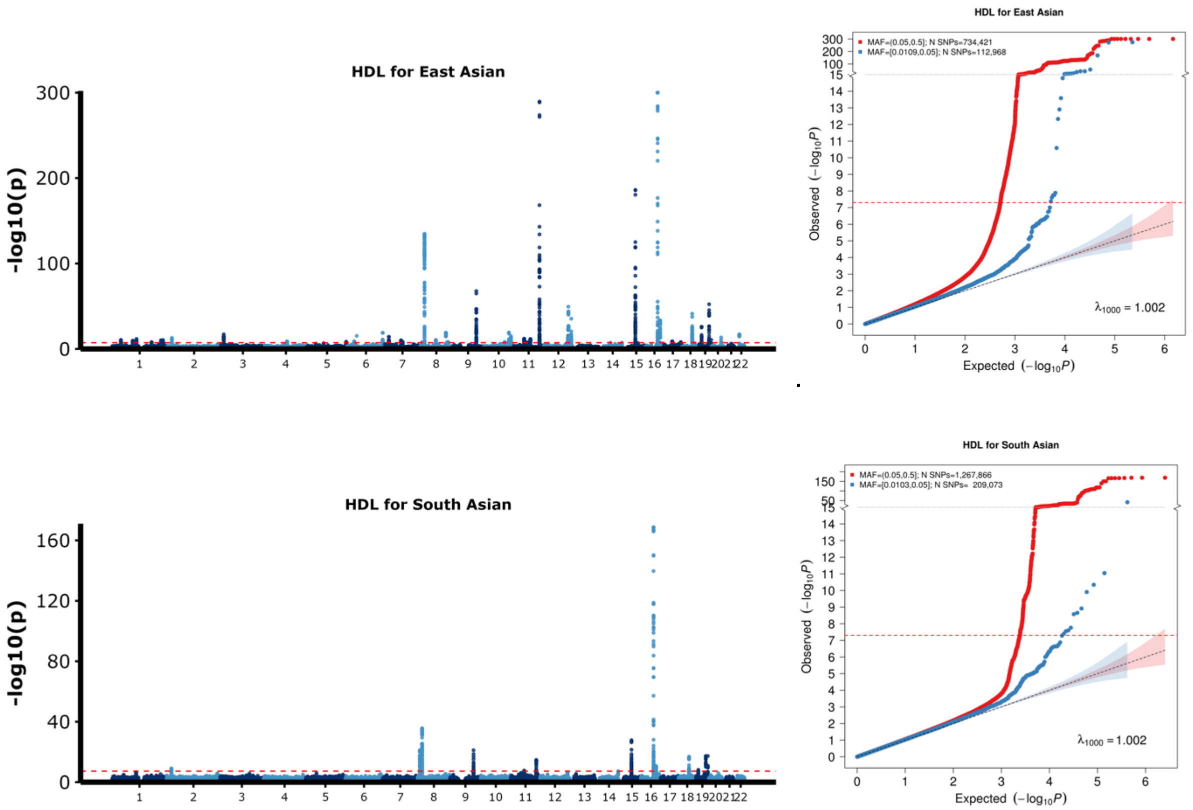


[1] For continuous traits, $\lambda_{1000}$ scales the genomic inflation factor $\lambda$ to a study with 1000 subjects using $\lambda_{1000} = 1 + 1000 * (\lambda - 1)/N$, where N is the total sample size. For binary traits, $\lambda_{1000}$ scales $\lambda$ to a study with 1000 cases and 1000 controls using $\lambda_{1000} = 1 + 1000 * (\lambda - 1) * (\frac{1}{N_{case}} + \frac{1}{N_{control}})$

**Supplementary Figure 17:** Manhattan plot and QQ plot[1] based on the Global Lipids Genetics Consortium (GLGC) GWAS summary statistics for **low-density lipoprotein cholesterol (LDL)** in five populations: European, African (primarily African American), Latino, East Asian, South Asian (from top panel to bottom panel). The red and blue shaded regions around the diagonal line in the QQ plots indicate the 95% confidence intervals expected under the null hypothesis of no association between genetic markers and the trait of interest, for minor allele frequencies (MAF) within the ranges (0.05, 0.5] and [0.01, 0.05], respectively. Under the null hypothesis, the p-value follows a uniform (0,1) distribution. The jth order statistic follows a Beta (j, N-j+1) distribution, where N is the total number of variants given a specific MAF cutoff.

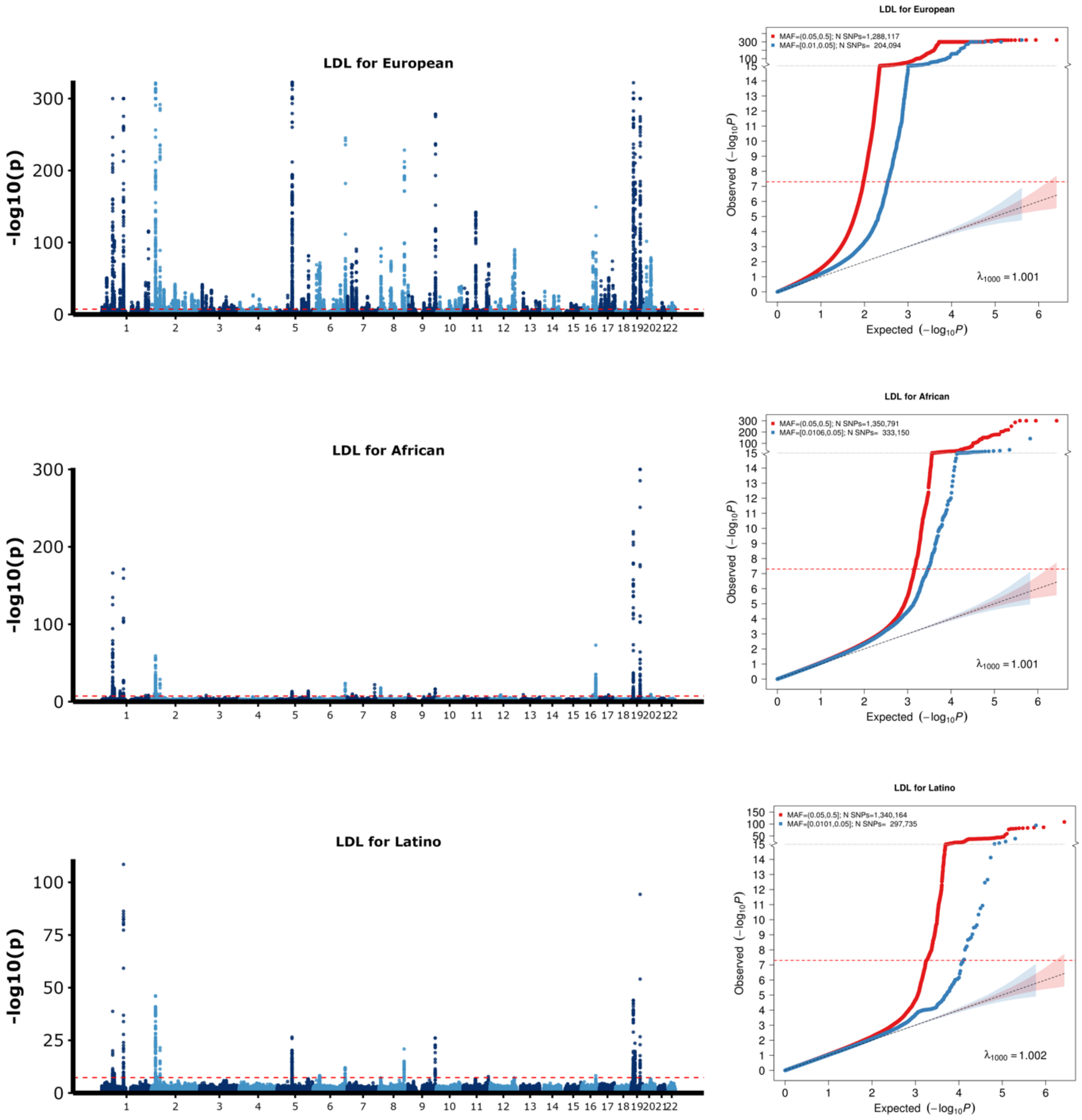


[1] For continuous traits, $\lambda_{1000}$ scales the genomic inflation factor $\lambda$ to a study with 1000 subjects using $\lambda_{1000} = 1 + 1000 * (\lambda - 1)/N$, where N is the total sample size. For binary traits, $\lambda_{1000}$ scales $\lambda$ to a study with 1000 cases and 1000 controls using $\lambda_{1000} = 1 + 1000 * (\lambda - 1) * (\frac{1}{N_{case}} + \frac{1}{N_{control}})$

**Supplementary Figure 17 continued: Manhattan and QQ plots for LDL in the East Asian and South Asian populations.**

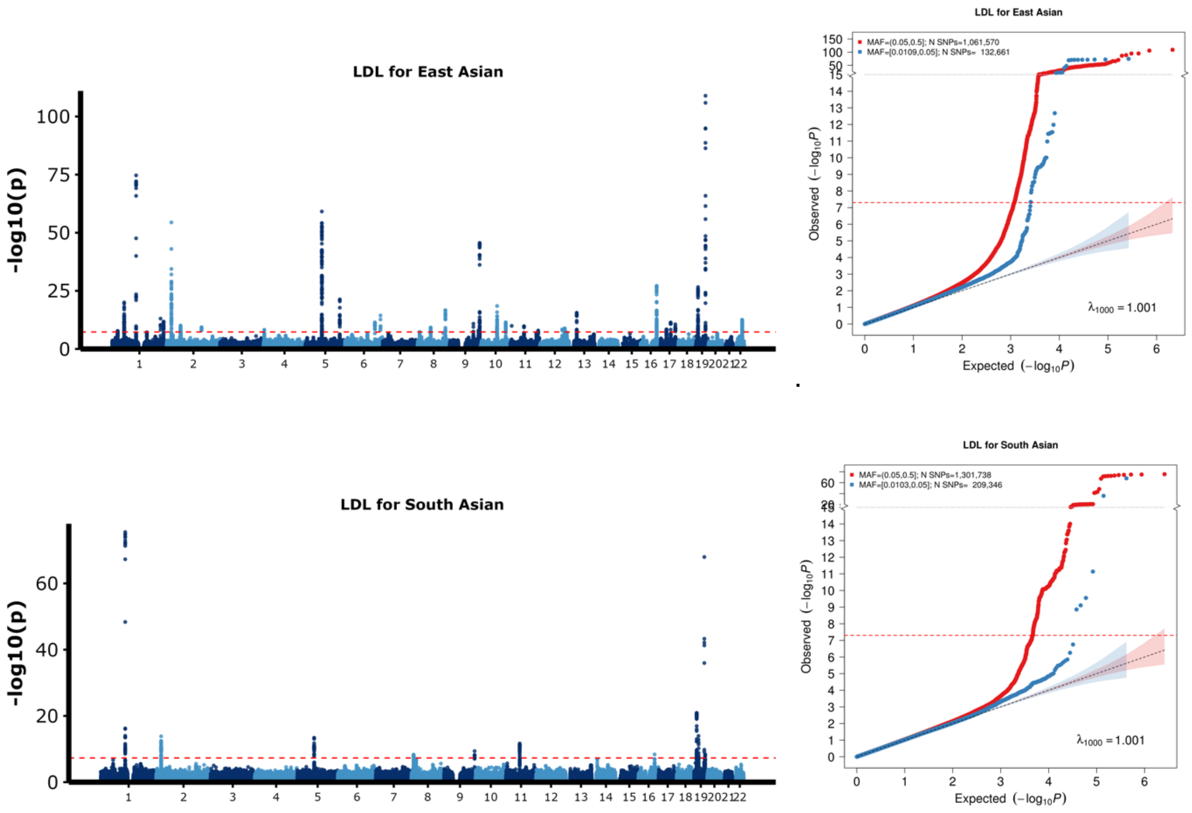


[1] For continuous traits, $\lambda_{1000}$ scales the genomic inflation factor $\lambda$ to a study with 1000 subjects using $\lambda_{1000} = 1 + 1000 * (\lambda - 1)/N$, where N is the total sample size. For binary traits, $\lambda_{1000}$ scales $\lambda$ to a study with 1000 cases and 1000 controls using $\lambda_{1000} = 1 + 1000 * (\lambda - 1) * (\frac{1}{N_{case}} + \frac{1}{N_{control}})$

**Supplementary Figure 18:** Manhattan plot and QQ plot[1] based on the Global Lipids Genetics Consortium (GLGC) GWAS summary statistics for **log triglycerides (logTG)** in five populations: European, African (primarily African American), Latino, East Asian, South Asian (from top panel to bottom panel). The red and blue shaded regions around the diagonal line in the QQ plots indicate the 95% confidence intervals expected under the null hypothesis of no association between genetic markers and the trait of interest, for minor allele frequencies (MAF) within the ranges (0.05, 0.5] and [0.01, 0.05], respectively. Under the null hypothesis, the p-value follows a uniform (0,1) distribution. The jth order statistic follows a Beta (j, N-j+1) distribution, where N is the total number of variants given a specific MAF cutoff.

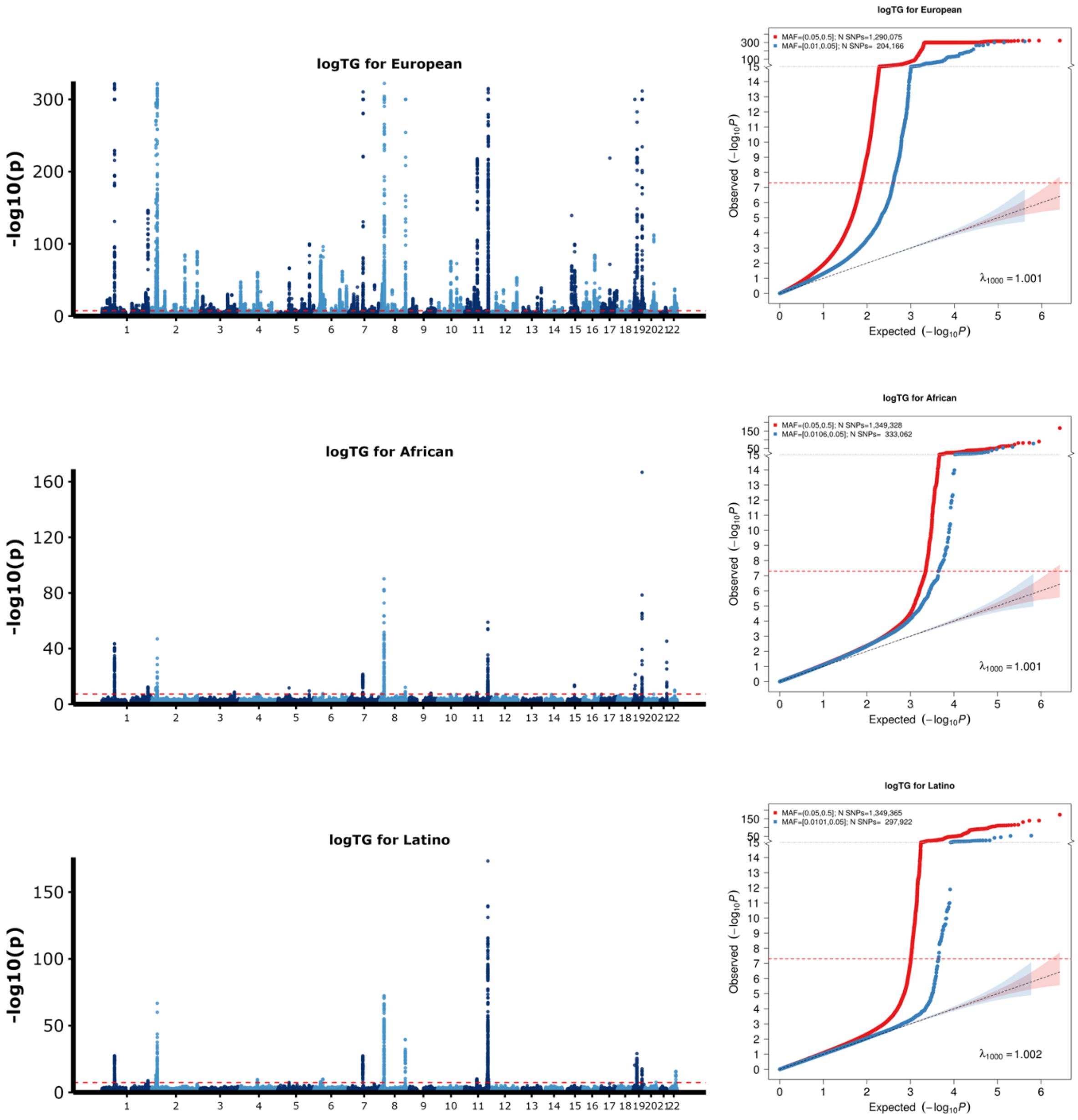


[1] For continuous traits, $\lambda_{1000}$ scales the genomic inflation factor $\lambda$ to a study with 1000 subjects using $\lambda_{1000} = 1 + 1000 * (\lambda - 1)/N$, where N is the total sample size. For binary traits, $\lambda_{1000}$ scales $\lambda$ to a study with 1000 cases and 1000 controls using $\lambda_{1000} = 1 + 1000 * (\lambda - 1) * (\frac{1}{N_{case}} + \frac{1}{N_{control}})$

**Supplementary Figure 18 continued: Manhattan and QQ plots for logTG in the East Asian and South Asian populations.**

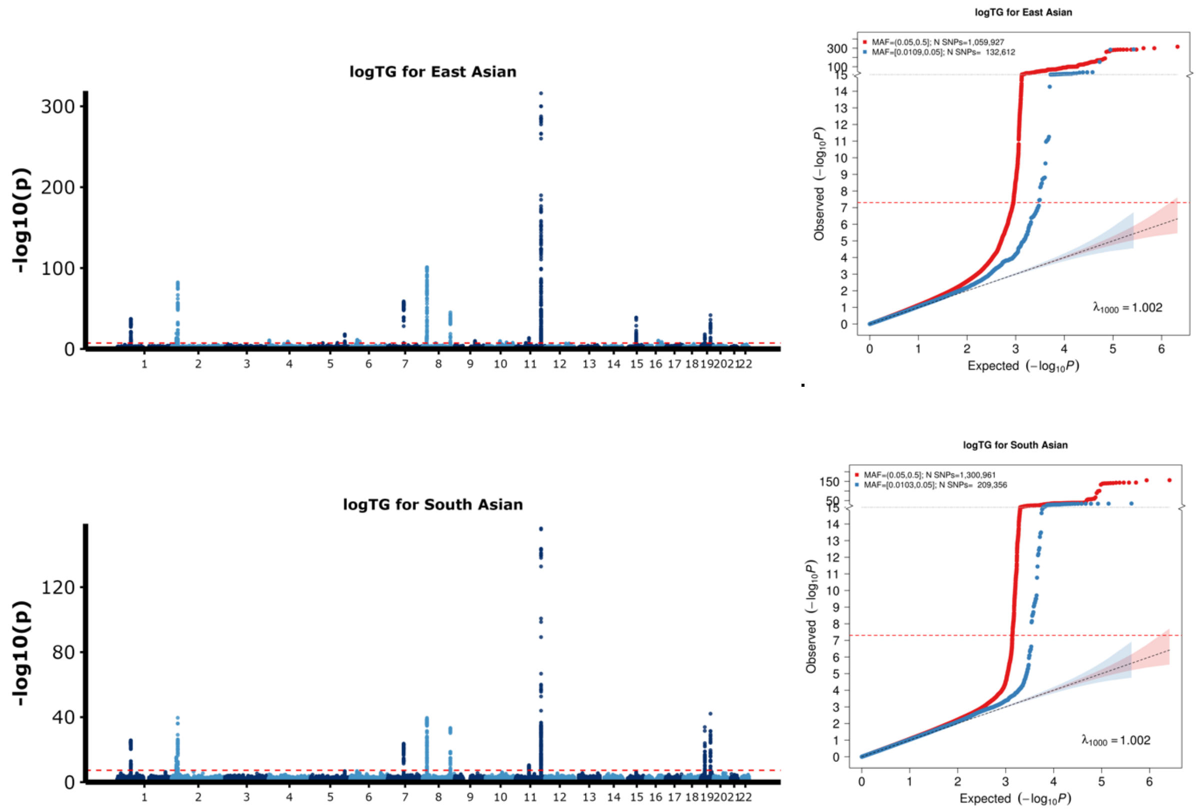


[1] For continuous traits, $\lambda_{1000}$ scales the genomic inflation factor $\lambda$ to a study with 1000 subjects using $\lambda_{1000} = 1 + 1000 * (\lambda - 1)/N$, where N is the total sample size. For binary traits, $\lambda_{1000}$ scales $\lambda$ to a study with 1000 cases and 1000 controls using $\lambda_{1000} = 1 + 1000 * (\lambda - 1) * (\frac{1}{N_{case}} + \frac{1}{N_{control}})$

**Supplementary Figure 19:** Manhattan plot and QQ plot[1] based on the Global Lipids Genetics Consortium (GLGC) GWAS summary statistics for **total cholesterol (TC)** in five populations: European, African (primarily African American), Latino, East Asian, South Asian (from top panel to bottom panel). The red and blue shaded regions around the diagonal line in the QQ plots indicate the 95% confidence intervals expected under the null hypothesis of no association between genetic markers and the trait of interest, for minor allele frequencies (MAF) within the ranges (0.05, 0.5] and [0.01, 0.05], respectively. Under the null hypothesis, the p-value follows a uniform (0,1) distribution. The jth order statistic follows a Beta (j, N-j+1) distribution, where N is the total number of variants given a specific MAF cutoff.

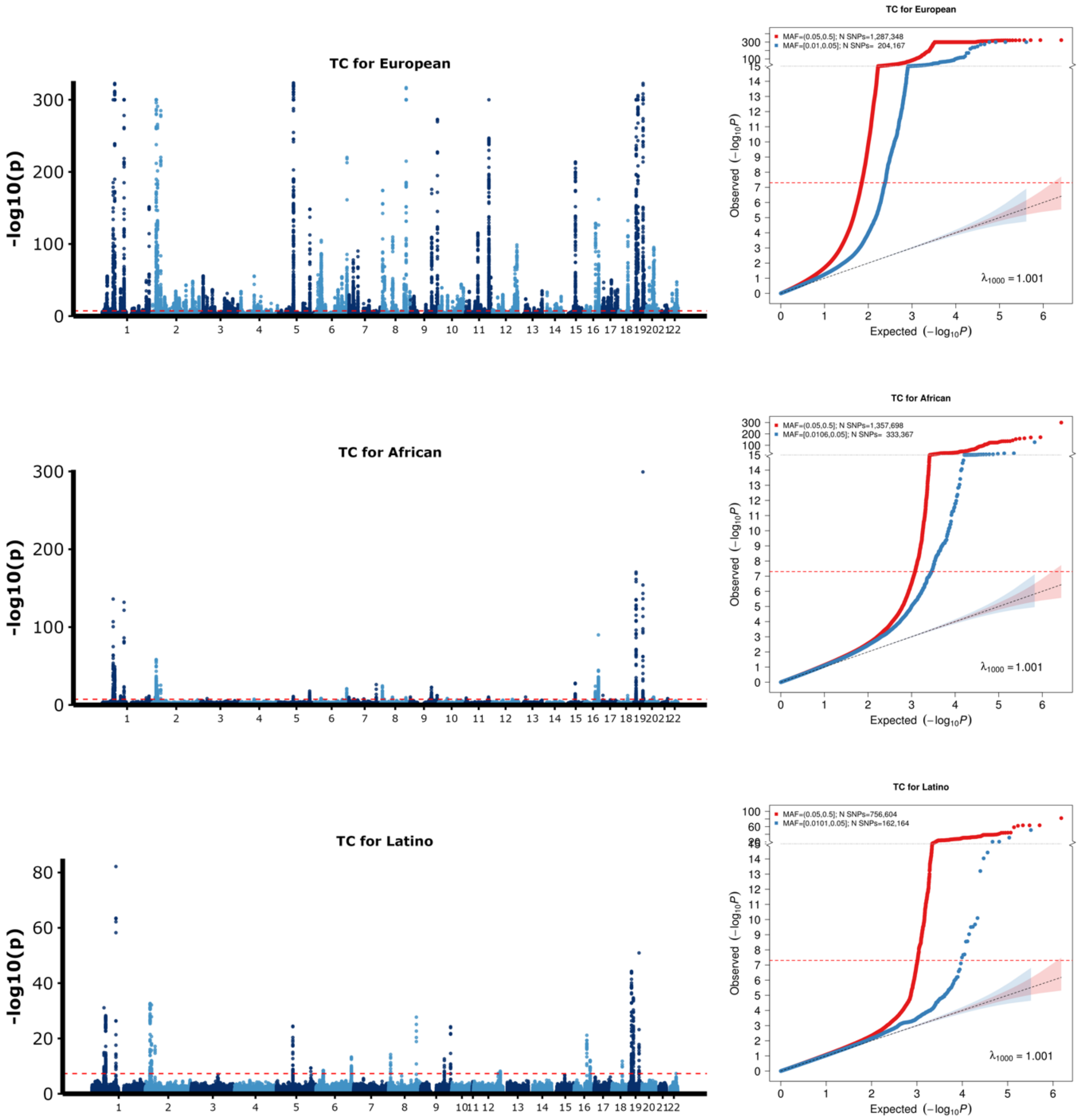


[1] For continuous traits, $\lambda_{1000}$ scales the genomic inflation factor $\lambda$ to a study with 1000 subjects using $\lambda_{1000} = 1 + 1000 * (\lambda - 1)/N$, where N is the total sample size. For binary traits, $\lambda_{1000}$ scales $\lambda$ to a study with 1000 cases and 1000 controls using $\lambda_{1000} = 1 + 1000 * (\lambda - 1) * (\frac{1}{N_{case}} + \frac{1}{N_{control}})$

**Supplementary Figure 19 continued: Manhattan and QQ plots for TC in the East Asian and South Asian populations.**

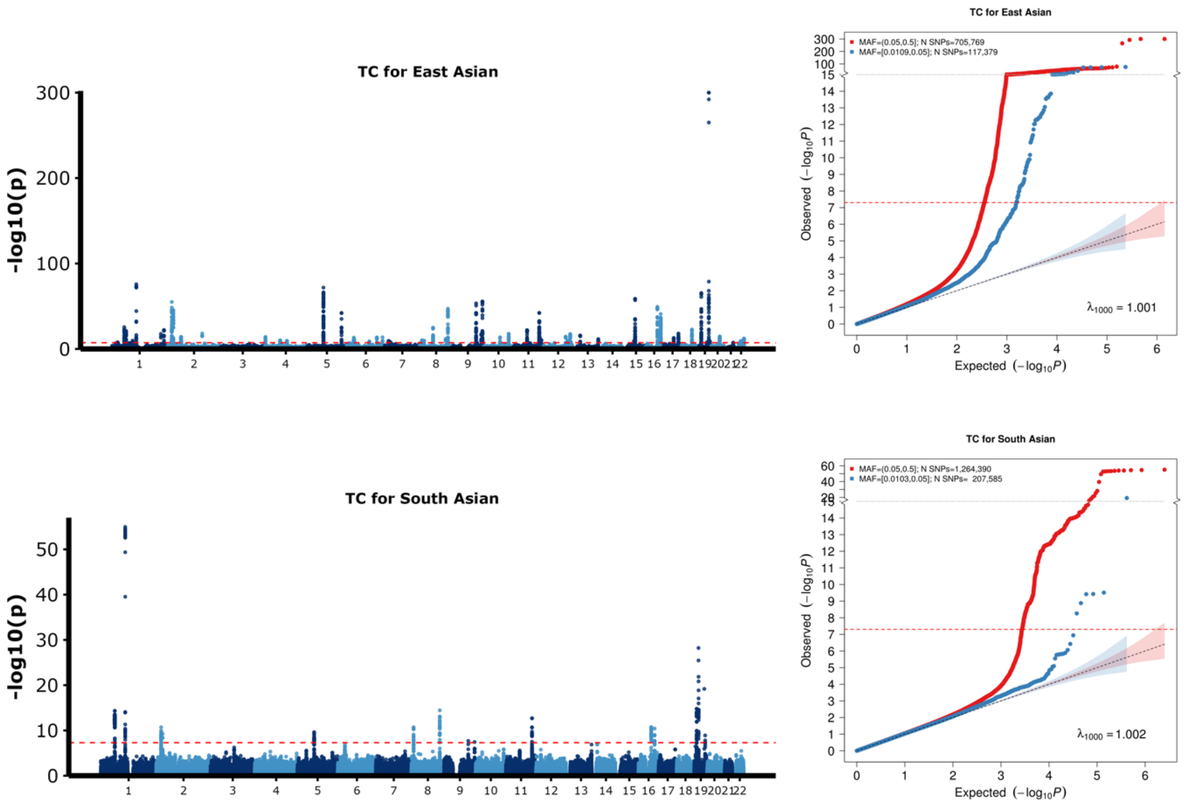


[1] For continuous traits, $\lambda_{1000}$ scales the genomic inflation factor $\lambda$ to a study with 1000 subjects using $\lambda_{1000} = 1 + 1000 * (\lambda - 1)/N$, where N is the total sample size. For binary traits, $\lambda_{1000}$ scales $\lambda$ to a study with 1000 cases and 1000 controls using $\lambda_{1000} = 1 + 1000 * (\lambda - 1) * (\frac{1}{N_{case}} + \frac{1}{N_{control}})$

**Supplementary Figure 20:** Manhattan plot and QQ plot[1] based on All of Us (AoU) GWAS summary statistics for **body mass index (BMI)** in three populations: European, African, Latino (from top panel to bottom panel). The red and blue shaded regions around the diagonal line in the QQ plots indicate the 95% confidence intervals expected under the null hypothesis of no association between genetic markers and the trait of interest, for minor allele frequencies (MAF) within the ranges (0.05, 0.5] and [0.01, 0.05], respectively. Under the null hypothesis, the p-value follows a uniform (0,1) distribution. The jth order statistic follows a Beta (j, N-j+1) distribution, where N is the total number of variants given a specific MAF cutoff.

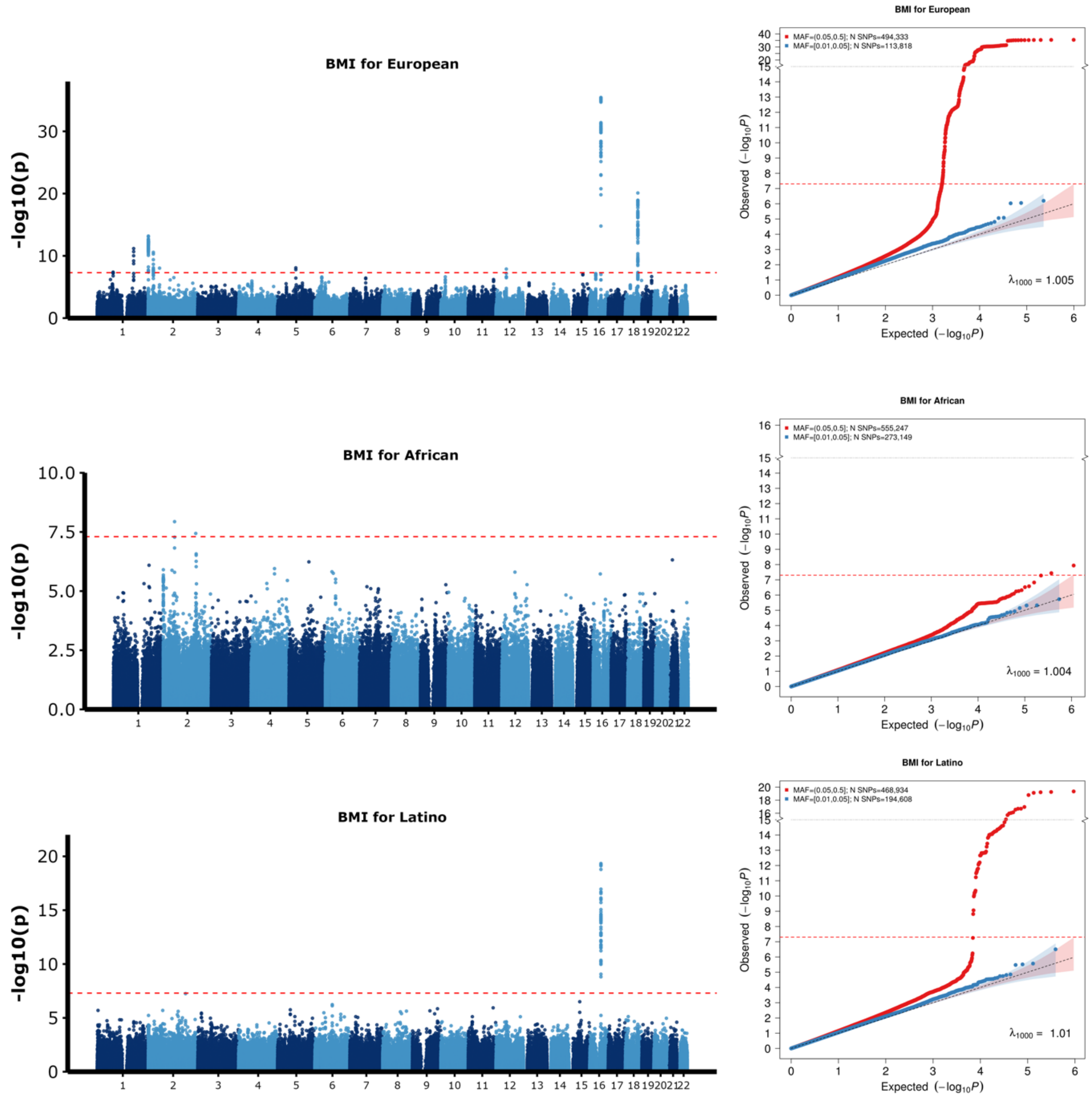


[1] For continuous traits, $\lambda_{1000}$ scales the genomic inflation factor $\lambda$ to a study with 1000 subjects using $\lambda_{1000} = 1 + 1000 * (\lambda - 1)/N$, where N is the total sample size. For binary traits, $\lambda_{1000}$ scales $\lambda$ to a study with 1000 cases and 1000 controls using $\lambda_{1000} = 1 + 1000 * (\lambda - 1) * (\frac{1}{N_{case}} + \frac{1}{N_{control}})$

**Supplementary Figure 21:** Manhattan plot and QQ plot[1] based on All of Us (AoU) GWAS summary statistics for **height** in three populations: European, African, Latino (from top panel to bottom panel). The red and blue shaded regions around the diagonal line in the QQ plots indicate the 95% confidence intervals expected under the null hypothesis of no association between genetic markers and the trait of interest, for minor allele frequencies (MAF) within the ranges (0.05, 0.5] and [0.01, 0.05], respectively. Under the null hypothesis, the p-value follows a uniform (0,1) distribution. The jth order statistic follows a Beta (j, N-j+1) distribution, where N is the total number of variants given a specific MAF cutoff.

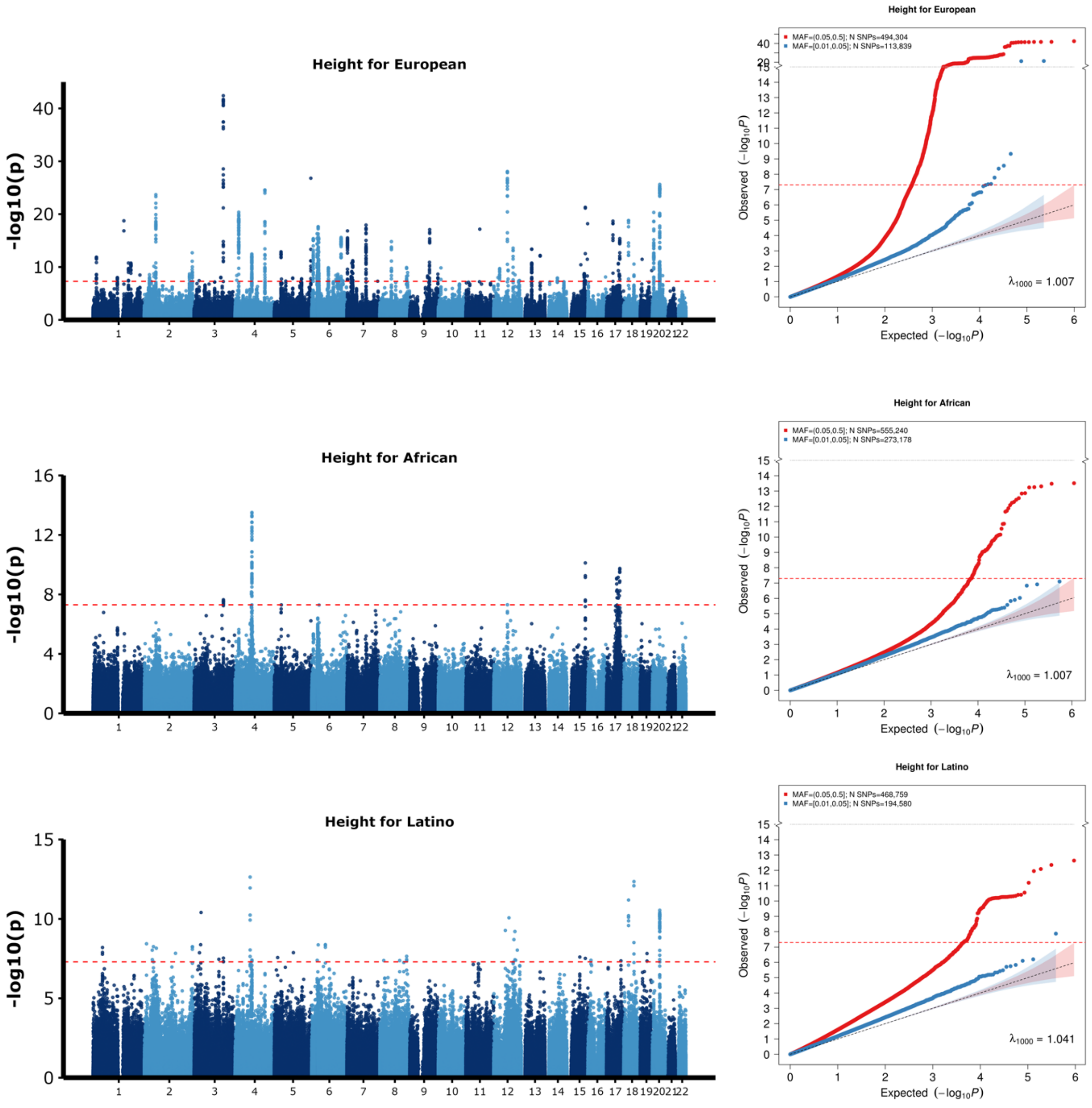


[1] For continuous traits, $\lambda_{1000}$ scales the genomic inflation factor $\lambda$ to a study with 1000 subjects using $\lambda_{1000} = 1 + 1000 * (\lambda - 1)/N$, where N is the total sample size. For binary traits, $\lambda_{1000}$ scales $\lambda$ to a study with 1000 cases and 1000 controls using $\lambda_{1000} = 1 + 1000 * (\lambda - 1) * (\frac{1}{N_{case}} + \frac{1}{N_{control}})$

**Supplementary Figure 22: Number of SNPs across five populations in 1000 Genomes Project (Phase 3).** SNPs are selected with MAF > 0.01 in at least one of five populations: African (AFR), American (AMR), East Asian (EAS), European (EUR) and South Asian (SAS).

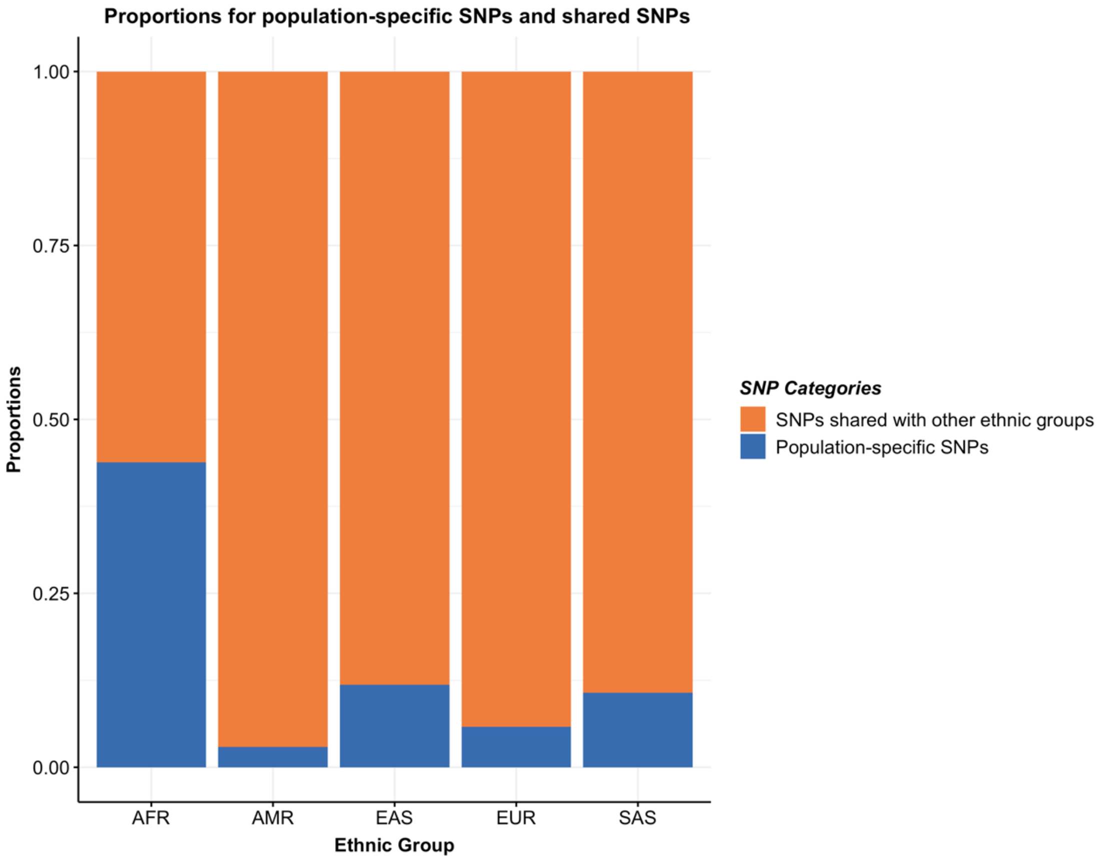

### Supplementary Note

#### Area under the ROC curve (AUC) to logit-scale variance

For continuous traits, the relative-$R^2$ is used to compare the performance of two different PRS methods. Relative-$R^2$ can be viewed as the ratio of the amount of outcome variance that two different PRS methods can explain. However, for binary traits, AUC is bounded between 0.5 to 1. Relative-AUC is not directly comparable to relative-R2 to measure the relative performance between two different methods. Therefore, we convert the AUC to logit-scale variance using the formula[1], $\sigma^2 = 2 * \phi(AUC)^{-1}$, where $\sigma^2$ is the amount of variance the PRS can explain on the logit-scale, $\phi$ is the cumulative distribution of the standard normal distribution. To compare the prediction performance of two different PRS for binary traits, we use relative logit scale variance for binary traits (**Supplementary Table 10-11**).

#### Super learning

Let $X = (PRS_1, PRS_2, \dots, PRS_M)$ be the input dataset. Suppose the predicted outcome are $\psi(X)_v$ for $v = 1,2, \dots V$ different prediction algorithms. For continuous traits, the objective function for each of the prediction algorithm is defined to minimize $\{Y - \psi(X)_v\}^2$. We propose a weighted combination of the $V$ different predictors as $F(X) = \sum_{v=1}^{V} \alpha_v \, \hat{\psi}(X)_v$ with $\sum_{v=1}^{V} \alpha_v = 1$ and $\alpha_v \geq 0$ for any $v$. Then the weights $\alpha$ are determined using the tuning dataset with:

$$\hat{\alpha} = \arg\min_{\alpha} \{Y - F(X)\}^2,$$

For binary traits, the objective function for each of the algorithm is to maximize the AUC. The weights $\alpha$ are determined using the tuning dataset by maximizing the AUC while using $F(X)$ to predict $Y$. With the optimal weights of the $V$ different algorithm, the super learner estimator is:

$$\hat{\psi}^{SL} = \sum_{v=1}^{V} \hat{\alpha}_v \, \hat{\psi}_v.$$

In practice, we use three different prediction algorithms implemented in the SuperLearner package[2] to generate the super learning estimate: Lasso[3], ridge regression[4] and neural networks[5]. The tuning parameters for these algorithms are put as the defaults as SuperLearner package. Other common prediction algorithms, such as random forest[6] and XGBoost[7], are also available in the R package, and can be added for constructing PRSs by the user. For binary traits, since the ridge regression algorithm is not supported by SuperLearner package now, we only use Lasso and neural networks in real data analysis. To use AUC as the objective function, we use the flag "method = method.AUC" in the SupearLearner package.

**CT-SLEB with more than two ancestries**

When more than two ancestries' summary statistics are provided, CT-SLEB is implemented as follows. To build the PRS, we first choose the target population. In the CT step, we only use the GWAS summary statistics from the target population and the European population. This step is implemented as such due to the potential computing burden by increasing the dimensions of CT step. After the CT step finishes, we use the summary statistics for all available ancestries for estimating the posterior effect-size for each SNP. By incorporating the other available GWAS summary statistics from non-European populations, the precision of the EB estimates is more accurate than only using the target population and European population. Suppose $L$ ancestries are available in the analyses, after the EB step, the posterior effect-size for the kth SNP is $\widehat{\boldsymbol{\beta}}_k^{EB} = (\hat{\beta}_{k1}^{EB}, \hat{\beta}_{k2}^{EB}, \dots, \hat{\beta}_{kl}^{EB}, \dots, \hat{\beta}_{kL}^{EB})$, where the $l$th element is the posterior effect-size for the $l$th ancestry. Let $l = 1$ represents the target population, $l = 2$ represents the European population, and $l = 3, \dots, L$ represents the other non-European populations. By cross-combing 6 $r^2$ thresholds * 2 $w_b$ windows * 81 value thresholds, we can compute 972 PRSs for each population using the posterior effect-size. In practice, we only compute 1944 PRSs using the posterior effect-size from the target population

($\sum G_k \hat{\beta}_{k1}^{EB}$) and the European population ($\sum G_k \hat{\beta}_{k2}^{EB}$). This explains why the computation time for five ancestries and two ancestries CT-SLEB is comparable. The EB step is very fast since we compute the posterior effect-size for one SNP at a time. The 972 PRSs based on other non-European populations are not included since the adding more PRSs can increase the computation time for PRSs calculations and significantly increase the time in the super learning step.

#### Mild and no negative selection simulation generation

Suppose $u_{kl}$ is the standardized effect size for $k$th causal SNP for the $l$th population. We consider generating the effect size of causal variants are related to allele frequency under mild negative selection ($u_{kl}^2 \propto [f_{kl}(1-f_{kl})]^{0.75}$) and no negative selection ($u_{kl}^2 \propto [f_{kl}(1-f_{kl})]$) scenarios. To generate the required effect-size, we first draw from a multivariate normal distribution:

$$v_{kl} \sim N\left(0, \frac{h^2}{C_l}\right), cov\left(v_{kl_1}, v_{kl_2}\right) = \frac{\rho h^2}{\sqrt{C_{l_1} C_{l_2}}},$$

where $v_{kl}$ is on a temporary generating scale, and $h^2$ is set to 0.4. Then, the temporary effect size is defined $u_{kl}^* = v_{kl}\{\sqrt{2 f_{kl}(1-f_{kl})}\}^{\alpha}$, where $\alpha = 0.75$ is for mild negative selection and $\alpha = 1$ was for no negative selection. To control the common SNPs heritability as 0.4, the standardized effect size is generated by scaling the temporary effect-size as $u_{kl} = u_{kl}^*\{\sqrt{\frac{0.4}{\sum_{l=1}^{C_l}(u_{kl}^*)^2}}\}$. The genetic correlation $\rho$ is defined on the generating scale. For both mild and no negative selection setting, $\rho$ is set to 0.8.

#### Genotyping of 23andMe data

DNA extraction and genotyping were performed on saliva samples by National Genetics Institute (NGI), a CLIA licensed clinical laboratory and a subsidiary of Laboratory Corporation of America. Samples were genotyped on one of five genotyping platforms.

The v1 and v2 platforms were variants of the Illumina HumanHap550+ BeadChip, including about 25,000 custom SNPs selected by 23andMe, with a total of about 560,000 SNPs. The v3 platform was based on the Illumina OmniExpress+ BeadChip, with custom content to improve the overlap with our v2 array, with a total of about 950,000 SNPs. The v4 platform was a fully customized array, including a lower redundancy subset of v2 and v3 SNPs with additional coverage of lower-frequency coding variation, and about 570,000 SNPs. The v5 platform, in current use, is an Illumina Infinium Global Screening Array (~640,000 SNPs) supplemented with ~50,000 SNPs of custom content. This array was specifically designed to better capture global genetic diversity and to help standardize the platform for genetic research. Samples that failed to reach 98.5% call rate were re-analyzed. Individuals whose analyses failed repeatedly were re-contacted by 23andMe customer service to provide additional samples.

**Ancestry determination of 23andMe data**

For our standard GWAS, we restrict participants to a set of individuals who have a specified ancestry determined through an analysis of local ancestry[8]. Briefly, our algorithm first partitions phased genomic data into short windows of about 300 SNPs. Within each window, we use a support vector machine (SVM) to classify individual haplotypes into one of 31 reference populations (https://www.23andme.com/ancestry-composition-guide/). The SVM classifications are then fed into a hidden Markov model (HMM) that accounts for switch errors and incorrect assignments, and gives probabilities for each reference population in each window. Finally, we use simulated admixed individuals to recalibrate the HMM probabilities so that the reported assignments are consistent with the simulated admixture proportions. The reference population data is derived from public datasets (the Human Genome Diversity Project, HapMap, and 1000 Genomes), as well as 23andMe customers who have reported having four grandparents from the same country.

Ancestries are defined as follow:

| **Ancestry** | **Classification criteria** |
|---|---|
| European | European + Middle Eastern > 0.97, European > 0.90 |
| East Asian | East Asian + Southeast Asian > 0.97 |
| South Asian | South Asian > 0.97 |
| Middle Eastern (& North African) | Middle Eastern + European > 0.97, Middle Eastern > 0.90 |
| African American + Latinos | European + African + East Asian + Native American + Middle Eastern > 0.90, African + Native American > 0.01 |

African Americans and Latinos are admixed with broadly varying contributions from Europe, Africa and the Americas. Therefore, no single threshold of genome-wide ancestry will be able to effectively discriminate African Americans and Latinos. However, the distributions of the length of segments of European, African and American ancestry are very different between African Americans and Latinos, because of distinct admixture timing between the three ancestral populations in the two ethnic groups. Therefore, we trained a logistic classifier that takes one customer's length histogram of segments of African, European and American ancestry, and predict whether the customer is likely African American or Latino.

**Selecting unrelated individuals within 23andMe data**

A maximal set of unrelated individuals was chosen for each analysis using a segmental identity-by-descent (IBD) estimation algorithm[9]. Individuals were defined as related if they shared more than 700 cM IBD, including regions where the two individuals share either one or both genomic segments IBD. This level of relatedness (roughly 20% of the genome) corresponds approximately to the minimal expected sharing between first cousins in an outbred population. When selecting individuals for case/control phenotype analyses, the selection process is designed to maximize case sample size by preferentially retaining cases over controls. Specifically, if both an individual case and an individual control are found to be related, then the case is retained in the analysis.

**Imputation of 23andMe data**

Imputation panels created by combining multiple smaller panels have been shown to give better imputation performance than the individual constituent panels alone[10]. To that end, we combined the May 2015 release of the 1000 Genomes Phase 3 haplotypes[11] with the UK10K imputation reference panel[12] to create a single unified imputation reference panel. To do this, multiallelic sites with N alternate alleles were split into N separate biallelic sites. We then removed any site whose minor allele appeared in only one sample. For each chromosome, we used Minimac3[13] to impute the reference panels against each other, reporting the best-guess genotype at each site. This gave us calls for all samples over a single unified set of variants. We then joined these together to get, for each chromosome, a single file with phased calls at every site for 6,285 samples. Throughout, we treated structural variants and small indels in the same way as SNPs.

In preparation for imputation we split each chromosome of the reference panel into chunks of no more than 300,000 variants, with overlaps of 10,000 variants on each side. We used a single batch of 10,000 individuals to estimate Minimac3 imputation model parameters for each chunk.

To generate phased participant data for the v1 to v4 platforms, we used an internally-developed tool, Finch, which implements the Beagle graph-based haplotype phasing algorithm[14], modified to separate the haplotype graph construction and phasing steps. Finch extends the Beagle model to accommodate genotyping error and recombination, in order to handle cases where there are no consistent paths through the haplotype graph for the individual being phased. We constructed haplotype graphs for all participants from a representative sample of genotyped individuals, and then performed out-of-sample phasing of all genotyped individuals against the appropriate graph. For the X chromosome, we built separate haplotype graphs for the non-pseudoautosomal region and each pseudoautosomal region, and these regions were phased separately. For the 23andMe participants genotyped on the v5 array, we used a similar approach, but using a new phasing algorithm, Eagle2[15]. We imputed phased participant data against the merged reference panel using Minimac3, treating males as homozygous pseudo-diploids for the non-pseudoautosomal region.

We compute association test results for the genotyped and the imputed SNPs. For case control phenotypes, we compute association by logistic regression assuming additive allelic effects. For tests using imputed data, we use the imputed dosages rather than best-guess genotypes. As standard, we include covariates for age, gender, the top five principal components to account for residual population structure, and indicators for genotype platforms to account for genotype batch effects. The association test *P* value we report is computed using a likelihood ratio test, which in our experience is better behaved than a Wald test on the regression coefficient. For quantitative traits, association tests are performed by linear regression. Results for the X chromosome are computed similarly, with male genotypes coded as if they were homozygous diploid for the observed allele.

**Principal component (PC) calculation using 23andMe data**

A principal component analysis was performed independently for each ancestry, using ~65,000 high quality genotyped variants present in all five genotyping platforms. It was computed on a subset of participants randomly sampled across all the genotyping platforms (137K, 102K, 1000K, 360K and 32K participants were used for African-American, East-Asian, European, Latino, and South-Asian, respectively). PC scores for participants not included in the analysis were obtained by projection, combining the eigenvectors of the analysis and the SNP weights.

**Quality control of 23andMe data**

The vast majority of SNPs are only imputed and not genotyped, and therefore they only have imputed GWAS results. A small proportion of SNPs (often rare) have only genotyped GWAS results. Finally, the majority of genotyped SNPs are also imputed. When choosing between imputed and genotyped GWAS results for these SNPs, if they both passed quality control (QC), we report the imputed result.

Variant QC is applied independently to genotyped and imputed GWAS results, and we flag the SNPs failing QC. For QC of genotyped GWAS results, we flagged SNPs that were only genotyped on our "v1" and/or "v2" platforms due to small sample size, and SNPs on chrM or chrY because many of these are not currently called reliably. Using trio data, we flagged SNPs that failed a test for parent-offspring transmission; specifically, we regressed the child's allele count against the mean parental allele count and flagged SNPs with fitted $\beta<0.6$ and $P<10^{-20}$ for a test of $\beta<1$. We flagged SNPs with a Hardy-Weinberg $P<10^{-20}$, or a call rate of <90%. We also tested genotyped SNPs for genotype date effects, and flagged SNPs with $P<10^{-50}$ by ANOVA of SNP genotypes against a factor dividing genotyping date into 20 roughly equal-sized buckets. We flagged SNPs with large sex effect (ANOVA of SNP genotypes, $r^2>0.1$). Finally, we flag SNPs with probes matching multiple genomic positions in the reference genome ('self chain').

For imputed GWAS results, we flagged SNPs with rsq<0.3, as well as SNPs that had strong evidence of a platform batch effect. The batch effect test is an F test from an ANOVA of the SNP dosages against a factor representing v4 or v5 platform; we flagged results with $P<10^{-50}$. Prior to GWAS, we identified, for each SNP, the largest subset of the data passing these criteria, based on their original genotyping platform -- either v2+v3+v4+v5, v4+v5, v4, or v5 only -- and computed association test results for whatever was the largest passing set. As a result, there are no imputed results for SNPs that fail these filters.

Across all results, we flag SNPs that have an available sample size of less than 20% of the total GWAS sample size. We also flag logistic regression results that did not converge due to complete separation, identified by abs (effect) >10 or stderr>10 on the log odds scale. We also flag linear regression results for SNPs with MAF < 0.1% because tests of low frequency variants can be sensitive to violations of the regression assumption of normally distributed residuals.

**Quality control of the GWAS summary statistics for 23andMe, Global Lipids Genetics Consortium (GLGC), and All of Us (AoU) data**

Before training the PRS models, the GWAS summary statistics for 23andMe, GLGC and AoU are cleaned using the following steps:

1. Keep the SNPs that are available on HapMap3 (HM3) + Multi-Ethnic Genotyping Arrays (MEGA) arrays.
2. Remove SNPs with duplicated positions.
3. Keep only common SNPs for each ancestry (ancestry specific MAF > 0.01). For GLGC data, due to the lack of ancestry-specific MAF, so we used the ancestry-

specific MAF from the 1000 Genomes Project Phase 3 (1KG) data for this step of filtering.

4. Remove SNPs with alleles are “AT”, “TA”, “CG”, “GC” to avoid undetectable flipping strands when matching with UK Biobank (UKBB) data in the tuning and validation step. This step is not necessary for 23andMe data because we also use 23andMe individuals for tuning and validation.
5. Remove SNPs from the analysis whose sample size is less than 90% of the total sample size.
6. Keep the SNPs that are present in the 1KG, which is used as the reference data for LD estimation.

**Genetic predicted ancestry in UKBB dataset**

We compute genetic ancestry for all individuals in UKBB who are not self-reported Whites. To balance the samples from all ancestries, we also include 8,000 unrelated Whites to form the set of UKBB individuals in this analysis. We use individuals from 1KG as the reference data for ancestry prediction. We initially compute 20 genetic PCs for all UKBB and 1KG individuals together by--pca 20 allele-wts in PLINK2.0[16]. The PCs are calculated based on a set of variants from gnomAD[17] (https://gnomad.broadinstitute.org/help/how-should-i-cite-discoveries-made-using-gnomad-data) that can be used to capture enough ancestral information. After obtaining the top 20 PCs, we train a random forest classifier with 1500 trees using PCs of 1KG individuals with their true ancestral labels. To train the random forest classifier, we use the R package randomForest version 4.6-14[18]. Finally, we apply this random forest classifiers to the PCs of UKBB individuals and get their genetic ancestry prediction.
